# A heterogeneous pharmaco-transcriptomic landscape induced by targeting a single oncogenic kinase

**DOI:** 10.1101/2024.04.08.587960

**Authors:** Ross M. Giglio, Nicholas Hou, Lingting Shi, Adeya Wyatt, Justin Hong, Mathini Vaikunthan, Charlotte Bellamy, Henry Fuchs, Vivian Lu, Ndubuisi Obasi, Jose Pomarino Nima, Seth W. Malinowski, Keith L. Ligon, J. Ricardo McFaline-Figueroa, Nir Yosef, Elham Azizi, José L. McFaline-Figueroa

## Abstract

Over-activation of the epidermal growth factor receptor (EGFR) is a hallmark of glioblastoma. However, EGFR-targeted therapies have led to minimal clinical response. While delivery of EGFR inhibitors (EGFRis) to the brain constitutes a major challenge, how additional drug-specific features alter efficacy remains poorly understood. We introduce SCHEMATIC, which integrates multiplex single-cell chemical transcriptomics with deep-generative classification to resolve chemotype-specific and shared programs and apply it to to define the molecular response of glioblastoma to EGFRis. We identify programs that differ by the chemical properties of EGFRis, including induction of adaptive transcription and modulation of immunogenic gene expression. We find that induction of an adaptive transcriptional program is associated with persistence of surviving cells after EGFR inhibition, and that concurrent EGFR/PI3K inhibition attenuates this program. We also find that pro-immunogenic expression changes associated with a subset of tyrphostin-family EGFR inhibitors are accompanied by enhanced antigen-specific cytotoxic T-cell killing *in vitro*. Our study provides a framework that considers each agent’s unique and often unknown poly-pharmacology to prioritize compounds pre-clinically that induce favorable molecular responses.

---

The response of cancer to therapy is driven by a complex interplay between a tumor’s genetic background and the mode of anti-tumor therapy. The development of bulk shRNA and CRISPR-based genome-wide screens has drastically increased our ability to identify the genetic requirements of cancer cell response to therapy. These screens use gross phenotypic assays (e.g., viability), specific molecular measurements (reporter activity), or, more recently, single-cell profiling as readouts^1–4^. Variability across the drug’s mechanisms of action and polypharmacology can also result in substantial heterogeneity in a tumor’s response to therapy. Additionally, the classification of a drug by its predicted or previously annotated target has not always aligned with functional outcomes^5,6^, and in some cases, has led to inefficacy or unwanted off-target toxicity in clinical settings^7,8^. Altogether, we lack a comprehensive understanding of how different approaches to targeting oncogenic activity lead to heterogeneity in the molecular response to therapy.

A prime example of variable response to therapy is observed in IDH-wild type glioblastoma (GBM) - the most common and aggressive primary brain cancer. Despite a multimodal treatment regimen composed of surgical resection, radiotherapy, chemotherapy, and tumor-treating fields, most patients succumb to their disease within 2 years of diagnosis^9,10^. Large-scale efforts have defined the genomic landscape of GBM. These studies identified near-ubiquitous overactivation of receptor tyrosine kinase (RTK) signaling, loss of the negative regulation of the RAS/MAPK and PI3K effector pathways, and inactivation of the Rb and p53 tumor suppressor pathways as core alterations in oncogenic signaling^11,12^. Unfortunately, this genetic characterization of the disease has yet to translate into a therapeutic gain despite numerous clinical evaluations of targeted therapies in GBM.

Amplification and over-activating mutations in the epidermal growth factor receptor (EGFR) tyrosine kinase are found in approximately half of all GBM patients^11^, with the most prevalent mutation being truncation of exons 2-7 of EGFRs extracellular domain (EGFRvIII) which render the kinase constitutively active and with extrachromosomal amplification identified as a common mechanism by which glioblastoma cells acquire high levels of EGFR^13^. Although targeting of EGFR has demonstrated clinical efficacy in EGFR-mutated non-small cell lung cancer, no such benefit has been identified in GBM despite strong selection for EGFR activity in the disease. This inefficacy is consequent to a myriad of factors, including poor brain penetrance of small molecule inhibitors across the blood-brain barrier, which may be partially overcome thanks to recent advances in drug delivery (blood-brain-barrier opening, direct CSF infusion)^14–16^.

Case studies detailing the clinical response with a brain-penetrant EGFR inhibitor demonstrate that some GBM lesions have complete response to EGFR targeting, leading to renewed interest in the use of EGFR inhibitors in GBM^17,18^. More recently, a retrospective study found that a subset of patients treated with a combination of an EGFR inhibitor (osimertinib) and an anti-angiogenic (bevacizumab) experienced a long-term benefit^19^. Resistance to treatment was associated with secondary mutations in RTK-associated pathway components (*MET*, *IGF1R*, *PTEN*, *PDGFR*). Preclinical studies have also shown that cells eliminate EGFR containing extrachromosomal DNA (ecDNA) and suppress ecDNA EGFR expression in response to EGFR inhibition^20^. These resistance mechanisms highlight the central role of EGFR and RTK signaling in the disease, the importance of understanding how GBM responds to the loss of EGFR activity, and the need to identify therapies that block RTK activity while minimizing the emergence of resistance. However, few studies have focused on the mechanisms of innate and adaptive resistance of GBM to small molecules even for those with adequate distribution to the brain.

Here, we apply ultra-high throughput single-cell chemical transcriptomic profiling to map variability in tumor cell response to targeting of a single oncogenic event in one tumor type. We focus on the single-cell molecular profiling of patient-derived neurosphere GBM models (patient-derived cell lines or PDCLs) treated with 72 compounds and biomolecules with anti-EGFR activity and controls across ∼1,800 unique conditions (PDCL x EGFRi x concentration x timepoint). We first examine the changes in gene expression networks associated with RTK signaling, tumor cell proliferation, and adaptive transcriptional resistance to identify EGFR inhibitors that decrease viability with minimal induction of a resistance program in surviving cells. We then use Multi-resolution Variational Inference (MrVI)^21^, a hierarchical probabilistic model that learns single cell-specific sample stratifications, to identify drug-dose combinations with similar effects on distinct molecular programs. We leverage the information on how each compound alters these programs in each genetic background to arrive at a molecular classification of small molecule inhibitors. We term this integrated workflow **SCHEMATIC** (**S**ingle-cell **CHEM**ic**A**l **T**ranscriptom**I**cs-based chemotype **C**lassification), which couples multiplex single-cell chemical transcriptomics with a deep-generative classifier, to resolve chemotype-specific and shared molecular response programs. In our classification of EGFR-targeting chemotypes by induced molecular programs, we focus on expression changes of clinical interest and prioritize strategies to ameliorate adaptive resistance and compounds that promote pro-immunogenic responses. Overall, we present SCHEMATIC as a chemical genomics framework well positioned to annotate favorable or unfavorable molecular phenotypes, allowing for preclinical prioritization of compounds.

## Results

### Single-cell chemical transcriptomics identifies shared and distinct expression changes driven by a drug’s polypharmacology

Transcriptomic profiling of chemically exposed cells can identify commonalities in molecular response across compounds with similar mechanisms of action^5^. Their recent application at single-cell resolution allowed for a detailed description of how chemical perturbation alters cellular states and provided insight into the mechanisms of action not apparent across population-averaged data^22^. However, studies have yet to examine whether this approach can detect subtle differences in response within a compound class that inform on a drug’s off-target effects and polypharmacology.

To determine whether single-cell chemical transcriptomics profiling can identify differences in response across compounds targeting the same molecular activity, we devised a pilot screen using six small molecules that inhibit EGFR activity (afatinib, brigatinib, CUDC-101, EAI045, neratinib, osimertinib). Four of the six inhibitors target EGFR irreversibly (afatinib, EAI045, neratinib, osimertinib), one binds an allosteric site (EAI045), and two have been designed with activity against another enzyme (brigatinib^23^: an ALK inhibitor with anti-EGFR activity; CUDC-101^24^: a bifunctional EGFR & HDAC class I/II inhibitor). We exposed 3 established glioblastoma cell lines (A172, T98G, U87MG) and 1 PDCL (BT333) to 7 concentrations (0.005 - 10 µM) of each compound or DMSO vehicle control for 24 hours. To determine the effect of each compound on transcription, we processed conditions across two replicate experiments via sci-Plex multiplexing and subjected cells to single-cell combinatorial indexing RNA-seq^25^ (**Fig. 1A**). After filtering, we collected 188,746 single-cell transcriptomes across our 4 GBM models and 43 exposures for a total of 172 unique conditions (**Supp. Fig. 1A-B**).

**Figure 1.**
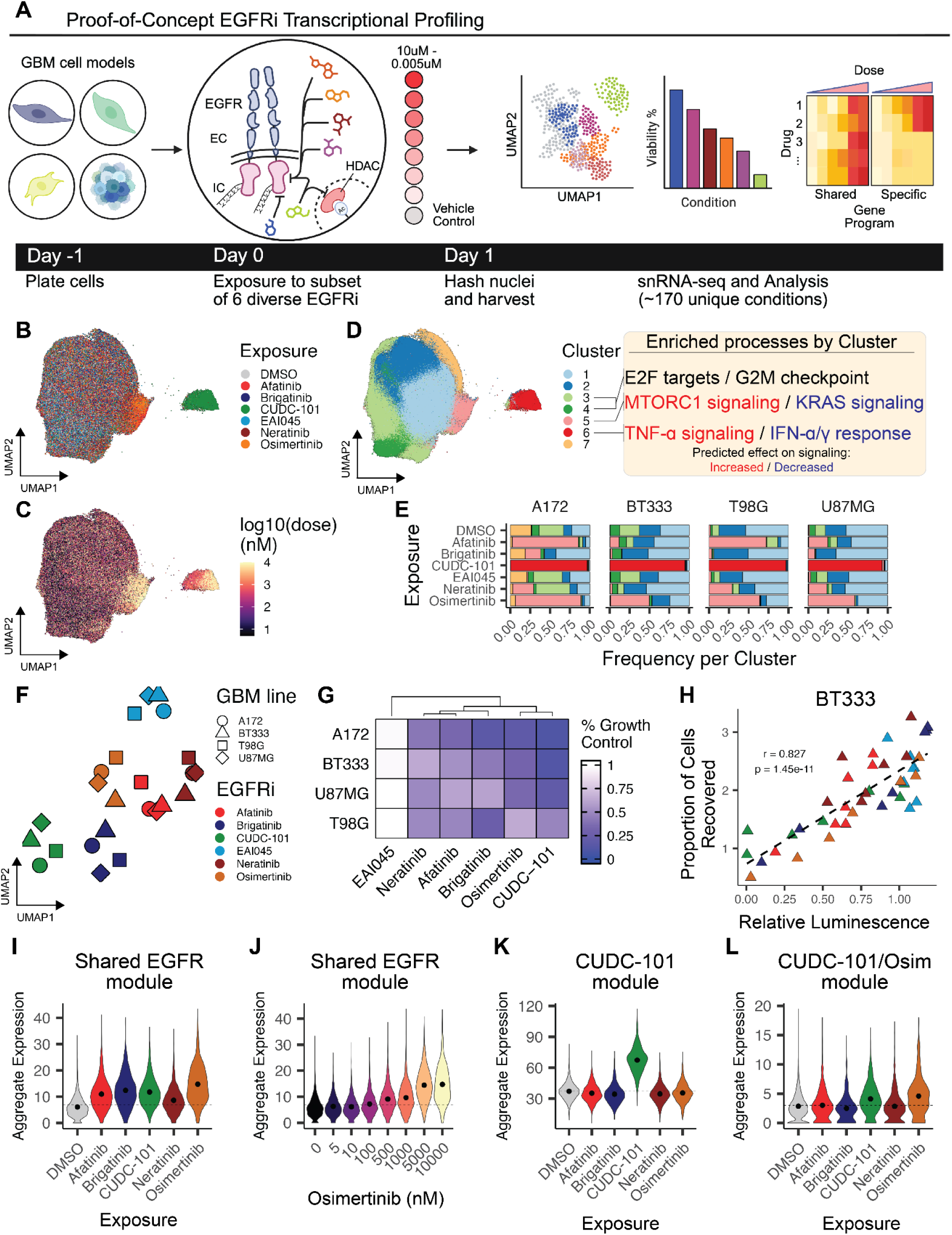
Single-cell chemical transcriptomics recovers drug-induced expression signatures that vary due to a compound’s polypharmacology. **A.** Schematic depicting a pilot screen of EGFR inhibitors in models of GBM. **B-C**. Integrated UMAPs of A172, T98G, U87MG, and BT333 GBM models exposed to one of 6 EGFR inhibitors or DMSO control. Color denotes the EGFR inhibitor (**B**) or the concentration of inhibitor (**C**) that a cell was exposed to. **D**. Left: Integrated UMAP as in (**B-C**) colored by the cluster assignment from Leiden-based community detection of in sample-integrated aligned PCA space. Right: Results of gene set enrichment analysis for MSigDB Hallmarks gene sets of biological processes that are significantly enriched (FDR < 5%, hypergeometric test) amongst differentially expressed genes (FDR < 0.1%, see Methods) for clusters associated with proliferation (clusters 3 & 4) and those that are dependent on EGFR inhibitor exposure (clusters 5 and 6). **E**. Distribution of GBM cells across clusters from **D**. **F**. UMAP projections summarizing the correlation (Pearson’s coefficient of effects size estimates across the union of differentially expressed genes) between cell line-drug effects on transcription, annotated by cell line and exposure. **G.** Percent growth control derived from CellTiter-Glo dose-response viability curves. Colors denote inhibitors as in (**F**). **H.** Proportion of cells recovered (% of total) of a given cell line, drug, dose within the sci-Plex screen versus its corresponding CellTiter-Glo luminescence relative to control. **I-J.** Violin plots of the aggregate expression of a shared EGFR inhibitor transcriptional module across BT333 cells exposed to a 10 µM dose of the specified EGFRi or DMSO control (**I**) or increasing concentrations of osimertinib (**J**). Dashed line denotes the median aggregate expression of DMSO control. **K-L**. Violin plots of the aggregate expression of a CUDC-101 specific (**K**) or shared CUDC-101/osimertinib (**L**) transcriptional module across BT333 cells exposed to a 10 µM dose of the specified EGFRi or DMSO control. Dashed line denotes the median aggregate expression of DMSO control.

Differential expression (DE) analysis identified robust changes in gene expression in response to exposure to 5 of the 6 EGFR inhibitors as a function of concentration (median of 733 DE genes per EGFRi, FDR < 0.1%, normalized beta > 0.05, **Supp. Fig. 1B-C**). Within our screen, we observed a general lack of response of tumor cells to EAI045. EAI045 (EGFR Allosteric Inhibitor 045) is a fourth generation inhibitor that selectively inhibits EGFR mutants that do not require receptor dimerization for their autophosphorylation with ∼1000-fold selectivity over wild-type EGFR^26^. The lack of a molecular response to EAI045 is consistent with the fact that none of the included GBM models harbor catalytic EGFR alterations. We then used integration^27^, dimensionality reduction^25,28^, and Leiden community detection^29^ to visualize the similarities and differences in response across our models, which varied as a function of the precise inhibitor used (**Fig. 1B-D)**. For example, exposure to high concentrations of either afatinib, brigatinib, neratinib or osimertinib led to the induction of a cell state associated with increased MTOR and decreased KRAS signaling (cluster 7, **Fig. 1D**). Conversely, cells exposed to high concentrations of the dual EGFR/HDAC inhibitor CUDC-101 gave rise to a distinct cell state associated with changes in TNF-ɑ signaling and an IFN response (cluster 6, **Fig. 1D**). To further explore whether the response to each inhibitor was similar across our GBM models, we performed a joint analysis to summarize the covariance of gene-level effects across the union of dose-dependent DE genes, which we previously showed can recover commonalities in drug-induced expression programs^22^ (**Fig. 1F, Supp. Fig. 1D-F**). This approach also suggests that the transcriptional response to a compound is similar across our models.

Next we explored the relationship between the molecular responses and orthogonal viability measurements. For each drug and GBM model, we measured viability using CellTiter-Glo (Promega) across 8 doses spanning 4 orders of magnitude. We used the area under the curve (AUC) of our viability concentration-response and normalized to vehicle control to calculate a percent growth control (**Supp. Fig. 2A-D**). Viability responses varied both by inhibitor and the GBM line (**Fig. 1G)** and was directly correlated to sci-Plex metrics like cell recovery **(Fig. 1H)** as shown previously^22^, and the effect on MKI67 expression **(Supp. Fig. 1G)**.

We next used hierarchical clustering to group modules of dose-dependent DE genes that covary across our experiment, identifying 17 and 15 modules of genes (each composed of > 30 genes) that are similarly and uniquely upregulated or downregulated, respectively, by one or more EGFRis (**Fig 1I-L**, **Supp. Fig. 3A-C**, **Supp. Table S1**). Amongst these gene modules, we identified a 48 gene signature that was upregulated in a dose-dependent manner broadly across all EGFR inhibitors, leading to a measurable effect on transcription (shared EGFR module, **Fig. 1I-K**). Interestingly, this signature had a significant and negative correlation with overall survival within the IDHwt GBM cohort^11^ from The Cancer Genome Atlas (TCGA PanCancer)^30^ (**Supp. Fig. 3D**). Given that the cohort is unlikely to contain patients treated with EGFR targeted therapy, this gene module may report on a general feature of populations of cells that survive cytotoxic or genotoxic therapy. Consistent with this hypothesis, this gene module correlates with increasing tumor grade and inversely correlates with survival across all gliomas (**Supp.** Fig. 3D-F).

In addition to identifying broadly shared programs, our analysis identified transcriptional changes specific to one or a subset of compounds. We identified modules of genes unique to CUDC-101 exposure, which likely report on transcriptional effects downstream of HDAC class I/II inhibition (**Fig. 1K**, **Supp.** Fig. 3A-C). We also identified a shared 70 gene signature between CUDC-101 and osimertinib-exposed cells (**Fig. 1L, Supp.** Fig. 2A-C). Interestingly, the EGFR targeting quinazoline functional group of CUDC-101^24^ is most structurally similar to that of afatinib^31^, not osimertinib, among the 6 compounds tested. Moreover, this module is weakly correlated with proliferation index (**Supp.** Fig. 3G-H), suggesting this shared program is not simply a consequence of loss of viability. Overall, and towards our goal of classifying compounds by their polypharmacology, our results show that a single-cell chemical transcriptomics approach captures differences between compounds targeting the same oncogenic alteration, including unexpected associations not readily explained by a drug’s molecular structure.

### Natively heterogeneous patient-derived GBM models vary in the expression of key RTK pathway components and evolution of cell states in response to EGFR inhibition

To comprehensively define pharmacological variability in the transcriptional response to EGFR inhibition (**Fig. 2A**), we profiled three PDCL cultures. In addition to being amenable to screening a large number of exposure conditions, these models maintain the genetic and molecular diversity observed in GBM tumors^32–34^. The three models chosen are representative of the diversity of EGFR status in the disease. All lines display amplification of the receptor, and two of three carry extracellular mutations that render EGFR constitutively active (**Fig. 2B**). In addition, the tumor suppressor landscape of these models is also representative of GBM, including mutations in negative regulators of the RTK pathway (*NF1*, *PTEN*) and the p53 and Rb pathways (*TP53*, *CDKN2A*, *CDKN2B*) (**Fig. 2B**).

**Figure 2.**
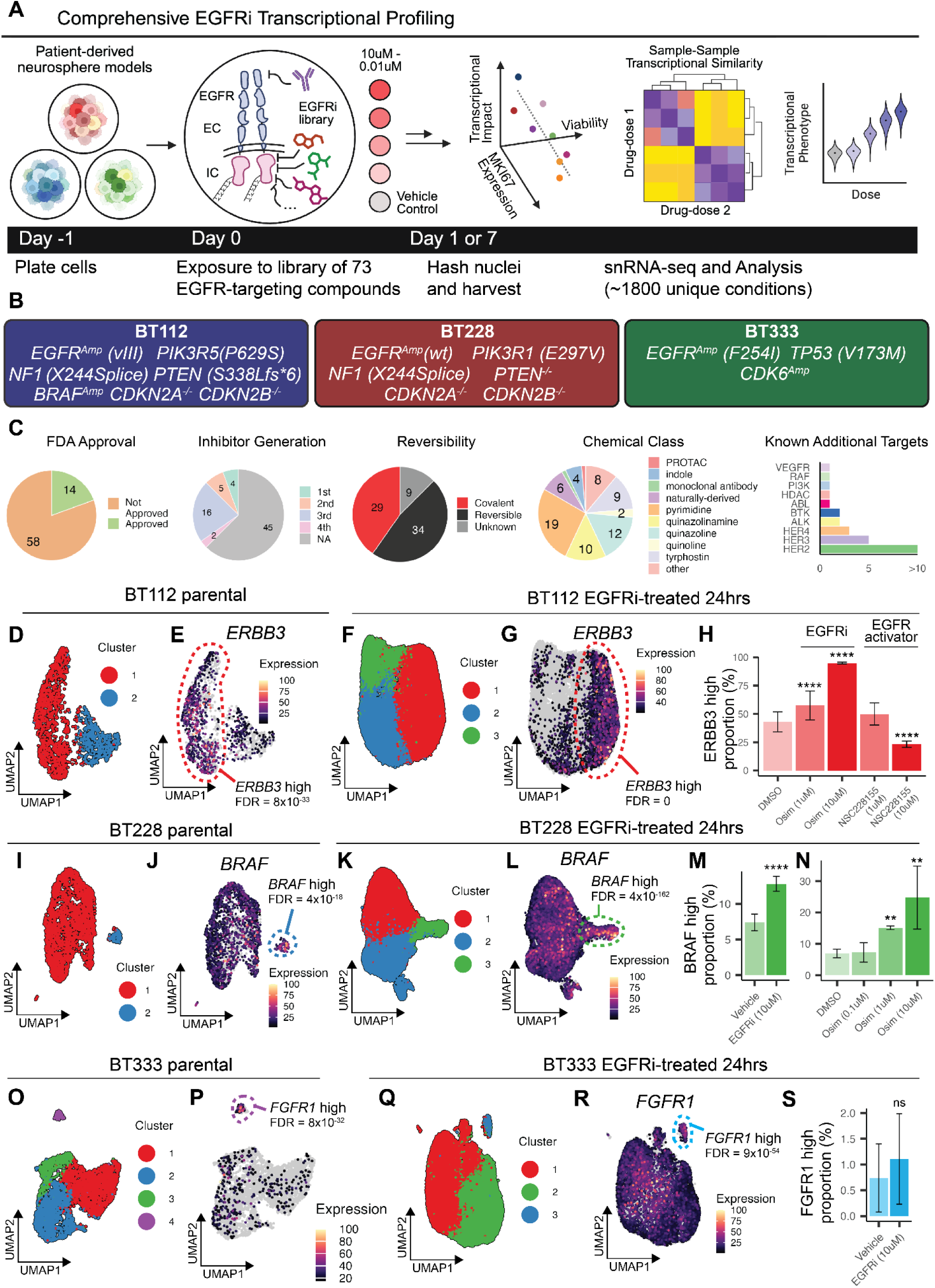
Comprehensive screening of patient-derived neurosphere GBM models reveals intra-model variation in RTK pathway expression and selection upon EGFR inhibition. **A**. Schematic depicting a comprehensive screen of EGFR inhibitors in models of GBM. **B**. Genetic status of EGFR and key GBM-associated genes in the patient-derived neurosphere models BT112, BT228, and BT333. **C**. Characteristics of the panel of 72 anti-EGFR compounds and biomolecules in our screen. **D-G**. UMAP embeddings of BT112 untreated parental cells (**D-E**) and 24 hour EGFRi library exposed cells (**F-G**), colored by leiden cluster (**D,F**) or *ERBB3* expression (as percent of maximum expression) (**E,G**). **H.** For a given treatment of BT112 cells, percent of cells present in *ERBB3* high-expressing cluster. **I-L**. UMAP embeddings of BT228 untreated parental cells (**I-J**) and 24 hour EGFRi library exposed cells (**K-L**), colored by leiden cluster (**I,K**) or *BRAF* expression (as percent of maximum expression) (**J,L**). **M.** For BT228 cells exposed to vehicle or any EGFRi in the library, percent of cells present in *BRAF* high-expressing cluster. **N**. As in (**M**), but for specified concentrations of osimertinib. **O-R**. UMAP embeddings of BT112 untreated parental cells (**O-P**) and 24 hour EGFRi library exposed cells (**Q-R**), colored by leiden cluster (**O,Q**) or *FGFR1* expression (as percent of maximum expression) (**P,R**). **S.** or BT333 cells exposed to vehicle or any EGFRi in the library, percent of cells present in *FGFR1* high-expressing cluster. Dotted circles on UMAPs denote subpopulations for BT112, BT228, and BT333 with a significant difference in expression of the specified gene (quasi-Poisson regression, Wald test, FDR < 1%). Error bars on barplots denote the standard deviation from the mean across replicates, and significant differences in percent of cells occupying cluster between a condition and its respective control (vehicle/DMSO) are denoted (Fisher’s exact test, **FDR<0.01, ****FDR<0.0001)

Multiplex single-cell RNA-seq profiling of basal (untreated) PDCL cultures identified substantial heterogeneity in the expression of genes associated with EGFR and RTK signaling. For example, while all three cell lines expressed EGFR, the expression of EGFR dimerization partners varied, with *ERBB2*, *ERBB3*, and *MET* highest in BT112 cells (**Supp. Fig. 4A**). The expression of other RTKs and RTK pathway components were similarly variable across models, including the expression of FGFR and RAF family members (**Supp. Fig. 4A**). We also identified heterogeneity in the expression of genes associated with TP53, RB, and CDK signaling (**Supp. Fig. 4B-C**).

Our analysis identified subpopulations within the three PDCL models profiled (**Fig. 2D, 2I, 2O**), which could not be explained solely by differences in proliferation or transcripts captured per cell (**Supp. Fig. 5**). Examining the genes that vary across subpopulations (**Supp. Table S2**) revealed RTK pathway components whose expression differed, including *ERBB3* expression^35^ in BT112, *BRAF* expression in BT228 cells and *FGFR1* expression in BT333 cells (FDR < 1%, **Fig. 2D-E, I-J, and O-P)**). Previous studies have demonstrated that transcriptional plasticity renders a subset of cancer cells resistant to the effect of BRAF kinase inhibition in *BRAF^V^*^600^*^E^*-driven melanoma cells^36^. Similarly, this pre-existing heterogeneity in RTK pathway-associated gene expression may contribute to their ability to adapt and recur after targeting oncogenic driver mutations such as EGFR.

We exposed PDCL models to each of 71 EGFR targeting small molecule inhibitors (**Supp. Table S3**), an EGFR neutralizing antibody (panitumumab), a small molecule activator of EGFR (NSC228155), and puromycin (viability control) each at 4 concentrations, or PBS, Media, or DMSO controls for 24 hours and in replicate for a total of ∼1,800 conditions. These EGFR targeting compounds serve as a comprehensive panel of EGFR inhibitors varying by generation, FDA approval, binding mechanism, activity against EGFR mutants, and known additional targets (**Fig. 2C**, **Supp. Table S3**). We subjected EGFR inhibitor-exposed samples to a modified version of our multiplex single-cell RNA-seq workflow (sci-Plex-v2) that incorporates advances in transcript capture^37,38^ and captures isoform information^38^ while remaining compatible with sci-Plex multiplexing, capturing a total of 181,485 cells with a median of 165 cells per unique combination of PDCL, drug, and concentration (∼1000 cells per PDCL and drug) (**Supp. Fig. 6A-C**), similar to the complexity per condition of prior large-scale single-cell chemical genomics screens^22^. In our large screen, we again identified subpopulations associated with increased expression of relevant RTK pathway genes. Specifically, for each PDCL, 3 clusters were identified among cells exposed to the inhibitor library for 24 hours. For BT112, cluster 1 was enriched for *ERBB3* expressing cells (**Fig. 2G**). Cells exposed to high doses of an EGFRi such as osimertinib, the strongest inducer of the shared EGFR signature in our pilot screen, occupied cluster 1 at a higher rate than DMSO control cells (**Fig. 2H**). In contrast, cells exposed to NSC228155, a small molecule activator of EGFR, were less likely to occupy the *ERBB3* high cluster (**Fig. 2H**). In BT228 cells, cluster 3 (**Fig. 2K**) was associated with *BRAF* high expressing cells (**Fig. 2L**) which appeared to expand as a function of EGFR inhibition (**Fig. 2M**). This effect was more pronounced when examining the proportion of *BRAF* high cells exposed to osimertinib (**Fig. 2N**). In contrast, BT333 *FGFR1* high cells did not appear to expand in response to EGFR inhibition (**Fig. 2Q-S**). These results are consistent with studies demonstrating that heterogeneity in the expression of factors associated with RTK signaling can be selected for upon treatment with inhibitors targeting RTK pathway overactivation^36^.

### The global transcriptional response to EGFR inhibition is only partially shaped by a drug’s effect on proliferative- and resistance-associated gene expression

Exposure to our library of EGFR-targeting compounds had profound effects on transcription, with a mean of 6761 DE genes per PDCL and 662 DE genes per compound (FDR < 1%, **Supp. Fig. 4E**, **Supp. Table S4**). We first investigated how a compound’s impact on global gene expression is linked to its potency in altering expression programs associated with proliferation. Across all PDCLs, we identified subsets of EGFRis that led to decreases in the expression of MKI67 and an aggregate score of proliferative gene expression (**Fig. 3A, Supp. Fig. 5A-C**). Clustering by dose-dependent decreases of MKI67 expression correlated well to luminescence-based viability estimates (**Fig. 3A *top annotation***, 1uM exposure), and subsets of inhibitors with effects on proliferation also displayed substantial effects on global transcription (**Supp. Fig. 6C, Supp. Fig. 7**) . For example, CUDC-101 and tyrphostin9 led to pronounced dose-dependent decreases in MKI67 and proliferative gene expression and had some of our screen’s largest effects on global transcription (as # of DEGs). However, although effects on MKI67 and total gene expression changes were correlated, the relationship was weaker for some inhibitors such as EGFRInhibitor in BT112, afatinib in BT228, and PD153035 in BT333 (**Fig. 3A, Supp. Fig. 8A**). Therefore, the results suggest that the mode of EGFR targeting differentially impacts the expression of diverse gene modules, with or without effects on proliferation.

**Figure 3.**
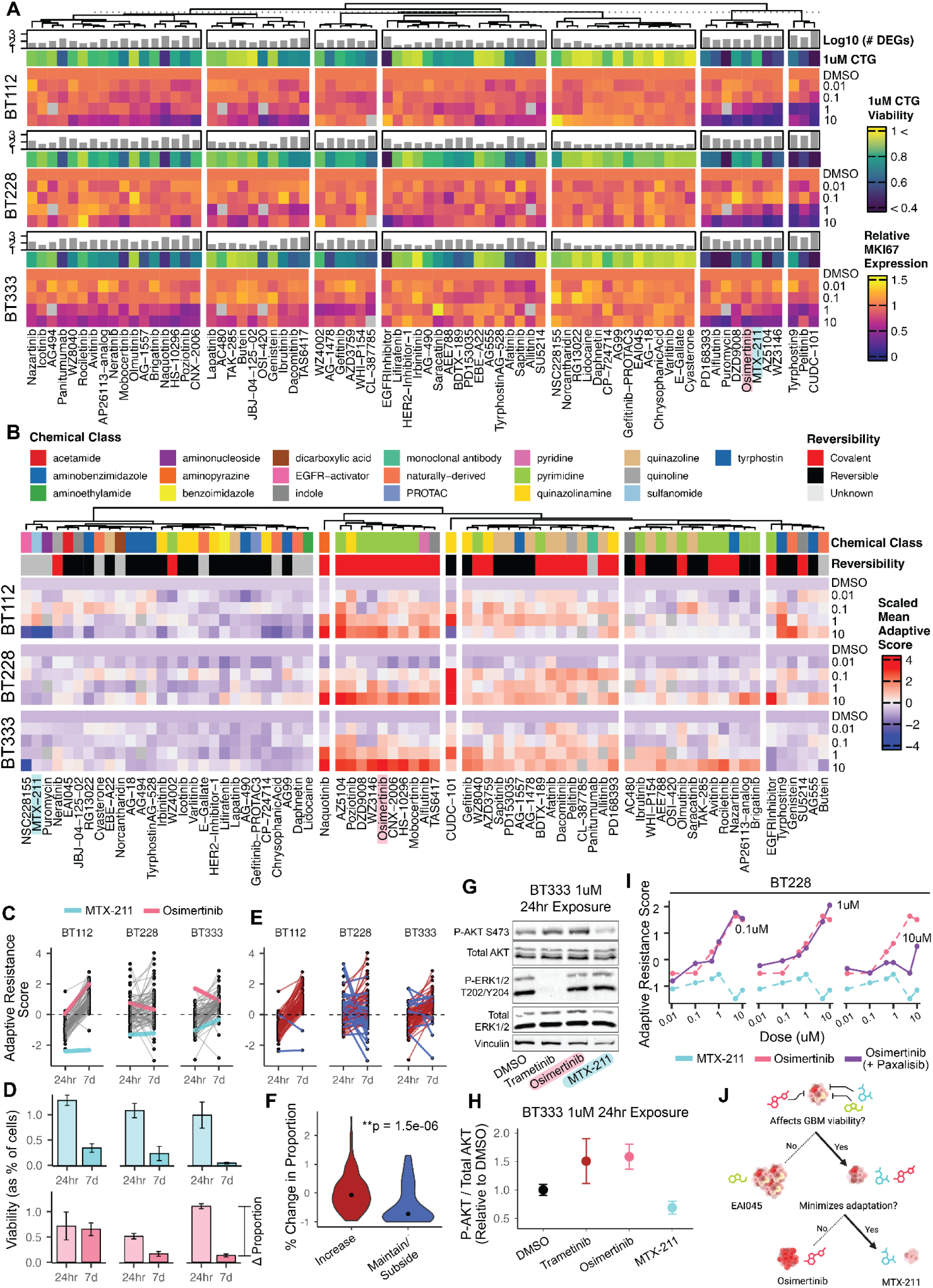
Response to EGFR inhibition is imperfectly correlated to the effect on proliferative transcription. **A.** Relationship between the total number of differentially expressed genes as a function of EGFR inhibitor concentration (quasi-poisson regression, Wald test FDR < 1%) for the specified EGFR inhibitor (*top, bar annotations*) with the relative expression of the proliferation marker MKI67 across BT112, BT228, and BT333 PDCLs. The chemicals osimertinib and MTX-211 are highlighted pink and blue, respectively. Luminescence-based measurement of viability of 1uM of the indicated inhibitor is annotated (*top, color annotations*) **B.** Heatmap of z-scored mean aggregate expression of genes previously associated with adaptation^39^ to inhibition of the RTK pathway for BT112, BT228, and BT333, annotated by chemical classification and EGFR binding mechanism. The chemicals osimertinib and MTX-211 are highlighted pink and blue, respectively. Chemicals are annotated by chemical class and binding mechanism to target (*top annotation*). **C**. Adaptive resistance changes between 24hr and 7d time points for 10uM EGFRi, with osimertinib (pink) and MTX-211 (blue) highlighted. **D**. Viability, as the proportion of cells treated with osimertinib (top) and MTX-211 (bottom) compared to all 10uM treated cells with respect to time point. Osimertinib decreases viability for BT228 and BT333 but not BT112, while MTX-211 decreases viability across PDCLs, consistent with adaptive resistance changes. **E.** Adaptive resistance changes between 24hr and 7d time points for 10uM EGFRi. Inhibitors are highlighted and binned by adaptive resistance transcription dynamics, specifically as increasing vs maintain/subside. **F**. Comparison of changes in proportion between 24hr and 7d for all GBM models between chemicals grouped by adaptive resistance transcription dynamics. The group in which there is minimized induction of the program is characterized by a significantly larger drop in viability (Wilcoxon test). **G.** Quantification of phosphorylated AKT levels for BT333 cells exposed to trametinib, osimertinib, MTX-211 or DMSO control for 24 hours, normalized to DMSO. Points denote the mean and error bars represent standard deviation of 3 replicates. **H**. Representative Western blot results of quantification in (**G**). **I**. Z-scored mean adaptive resistance expression for BT228 cells as a function of dose. Dotted lines represent monotherapy of MTX-211 and osimertinib and solid lines represent osimertinib at a specified constant paxalisib concentration. Paxalisib minimizes the osimertinib dose-dependent induction of adaptive resistance transcription. **J.** A model demonstrating the prioritization of strategies for GBM that decrease viability and minimize the induction of adaptation.

Given the minimal efficacy of EGFR inhibitors in inducing a clinical response in GBM patients in the clinic, we next examined the expression of programs associated with acquired resistance to therapy. Recently, we identified an adaptive resistance signature across established GBM cell lines and glioma stem cell models in response to the inhibition of RTK signaling, most prominently by inhibition of MEK via trametinib^39^. Across the EGFRi signatures from our pilot screen, we found that the shared EGFR signature (**Fig. 1I-J**), associated with decreased patient survival (**Supp. Fig. 3D**) and tumor grade (**Supp. Fig. 3E**), had the highest overlap with our adaptive program (**Supp. Fig. 3C**), suggesting it perhaps reports on tyrosine kinase activity (at baseline or after adaptation). To determine whether EGFR inhibitors in our screen differentially activate this program, we scored cells by its aggregate expression (**Fig. 3B**). Amongst the compounds that led to the strongest induction of the adaptive signature were several compounds with pronounced effects on proliferative gene expression, such as osimertinib, AZ5104 and WZ3146 (**Fig 3A**, **Supp. Fig. 8B**), members of the pyrimidine chemical class of EGFR inhibitors, a class that is enriched amongst compounds with the strongest effect on adaptive gene expression (**Fig. 3B**, *top annotation*). Importantly, this induction was not observed for our puromycin viability control, suggesting that induction of the adaptive program is not merely a consequence of loss of viability. This analysis also identified several compounds that led to decreased proliferative gene expression with minimal induction of adaptive gene expression. For example, MTX-211, a dual EGFR and PI3K inhibitor, displayed the strongest ability to affect proliferative gene expression while minimizing adaptive gene expression in all PDCL models (blue highlight in **Fig. 3A-B**, **Supp. Fig. 8B)**.

We next investigated the association between the induction of adaptive transcription and the ability of cells to persist upon repeated exposure to EGFR inhibitors. We exposed PDCLs to an initial dose of the drug at day 0, a rechallenge at day 3, and profiled cells 7 days post-initial exposure using sci-Plex-v2. Examining the dynamics of adaptive transcription between cells exposed to acute or repeated exposure revealed chemical and PDCL-dependent changes in the acquisition and maintenance of the adaptive resistance program. For instance, adaptive program scores decreased for BT228 and BT333 cells after repeated exposure to osimertinib but not for BT112 (**Fig. 3C**, *top*). Examining the proportion of cells at each condition as a proxy for viability^22^ revealed that decreases in the adaptive program were associated with decreases in cell viability to repeated osimertinib exposure in BT228 and BT333 but not BT112 (**Fig. 3C**, *bottom*). In contrast, cell viability decreased across all PDCLs in response to repeated MTX-211 exposure, consistent with minimal acute or long-term induction of the adaptive transcription program (**Fig. 3D**). We then confirmed, across the same dosing schedule, that exposure to MTX-211 led to a larger decrease in BT112 control growth compared to osimertinib (**Supp. Fig. 9A**). More broadly, we observed that chemicals that increase adaptive transcription upon repeated exposures were less effective in decreasing viability compared to compounds where the program is not induced, but rather maintained or subsided (**Fig. 3E-F**).

The nature of MTX-211 as a dual EGFR/PI3K inhibitor^40^ suggests that combinatorial targeting of PI3K activity is responsible for minimizing the adaptive response to EGFRi, an observation consistent with previous studies^41–43^. We therefore examined the levels of phosphorylated AKT after exposure to 1uM osimertinib, MTX-211, or trametinib, a MEK inhibitor that induces adaptive kinome rewiring response in models of glioblastoma^39^ and breast cancer^44,45^. After 24 hours of exposure, trametinib and osimertinib both increased pAKT compared to vehicle exposure (∼1.5X to DMSO), while cells treated with MTX-211 maintained lower levels of pAKT compared to trametinib, osimertinib and vehicle control (0.56X to DMSO) (**Fig. 3G-H, Supp. Fig. 9B-C**).

While our work suggests that the first-in-class bifunctional compound MTX-211 induces favorable molecular phenotypes by suppressing adaptation, recent studies have identified it as a substrate of the ABCG2 efflux transporter^46^, which restricts drug delivery to the brain^47^. This makes MTX-211 unlikely to possess the ADME properties required for the treatment of brain malignancies. We hypothesized that exposure with a combination of two inhibitors with favorable pharmacokinetic properties could phenocopy MTX-211. To test whether combinatorial targeting of EGFR and PI3K by two brain penetrant inhibitor can recapitulate the properties of MTX-211 we profiled PDCL models for their transcriptional response to osimertinib (EGFRi) and paxalisib (PI3Ki), two clinically-relevant brain penetrant small molecules^48,49^, alone or in combination (**Supp. Fig. 10A**). We observed that the osimertinib induced adaptive program was significantly blocked by increasing concentrations of paxalisib (**Fig. 3I**, **Supp. Fig. 10B-C**). These results demonstrate that combination exposure with two clinically-relevant inhibitors can phenocopy the potentially beneficial properties of an inhibitor profiled in our unbiased survey of EGFR inhibitors.

Taken together, our results provide evidence of a relationship between the induction of an adaptive transcriptional response and the persistence of cells under inhibitor pressure. We demonstrate that the induction of the adaptive transcriptional response can be attenuated by co-targeting of EGFRi and PI3Ki, a finding driven by the transcriptional profile imparted by a dual EGFR/PI3K inhibitor included in the screen. Our results provide evidence that a survey of the differences in the induction of molecular programs across related inhibitors can nominate chemicals or combination strategies that consider metrics of both cytotoxicity and molecular phenotype (**Fig. 3J**).

### A classification of anti-EGFR agents by their induction of molecular programs

We next sought to classify our panel of EGFRis by summarizing their relative effect on gene expression consequent to their unique and shared polypharmacology, resulting in a workflow we term SCHEMATIC (**S**ingle-cell **CHEM**ic**A**l **T**ranscriptom**I**cs-based chemotype **C**lassification) (**Fig. 4A**). We used MrVI^21^, a deep generative model that performs sample stratification at single-cell resolution, to annotate EGFRi-induced transcriptional responses and classify EGFRis by their ability to induce them. MrVI’s hierarchical probabilistic framework presumes cells to be generated from nested experimental designs, such as our study, in which each sample, defined as a drug-dose treatment condition, is drawn from one of several experimental batches. MrVI learns two distinct latent feature spaces from a given scRNA-seq dataset, a sample-unaware *U*-space that is decoupled from the sample-of-origin, and a sample-aware *Z*-space that incorporates sample-of-origin effects while accounting for technical factors across both spaces. These latent spaces can be used to dissect sample-specific heterogeneity in cell states across populations of interest in downstream analyses (**Supp. Fig. 11**).

**Figure 4.**
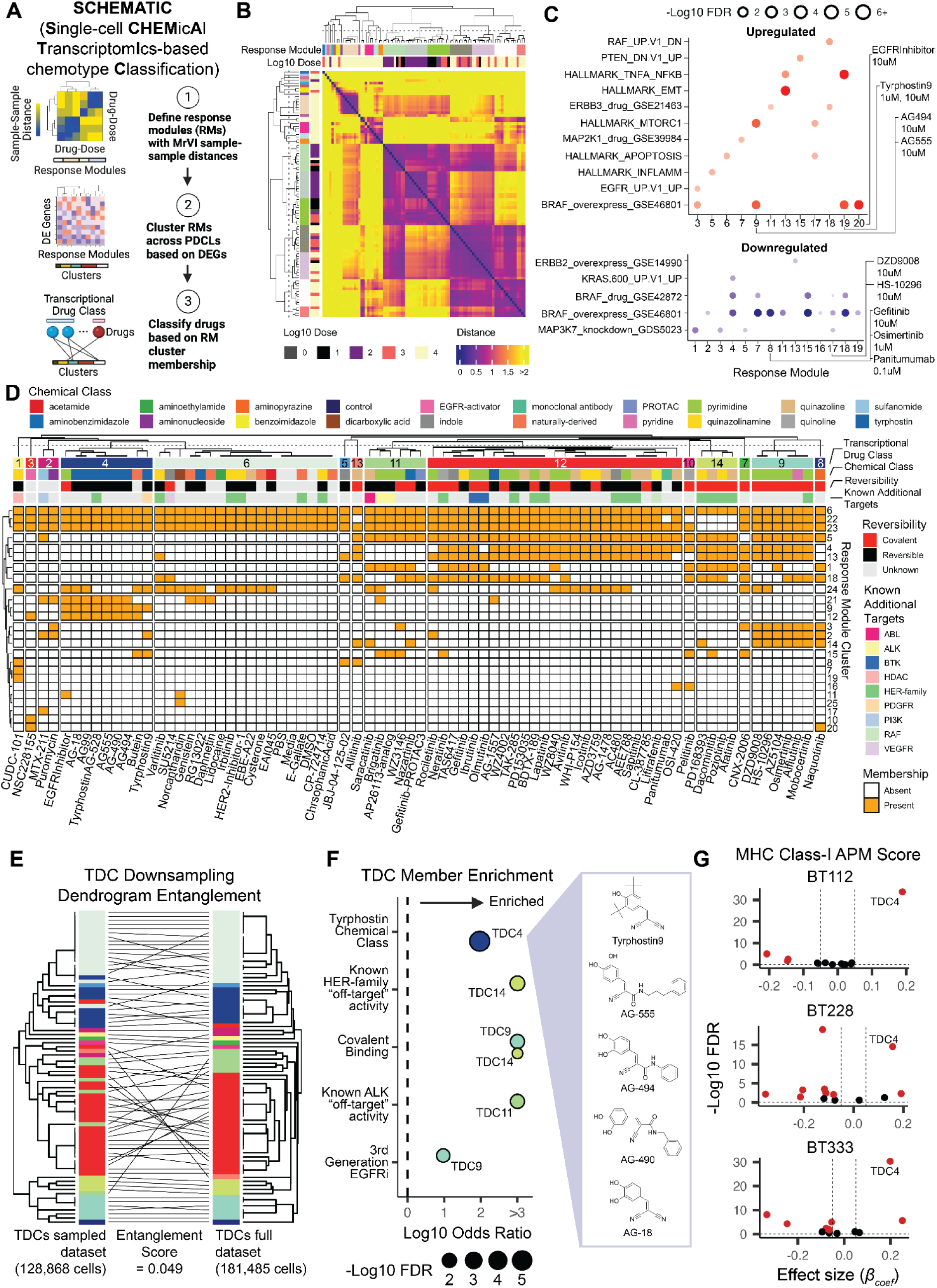
SCHEMATIC EGFR inhibitor classification using MrVI. **A.** SCHEMATIC overview showing MrVI-based RM discovery, covariate-specific DE, and TDC assignment. **B.** Heatmap of the mean normalized distances between drug-dose combinations across BT228 cells, estimated with MrVI. Hierarchical clusters were identified and labeled as Response Modules (RMs) **C.** Gene set enrichment analysis results for the up-regulated (top) and down-regulated (bottom) genes of BT228 RM transcriptional signatures (FDR < 5%). Representative gene sets were chosen for visualization, and full GSEA results can be found in **Supp. Fig. 20**. **D.** Binary heatmap of each drugs’ membership across RM clusters. Hierarchical clustering of the drugs reveals generalized transcriptional drug classes (TDCs) of EGFRi. **E.** Dendrogram entanglement visualizing the similarity of hierarchical clusters of TDCs derived at two different cell depths. Entanglement score of 0 signifies completely matched dendrograms while a score of 1 signifies completely unmatched. **F.** Enrichment of transcriptional drug classes by Fisher’s exact test for select chemical features such as additional or off-targets, binding mechanism to target, or inhibitor generation. **G.** Volcano plots displaying enrichment for APM MHC-CI score across all TDCs for each PDCL. Beta coefficients and FDR values were obtained from a linear regression model, in which APM MHC-CI score was fit dose of chemicals within a TDC. TDC4, represented by tyrphostin-family compounds, is highlighted as enriched for APM MHC-CI in each PDCL.

We trained separate models for each of our 3 PDCLs with unique drug-dose combinations defined as the sample-of-origin and batch replicate as the technical factor. We used the union of the top 100 highly variable genes for each unique PDCL-drug subset as features (total of 2,767 genes). MrVI defines the distance between samples for a given cell as the Euclidean distance between sample-specific counterfactual cell states, also known as the sample-sample distance. The single-cell resolution of the sample-sample distances reveals heterogeneous transcriptional responses to drug-dose treatment, even within individual drug-dose conditions (**Supp. Fig. 12, Supp. Fig. 13 & Supp. Fig. 14**).

Within SCHEMATIC, MrVI then summarizes drug-dose effects to define response modules (RMs) that group conditions inducing similar single-cell programs. Hierarchical clustering of the mean sample-sample distances across cells revealed 22, 21, and 23 distinct groups or RMs of drug-dose combinations that resulted in similar transcriptional responses in BT112, BT228, and BT333 PDCLs (**Fig. 4B**, **Supp. Fig. 15A, & Supp. Fig. 16**). For example, in BT228, control treatments (DMSO, PBS, and media) and cells exposed to several EGFRis at low concentrations comprised RM14, which altogether represent exposures with minimal effects on transcription. The other response groups vary in their composition of chemicals and concentrations; for instance, RM1 and RM3 are comprised of CUDC-101 at high (1uM and 10uM, respectively) concentrations, while RM8 is represented by 29 unique chemicals ranging from 100nM to 10uM (**Supp. Fig. 16**).

We used covariate-specific differential expression analysis to identify differentially expressed (DE) genes in cells of each RM compared to DMSO as a function of the counterfactual-based sample-sample distances identifying 563, 929, and 1021 DE genes for BT112, BT228, and BT333 PDCLs (LFC > 0.1 and FDR < 5%) (**Supp. Fig. 17, Supp. Fig. 18 & Supp. Fig. 19,** see *Methods*). Gene set enrichment analysis across DE genes using the MSigDB Hallmark, MSigDB Oncogenic Signatures, Kinase Perturbations from GEO up and Kinase Perturbations from GEO down gene set collections revealed that RMs are enriched for genes associated with diverse biological processes, including the cell cycle, PI3K/mTOR signaling, kinase perturbation, induction of epithelial to mesenchymal transition (EMT) processes, increased KRAS or BRAF signaling, and TNF-alpha signaling through the NFkB pathway (**Fig. 4C**, **Supp. Fig. 20**).

SCHEMATIC next consolidates RM membership into transcriptional drug classes (TDCs) and visualizes class-specific program strength and heterogeneity. We hierarchically clustered RMs across PDCLs based on the normalized covariate-specific log-fold changes (LFC) in gene expression for the union of RM DE genes (**Fig. 4A, Supp. Fig. 15B**, **Supp. Table 5**). Although these RM clusters are generally occupied by one PDCL, they allow us to further summarize drug effects by collapsing drugs with similar RM cluster membership profiles (**Fig. 4D**, **Supp. Fig. 21 & Supp. Fig. 22**). This process resulted in an unbiased grouping consisting of 14 TDCs across 24 RM clusters with similar PDCL- and drug-specific responses (**Supp. Table 6**).

Seven TDCs consisted of multiple agents, while seven TDCs consisted of single-agents, some of which were expected based on known protein target profiles. For example, CUDC-101 (EGFRi/HDACi) and NSC228155 (EGFR activator, dimerization-domain binder^50^) were classified into distinct groups, TDC1 and TDC3, respectively. Control chemicals DMSO, PBS, and media were grouped in TDC6 along with compounds like EAI045, EBE-A22 (the inactive metabolite of PD153035), and cyasterone all of whom had little effect on transcription and viability (**Fig. 3A, Supp. Fig. 6M**). Similarly, closely related compounds with strong effects on gene expression and similar EGFR off-target profiles grouped together; for instance, the ALK inhibitor brigatinib and its analog-AP26113 were both members of TDC11, and osimertinib and its active metabolite AZ5104 were both members of TDC9. We characterized the TDCs by their enrichment of drugs with shared chemical features (Fisher’s exact test, **Fig. 4F**), including enrichment for known additional or off-targets, binding mechanism to target, or inhibitor generation. Interestingly, TDC4, which grouped separately from the vehicle control containing clade and known clinically-effective EGFR inhibitors, was enriched for tyrphostin class chemicals (AG18, AG99, AG490, AG494, TyrphostinAG-528, AG555, and Tyrphostin9) (**Fig. 4F**, *boxed*). Tyrphostin (“tyrosine phosphorylation inhibitors”) compounds were some of the earliest small molecule ATP-competitive protein tyrosine kinase inhibitors that laid the groundwork for the development of later clinically effective generations of inhibitors^51^. This unbiased grouping of structurally similar tool compounds indicates a shared differential transcriptional signature detected by SCHEMATIC.

Overall chemicals within the same TDC induced similar transcriptional changes than chemicals within other TDCs (**Supp. Fig. 23**). We assessed the robustness of SCHEMATIC assigned TDCs by downsampling cell coverage. We find that our drug-level detection of transcriptional effects was stable across varying levels of cell coverage per condition (**Supp. Fig. 24A-B**). Additionally, within SCHEMATIC, MrVI-derived counterfactual distances were highly correlated at the different coverages (**Supp. Fig. 24C**) and downstream class-level inferences were highly consistent as assessed by dendrogram entanglement (**Fig. 4E**) and cophenetic distance correlation (**Supp. Fig. 24D**). The results underscore the robustness of our assigned TDCs.

Interestingly, several drugs fell into or were excluded from TDCs in a manner that is not readily explained by their specificity (wild type vs mutant EGFR), drug generation, or chemical class. For instance, saracatinib, a SRC family TKI^52^ with an order of magnitude higher activity against wild type vs. mutant EGFR^53^, clustered within TDC11 with WZ3146 and nazartinib, both mutant EGFR (T790M) selective inhibitors^54,55^. Recently, WZ3146 was also found to target the SRC family kinases LYN and FYN^56^, suggesting that nazartinib may have a similar off-target profile. TDC11 also contains the pan-TKI inhibitors brigatinib, its analog-AP26113, and the molecular degrader gefitinib-PROTAC3. Interestingly, structural predictions and in vitro studies suggest that gefitinib can target SRC and the Src family kinase LCK as off-targets^57^. The differential TDC grouping for gefitinib-PROTAC3 and gefitinib and the former’s inclusion in TDC11 may highlight an unknown effect specific to the molecular degrader with regard to decreased activity of Src-family kinases.

### A subset of EGFR inhibitors modify programs associated with tumor cell immunogenicity

GBM is one of the least immunogenic tumor types, displaying low expression of the antigen processing and presentation machinery (APM) and low infiltration of cytotoxic immune cells^58,59^. Several studies have identified changes in tumor cell immunogenicity as a function of inhibiting the kinase activity of EGFR^60–62^ or MEK^63,64^. Furthering prior studies, our profiling across a large number of compounds can provide insight into chemical classes of EGFR inhibitors that drive pro-immunogenic response and report on whether an off-target effect contributes to this clinically beneficial outcome. We therefore evaluated whether any TDCs were enriched for genes associated with the MHC class I (MHC-CI) or MHC class II (MHC-CII) antigen processing and presentation machinery^65,66^. Our analysis identified TDC4, whose membership is comprised of tyrphostin compounds such as AG18, AG99, AG490, AG494, AG555, and tyrphostin9, as significantly enriched for MHC-CI APM expression induction across all three PDCLs (**Fig. 4F-G**).

To more directly determine whether EGFR inhibition leads to increased APM expression in GBM PDCLs, we scored EGFRi-exposed PDCL cells for the expression of MHC-CI and MHC-CII genes and used linear regression to identify compounds with a dose-dependent effect on APM expression. We identified 42 compounds significantly affecting MHC-CI APM expression and 46 that significantly affected MHC-CII APM expression across one or more PDCLs (**Fig. 5A & Supp. Fig. 25A**). Consistent with our TDC-level analysis, there was a marked enrichment for tyrphostin class compounds amongst MHC-CI APM-modifying EGFR inhibitors (**Fig. 5A**). Of note, several of these tyrphostin compounds had minimal effects on proliferative and moderate effects on global gene expression (AG18, AG490, AG494, AG555; **Fig. 3A, Supp. Fig. 6M**) yet led to consistent increases in APM scores. Examining changes at the level of individual genes revealed that changes induced by tyrphostins are distinct from other classes. For example, across conditions that increased MHC-CI APM expression in 2 or more PDCLs, all exposures were associated with increased *B2M*, *CALR*, *CANX*, *HLA-A*, *HLA-B*, and *HLA-C* expression. In contrast to tyrphostins, however, exposure to CUDC-101 either did not alter or was associated with the downregulation of factors such as *ERAP2*, *PSMB5*, *PSMB6*, *PSMB7*, and *TAPBP* (**Fig. 5B**, dashed boxes).

**Figure 5.**
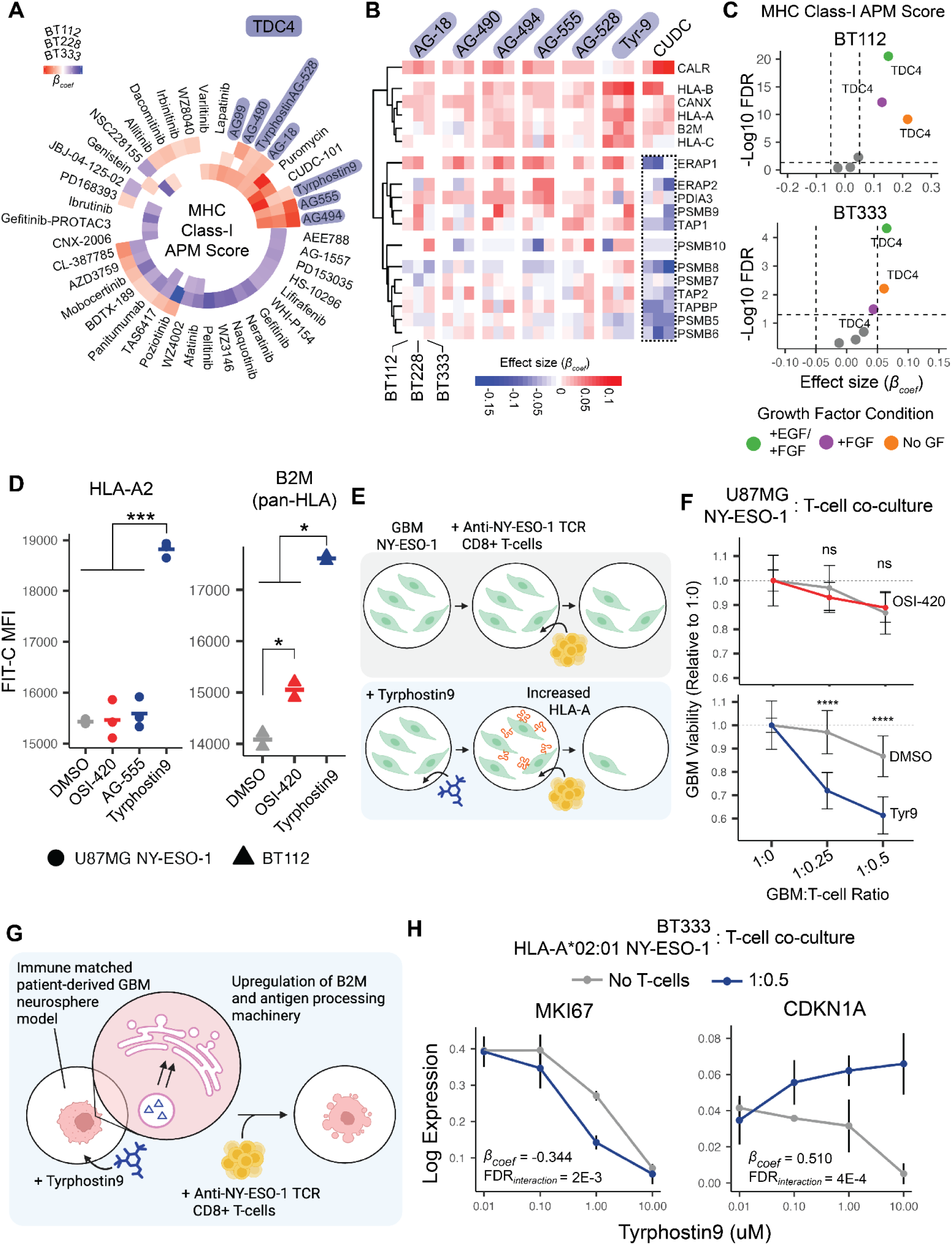
A subset of EGFRis modulate immunogenicity. **A**. Circos heatmaps of beta coefficients for the dose term of a linear regression model of the effect of EGFR inhibition on the expression of genes associated with the MHC class I antigen processing and presentation machinery (APM) in one or more PDCLs. Only significant coefficients (FDR < 5%, Wald test) are shown. **B**. Heatmap of the effect of EGFR inhibition on genes associated with the MHC-CI APM for EGFRis that significantly increase aggregated MHC-CI APM expression in two or more PDCLs from (**A**). **C**. Dose-dependent APM MHC-CI score induction of drugs grouped by TDC4 or not TDC4. TDC4 induces APM MHC-CI expression increase across growth-factor conditions and PDCL. **D**. U87MG NY-ESO-1 HLA-A*02:01 protein expression quantified as mean fluorescent intensity (*** p < 0.001, t-test with Benjamini-Hochberg multiple hypothesis correction) (*left)*. BT112 B2M (pan-HLA) protein expression quantified by mean fluorescent intensity (*p < 0.05, t-test with Benjamini-Hochberg multiple hypothesis correction) (*left).* **E**. Proposed mechanism of tyrphostin9-induced endogenous HLA upregulation increasing GBM susceptibility to T-cell-mediated killing. **F**. U87MG NY-ESO-1 cell viability post-exposure to tyrphostin9 (10uM), OSI-420 (10uM), or DMSO control in combination with increasing T-cell exposure (1:0.25, 1:0.5) quantified as CFSE pixel area of GBM:T-cell conditions 1:0.25 and 1:0.5, relative to 1:0 (drug alone) (n_DMSO,T-cell_ = 18, n_10uM drug,T-cell_ = 9). Wilcoxon test, **** denotes p < 0.0001. **G**. Proposed mechanism of tyrphostin9-induced antigen processing machinery upregulation increasing GBM susceptibility to T-cell-mediated killing in an engineered GBM neurosphere model. **H**. Mean log expression for genes MKI67 (*left*) or CDKN1A (*right*) of engineered BT333 as a function of tyrphostin9 exposure with or without T-cells. Grey points signify BT333 cells cultured alone and blue points signify a BT333:T-cell co-culture at a ratio of 1 GBM cells to 0.5 T-cell. Error bars denote standard deviation from the mean across replicates. The displayed coefficient and FDRs (Wald test) represent the tyrphostin9:T-cell interaction term of a negative binomial regression model for the indicated gene.

Surprisingly, our puromycin viability control induced APM expression in 2 of 3 PDCLs. Previous studies have reported that low-dose puromycin has anti-tyrosine kinase activity independent of its effect on translation^67^. However, given that only a narrow subset of EGFRis modulate APM, it is more likely that other factors downstream of puromycin exposure (puromycilation, inhibition of translation) alter APM expression independent of modulation of tyrosine kinase, or at least EGFR, activity. We also identified broader effects of EGFRis on signaling pathways associated with immunogenicity, such as significant increases in *JAK1*, *STAT1*, and *STAT3* expression by the dual EGFR/HDAC targeting agent CUDC-101 (**Supp. Fig. 25C**). However, while changes to MHC-CI APM expression were observed across all models, changes in MHC-CII APM were largely confined to BT333 (**Supp. Fig. 25A-B**), such as increases in the MHC-CII antigen-presenting subunits *HLA-DMA* and *HLA-DMB*^67,68^ (**Supp. Fig. 25D-E**).

Given that the levels of growth factors added to maintain PDCL cultures may not be representative of the relatively low levels observed in the adult brain^69^. We performed a screen of a subset of inhibitors, examining the growth factor dependence of the drug-induced increase of APM MHC-CI gene expression. Specifically, we exposed EGFR mutant and amplified BT112 and BT333 PDCLs to a subset of EGFRis that reflect distinct TDCs in media supplemented with either 1) +EGF, +FGF (standard media), 2) +FGF (EGF deficient media), or 3) no growth factors (NoGF) **(Supp. Fig. 26A**). As before, we fit a linear regression model of APM MHC-CI score to the dose of chemicals for TDC4 or non-TDC4 compounds for the subset of inhibitors. The dose-dependent transcriptional induction of the APM MHC-CI program was consistent for TDC4 compounds across varied growth factor conditions (**Fig. 5C**), indicating that this potentially actionable phenotype could arise in the lower EGF environment like the brain.

We next confirmed that changes in the APM MHC-CI signature translates to increases in APM expression at the protein level, observing increases in HLA-A and B2M in patient-derived and established GBM models only upon exposure to TDC4 EGFRis (**Fig. 5D** & **Supp. Fig. 23F-G**). We sought to determine whether tyrphostin-induced changes associated with tumor immunogenicity lead to functional changes in the ability of immune cells to target GBM cells for destruction *in vitro*. Accordingly, we first used an allogeneic GBM:T-cell co-culture system consisting of U87MG GBM cells overexpressing the tumor antigen NY-ESO-1^70^ and T-cells expressing anti-NY-ESO-1 T-cell receptors (TCR)^71^, to model GBM:T-cell interactions in vitro as performed by previous studies^72–74^ (**Fig. 5E)**. These engineered GBM models respond to on-target killing by cytotoxic T-cells, requiring less T-cells per GBM cell and time in co-culture compared to their non-engineered parental counterparts^74^. We labeled the engineered GBM cells with CFSE to track changes in cell numbers as a function of fluorescent area (**Supp. Fig. 27A**). We pre-treated NY-ESO-1 GBM cells with a range of concentrations to tyrphostin family EGFRi’s (AG18, AG490, AG494, AG555, tyrphostin9) that increased APM expression across our PDCLs, EGFRi’s that do not alter APM expression but have a range of effects on transcription (CL-387785, OSI-420, osimertinib, PD153035) and DMSO vehicle control. We then co-cultured pre-treated NY-ESO-1 GBM and terminally differentiated anti-NY-ESO-1 CD8^+^ T-cells at GBM:T-cell ratios of 1:0, 1:0.25, and 1:0.5, the latter ratios resulting in approximately 5 to 20% decreases in GBM cell viability 48 hours post-co-culture, respectively. To account for differences in viability across inhibitors, we normalized the fluorescent area to that of the respective no T-cell condition (**Fig. 5F, Supp. Fig. 27B-C**). Exposure of cells to tyrphostin9 led to a significant increase in T-cell mediated killing (**Fig. 5F, Supp. Fig. 27A-C**). In addition, exposure of cells to AG490, AG494, and AG555 led to moderate increases in T-cell mediated killing at the GBM:T-cell ratio of 1:0.5 (**Supp. Fig. 27C**), consistent with their effect on APM expression (**Fig. 5A-B**). The results suggest that tyrphostin9-induced changes, like endogenous HLA-A upregulation (**Fig. 5D**), increase susceptibility to cytotoxic T-cell mediated killing (**Fig. 5F**).

In order to assess the functional effects of tyrphostin9 in a patient-derived GBM neurosphere model, we engineered the PDCL BT333 to express both the HLA-A*02:01 allele and NY-ESO-1 antigen to immune match with anti-NY-ESO-1 TCR T-cells, as previously described^74^. Because of the exogenous overexpression of HLA-A and NY-ESO-1, this model can be used to assess whether changes in the antigen, MHC subunit B2M, or the antigen processing machinery contribute to enhanced T-cell killing (**Fig. 5G**). Engineered BT333 cells were pre-treated with tyrphostin9 at increasing concentrations (0.01, 0.1, 1, and 10 µM) or DMSO control for 24 hours. Following drug pre-treatment and IFN-γ stimulation, the medium was removed and replaced with fresh medium, either without or with CD8⁺ NY-ESO-1 TCR-transduced T cells, at a GBM:T-cell ratio of 1:0.5, a modest ratio intended to highlight enhanced killing from baseline. We then harvested GBM nuclei at 18 hours post-co-culture for single-cell RNA-seq profiling, capturing 21,892 cells with a median of 890 cells per condition (**Supp. Fig. 27D**). We examined the impact of tyrphostin9 on T-cell induced changes in *MKI67*, and *CDKN1A*, which encodes the cell-cycle inhibitor and p53 target p21 and serves as a marker of cell cycle arrest^75^. Tyrphostin9 alone induces a decrease in *MKI67* (**Fig. 5H**, *left*, grey points); however, the combination of T-cells and tyrphostin9 led to a strong decrease in *MKI67* expression, apparent at the intermediate 1uM exposure. Negative binomial regression of the interaction between tyrphostin9 dose and T-cell addition confirmed a significant decrease in *MKI67* (FDR = 0.002). For *CDKN1A*, co-exposure to cytotoxic T-cells and tyrphostin9 led to a robust increase in *CDKN1A* expression (FDR = 0.0004), representing a reversal of the dose-dependent decrease observed to tyrphostin9 alone (**Fig. 5H**, *right*). Collectively, the results of GBM:T-cell co-culture experiments suggest that tyrphostin9 can enhance the ability of cytotoxic T-cells to induce tumor cell killing in an on-target HLA-dependent and independent manner.

To the best of our knowledge, the known polypharmacology of these inhibitors does not provide a straightforward explanation of which molecular target is responsible for these changes in immunogenicity, and further screening with orthogonal measurement is needed to identify the causal targets. However, our results suggest that EGFR is not the primary effector, given the small subset of compounds that induce the response (**Fig. 5A-B**), their differing potency in modifying proliferative gene expression (**Fig. 3A**), and the lack of compensatory feedback that characterizes the most potent EGFR inhibitors in our screen (**Fig. 3B**). Our approach nominates tyrphostin9 as a compound for further medicinal chemistry review and *in vivo* mechanistic studies. Overall, our findings further imply that the choice of EGFR inhibitor can result in complex molecular state changes with the potential to alter the immunogenicity of GBM.

## Discussion

Biochemical and chemical proteomic studies have established the fact that kinase inhibitors have varying degrees of selectivity for their intended targets^76–79^. In some cases, a drug’s higher degree of polypharmacology may enhance clinical efficacy^80^, while for others, off-target effects may result in unacceptable side-effects that lead to its failure in the clinic. In the context of glioblastoma, clinical trials for EGFR inhibitors have yielded little success despite the high prevalence of activating EGFR mutations in the patient population and despite the success in targeting EGFR across other tumor types. It is apparent that the molecular response in targeting this oncogenic activity is not fully understood. Here, in targeting EGFR with a diverse inhibitor library composed of small molecules and a biologic, each with different chemical structure and binding properties, we define a landscape of responses to an entire inhibitor class and further group the chemical agents into transcriptional drug classes based on the shared induction of molecular responses.

Our high-throughput targeted screens are complementary to traditional measurements of inhibitor efficacy like phosphorylation measurement or target binding assays. Capturing downstream transcriptional snapshots of inhibitor responses allowed for the comparison of inhibitor efficacy by standard screening metrics such as proliferation (e.g. MKI67 or aggregate proliferation score) that are similar to those obtained with traditional viability-based assays.

However, we show that an inhibitor’s impact on viability is not always correlated with global transcriptional response, indicating that inhibitor responses extend beyond viability, consistent with previous chemical genomic profiling efforts^5,22^. Moreover, we show that EGFRi differentially induce an adaptive resistance molecular response recently associated with MEK inhibition^39^ and that accounting for this adaptive response will help lead to identifying clinically effective compounds. We further validated the impact and relevancy of select phenotypes by exploring their dependence on exogenous growth-factor levels. It is important to note that the transcriptional changes measured in our study may be dependent on one or more mechanisms of EGFR activation. For instance, EGFR ecDNA, chromosomal amplification, and mutation may each contribute to pre-exposure cell state variability and chemotype-specific responses.

The generation of single-cell chemical genomics datasets necessitates the development of novel computational approaches that efficiently summarize the large-scale high-dimensional data to gain novel biological insight into the effects of chemical perturbation on cellular and molecular states. Recent studies have developed techniques that study transcriptional differences in patients, cell types/states, and molecular perturbation screens^81–83^. In addition to developing a large data resources that can help refine these efforts, within SCHEMATIC, we demonstrate the suitability of MrVI^21^, a recently developed deep generative model designed to estimate sample (drug-dose) heterogeneity at the cellular level while jointly correcting for batch effects, for summarizing transcriptional responses in large-scale genomics screens and grouping chemical agents by their shared induction of cellular states. Our SCHEMATIC workflow provides a general framework to group related inhibitors by their single-cell programs, linking mechanisms to actionable outcomes. SCHEMATIC enabled the discovery of robust and likely off-target effects related to increased immunogenicity due to the induction of the antigen processing and presentation machinery and provides rationale for further pre-clinical *in vivo* studies, including *in vivo* single-cell perturbation screening^84^. Future rational design of inhibitors could incorporate similar approaches to identify compounds that simultaneously block oncogenic kinase activity and have desirable off-target effect profiles that induce responses synergistic with combination immunotherapy.

Our results provide motivation for the single-cell profiling of chemical inhibitors against a single target toward the multiplex characterization and annotation of designed and off-target molecular programs. Thanks to advances in combinatorial indexing single-cell RNA-seq library generation methods^37,38^, hundreds of thousands to millions of cellular transcriptomes across thousands of unique conditions can be obtained robustly and at reasonable cost^22,39,85^, demonstrated recently with the Tahoe-100M dataset^86^. Previous studies have included large-scale efforts to establish atlases of molecular responses to a massive amount of perturbations across a spectrum of cancer cell lines, utilizing probe-based fluorescence for targeted gene expression measurements^5^, with limitations in drug repurposing applications^87^. While these efforts can be beneficial for applications like annotating broad inhibitor mechanisms of action, there is a need for high-resolution cellular resolved characterization of responses to chemicals, limited to a chemical class and targeting a specific cancer type. Future endeavors that would further strengthen our approach include novel ways to incorporate methods like lineage tracing^88^, beneficial for answering questions of clonal expansion under inhibition^89^, across large-scale experiments with limited cell, and therefore clonal, coverage per condition. Additionally, while we obtain a global view of EGFRi responses by studying three genetically-diverse models, future screens will need to incorporate EGFR genotype as a variable across a larger number of models to discover the effects of GBM-specific mutations or through the study of isogenic alterations in wildtype backgrounds. Beyond the study of EGFR inhibitors, SCHEMATIC can generalize to other targets and chemotypes, guiding combination design and pre-clinical triage across physiologically-relevant natively heterogeneous pre-clinical tumor models.

## Materials and Methods

### Cell culture

A172, T98G, and U87MG glioblastoma cell lines were purchased from ATCC. Cells were cultured in DMEM media (ThermoScientific) supplemented with 10% fetal bovine serum and 1% penicillin/streptomycin (P/S, ThermoScientific) according to ATCC instructions. Glioma neurosphere cell lines BT112, BT228, and BT333 were obtained from the DFCI Center for Patient Derived Models (CPDM) under a material transfer agreement and maintained as described previously (Touat et al 2020 Nature). Briefly, cells were grown in Neurocult NS-A Proliferation Media (StemCell) supplemented with 0.0002% heparin (StemCell Technologies), EGF (20 ng/ml), and FGF (20 ng/ml; Miltenyi) in a humidified atmosphere of 5% CO2 at 37 °C on low-attachment plates ^90^ and were dissociated with Accutase (StemCell Technologies) for passaging and plating.

### Pilot screen chemical perturbation procedure and materials

Cells were plated at a density of 2.5E4 cells/well onto 96-well flat-bottom plates for adherent cell lines and V-bottom plates for spheroid cell lines and were allowed to acclimate overnight. Chemical agents (SelleckChem) were purchased as a powder and resuspended in DMSO as 10mM (afatinib, CUDC-101, EAI045, neratinib, osimertinib) or 5mM (brigatinib) stocks according to the manufacturer’s instructions. Dose concentrations (5nM, 10nM, 50nM, 100nM, 500nM, 1uM, 5uM, 10uM) were prepared in media as 10X stocks and were added to cells with all wells were normalized to 0.2%v/v DMSO (final conc. in well). Nuclei were hashed, fixed, and harvested 24 hours post-exposure.

### Large screen chemical perturbation procedure and materials

Cells were plated at a density of 2.5E4 cells/well on 96-well V-bottom plates as in the pilot screen. Chemical agents (SelleckChem) were purchased as frozen 10mM stocks in respective solvents (DMSO, 1X PBS). 10-fold dose concentrations (0.01uM, 0.1uM, 1uM, 10uM) were prepared in media at 10X stocks and added to cells. Wells were DMSO or PBS normalized according to drug solvent, and vehicle-treated wells were included. Nuclei were hashed, fixed, and harvested at 24 hours post-exposure. For 7d time point exposures, cells were drugged initially in the same manner as in the 24hr condition. At 3 days post-exposure, wells were supplemented with 10uL of drug in media equimolar to the well condition. Finally, nuclei hashed, fixed, and harvested at 7 days post-initial exposure.

### CellTiter-Glo and cell counting viability measurements

For pilot screen inhibitors, A172, BT333, U87MG, and T98G cells were plated at 5E3 cells in 96-well plates and were acclimated overnight. As before, dose concentrations (5nM, 10nM, 50nM, 100nM, 500nM, 1uM, 5uM, 10uM) were prepared in media as 10X stocks and were added to cells in duplicate with all wells were normalized to 0.2%v/v DMSO (final conc. in well). After 4 days of exposure, CellTiter-Glo (Promega) was prepared according to manufacturer instructions and added 1:1 to cells. After shaking and incubating in foil for 10 minutes, the mixture was moved to 96-well opaque wall plates, and luminescence was read with a BioTek Synergy H1 plate reader. A four parameter dose-response curve was fit to luminescence relative to DMSO using the R package *drda* for further calculation of IC50 and area under the curve. For large screen inhibitors, BT112, BT228, and BT333 cells were plated at 5E3 cells in 96-well plates and were acclimated overnight. Inhibitors or vehicle controls were diluted in media to 10X stock for 1uM final concentration and were added to cells in duplicate. After 5 days of exposure, CellTiter-Glo was added to cells and measured as with the pilot screen inhibitors. Viability to 1uM exposure was calculated as luminescence relative to DMSO.

For direct comparison of MTX-211 and osimertinib viability, BT112 cells were plated at 1E6 cells in 10cm dishes and were acclimated overnight. Cells were exposed to 1uM osimertinib, MTX-211, or DMSO control in triplicate. An equimolar re-challenge was added at 4 days post-initial exposure. At 7 days post-initial exposure, cells were collected, dissociated with Accutase, and counted with trypan blue using the Countess cell counter (Invitrogen). Live cell counts were normalized to DMSO control for comparison.

### Dual-inhibitor phenocopy experiment procedure and materials

Paxalisib (GDC-0084, SelleckChem) was resuspended in DMSO and stored as 10mM stocks. Cells were plated at a density of 2.5E4 cells/well on 96-well V-bottom plates and were acclimated overnight as in the pilot screen. For monotherapy exposures, dose concentrations (0.01uM, 0.1uM, 0.5uM, 1uM, 5uM, 10uM) were prepared in media as 10X stocks. For paxalisib + osimertinib combination exposures, osimertinib concentrations (0.01uM, 0.1uM, 0.5uM, 1uM, 5uM, 10uM) were prepared in paxalisib (0.01uM, 0.1uM, 1uM, 10uM) containing media at 10X stocks. All conditions were normalized so that final DMSO concentrations in wells was 0.2%v/v. Chemical stocks were added to cells at 1:10 dilution, and nuclei were hashed, fixed, and harvested at 24 hours post-exposure.

### Growth-factor condition and chemical perturbation combination procedure and materials

BT112 and BT333 cells in standard culture were replated in 10cm dishes in NS-A proliferation media supplemented with either 1) +EGF (20 ng/ml)/+FGF (20 ng/ml), 2) +FGF (20 ng/ml), or 3) no growth factors (NoGF). After 96 hours of culture, cells were dissociated and plated at a density of 2.5E4 cells/well under respective growth factor conditions. After 24 hours of acclimating in the 96 well plate, a narrow set select chemicals (brigatinib, saracatinib, CUDC-101, osimertinib, MTX-211, AG-18, AG-490, AG-494, tyrphostinAG-528, AG-555, tyrphostin9, afatinib) or DMSO was added to cells at 7 dose concentrations (5nM, 10nM, 50nM, 100nM, 500nM, 1uM, 5uM, 10uM) as in the pilot screen under respective growth factor conditions. Nuclei were hashed, fixed, and harvested at 24 hours post-chemical exposure.

### Detection and quantification of downstream phosphorylated MAPK proteins

BT333 cells were plated in 6-well plates at a density of 5E5 cells/well and were acclimated overnight. Cells were exposed to 1uM trametinib (Selleck), osimertinib, MTX-211, or DMSO control (0.01%). After 24 hours of exposure, neurospheres were collected, centrifuged at 200xg for 5 minutes at 4°C, and washed twice with cold 1X PBS. Pellets were snap frozen with liquid nitrogen and stored at -80°C for further processing. Cells were washed with PBS and lysed in Phosphate Lysis Buffer (pH 7.4; 50mM sodium phosphate (38mM sodium phosphate dibasic, 12mM sodium phosphate monobasic); 1mM sodium pyrophosphate; 20mM sodium fluoride; 2mM EDTA; 1% Triton X-100) supplemented with protease and phosphatase inhibitors (1 mM PMSF; 50µg/mL leupeptin; 1mM Na_3_VO_4_; 2mM benzamidine; 1µM microcystin; 1mM DTT). Lysates were briefly sonicated prior to quantification by Bradford Assay. Samples were boiled in sample buffer (250mM Tris HCl, 8% (w/v) SDS, 40% (v/v) glycerol, 80µg/mL bromophenol blue; 2.86M β-mercaptoethanol) for 5 minutes at 95 °C. 20 µg protein per sample was analyzed by 10% SDS–PAGE and transferred to 0.45µM Nitrocellulose membranes (BioRad) for 2h30mins at 90V in transfer buffer (200 mM Glycine, 25 mM Tris Base, 20% Methanol) at 4 °C. Membranes were blocked in 5% non-fat dry milk solution in PBST for 1 hour at room temperature, washed with PBST, and incubated overnight at 4 °C with primary antibody (1:1000 in 1% BSA). Primary antibodies were as follows: Phospho-AKT (Ser473) Rabbit Monoclonal Antibody (D9E, Cell Signaling); AKT (Pan) Rabbit Monoclonal Antibody (C67E7, Cell Signaling); Phospho-p44/42 MAPK (ERK1/2)(Thr202/Tyr204) Rabbit Monoclonal Antibody (D13.14.4E, Cell Signaling); p44/42 MAPK (Erk1/2) Mouse Monoclonal Antibody (L34F12, Cell Signaling); Vinculin Rabbit Monoclonal Antibody (E1E9V, Cell Signaling). Membranes were washed with PBS-T, incubated with secondary antibodies (1:5000 in 5% non-fat dry milk solution in PBST) for 1 hour at room temperature. Secondary antibodies were: IR Dye 800cw Donkey anti-rabbit and Donkey anti-mouse (Licorbio), and washed in PBST before developing with near-infrared fluorescent imaging (Licor).

### Nuclei hashing and fixation

Nuclei hashing and fixation procedures were adapted from Srivatsan et al. and Sziraki et al. Briefly, adherent cells were trypsinized and moved to V-bottom plates, and spheroid cells were dissociated in place. Upon washing with ice-cold 1X PBS, cells were lysed with EZ Lysis Buffer (Sigma) supplemented with 1% diethyl pyrocarbonate (Sigma), 0.1% SuperaseIn RNase Inhibitor (Thermo), and 500 fmol of hashing oligo. After lysis, nuclei were fixed with the addition of 1.25% formaldehyde in 1.25X PBS (final well conc. 1% and 1X, respectively) and incubated on ice for 10 minutes. Nuclei were pooled into a plastic reservoir and moved into a 50mL conical for centrifugation at 650xg for 5 minutes at 4°C. Supernatant was removed from the nuclei pellet, and nuclei were washed once with nuclei suspension buffer (NSB; 10 mM Tris-HCl, pH 7.4, 10 mM NaCl, 3 mM MgCl2, 1% Superase RNA Inhibitor (Thermo Fisher), 1% 0.2mg/mL Ultrapure BSA (New England Biosciences)). Nuclei were resuspended in NSB, slow-frozen in 10% DMSO, and stored at -80°C until sci-RNA-seq processing.

### Library preparation and sequencing

Hashed nuclei were thawed and subjected to 3-level combinatorial indexing protocols adapted from previous methods (Cao et al, Martin et al, Sziraki et al). Nuclei were spun, resuspended in NSB, and sonicated at low power for 12s (Bioruptor). Upon counting, 21uL nuclei were moved to 96-well low adhesion PCR plates with 2uL 10mM dNTP, 2uL 100uM indexed oligo-shortdT primers, 2uL 100uM indexed random hexamer primers, and 14uL of a reverse transcription master mix consisting of 14.29% 100mM DTT, 14.29% 100mM RNaseOUT Ribonuclease Inhibitor, 57.14% 5X SuperScript IV First-Strand Buffer, and 14.29% SuperScript IV Reverse Transcriptase. Reverse transcription was carried out with an increasing temperature gradient. Post-reverse transcription, nuclei were pooled and distributed as 10uL into a 96-well plate(s) for ligation steps. Briefly, 8uL of indexed ligation primers were added to each well, along with a 4.8uL 3:2 master mix of T4 ligase buffer:T4 ligase (New England Biosciences, NEB) and 9.4uL of nuclei buffer with BSA (NBB; 10 mM Tris-HCl, pH 7.4, 10 mM NaCl, 3 mM MgCl2, 1% 0.2mg/mL Ultrapure BSA). Ligation was carried out at 25°C for 1 hour. Resulting nuclei were pooled, washed with NBB, and distributed as 1500 nuclei in 5uL NBB per well, where some plates were stored for future processing. Next, 5uL of a second strand synthesis mix consisting of 60% elution buffer (Qiagen), 27% second strand synthesis buffer (NEB), and 13% second strand synthesis enzyme mix (NEB) was added, and second strand synthesis was carried out at 16°C for 3 hours. Post-second strand synthesis, tagmentation was performed at 55°C for 5 min after the addition of 1/50uL of N7-adaptor loaded Tn5 and subsequent quenching with DNA binding buffer (Zymo) for 5 min at room temperature. Resulting dsDNA was purified using a 1X SPRIbead clean-up within the 96-well plate, and the resulting product was subjected to USER digestion (80% ddH2O, 10% 10X rCutsmart, 10% USER enzyme (NEB)). The dsDNA was eluted in buffer EB then moved to a clean 96-well plate. In addition to 16uL of eluted product, 2uL P5 PCR primer and 2uL P7 PCR primer were added to wells in an indexed well-specific combination. Further, 20uL 2X NEBnext PCR master mix (NEB) was added, and PCR to add the adaptors was carried out. The final PCR product was pooled and subjected to a 0.7X SPRIbead cleanup for library cDNA purification and 1X cleanup for hash fraction purification. Library concentrations were determined by Qubit (Invitrogen) and were visualized by TapeStation DNA D1000. The resulting libraries were sequenced on the Illumina NextSeq550 for the pilot screen or on the Element Biosciences AVITI for the large EGFRi screen and PI3Ki + EGFRi screen, according to the manufacturer’s instructions. The growth-factor follow-up screen was sequenced on Illumina NovaSeq X Plus by Novogene Co.

### Data preprocessing and generation of count matrix

Raw base call files were obtained from Illumina BaseSpace or AVITI storage and were used to generate fastq files using bcl2fastq v2.20.0.422 or bases2fastq version 1.5.0.962525890, respective of sequencing platform. A custom data processing pipeline, adapted from Srivatsan et al., was used to process fastq data into a single-cell count matrix. First, reverse transcription and ligation barcodes were assigned to reads with a mismatch allowance of 1bp, and reads assigned to oligo-shortdT primers were separated from those assigned to random hexamer primers. After index assignment, polyA sequences were trimmed using TrimGalore version 0.6.10 and CutAdapt version 2.6. Upon polyA trimming, reads were aligned to human GRCh38 using the STAR aligner version 2.7.9a. Aligned reads were filtered for quality and duplicates and were assigned to genes using bedtools version 2.26.0, as described previously. The resulting unique read assignments from both primers were combined and collapsed by cell and gene, and a *celldataset* (CDS) object was generated using the raw sparse count matrix, cell annotations, and gene annotations with the R package *monocle3*. Cell barcodes were determined to be cells upon filtering the CDS by a UMI cutoff determined visually with the kneeplot of cell rank by UMI count. Lastly, doublets were detected with *scrublet* and were filtered based on the doublet score distribution.

In parallel, hash assignments were determined from demultiplexed untrimmed oligo-shortdT reads as described previously (Srivatsan et al, McFaline-Figueroa et al). Briefly, hash barcodes were assigned to reads with a mismatch allowance of 1bp and if the read was adjacent to repeated A sequences, corresponding with hash sequence design. Duplicate hash reads were filtered by UMI and were collapsed into hash assignment counts by cell. Hashes were assigned to cells by two criteria: (1) a cell having ≥20 (pilot) and ≥5 (large screen) hash UMIS and (2) a ratio of the cell’s top hash UMI to second best hash UMI of 3 (pilot) or 2.5 (large screen). The *monocle3* package was used to manipulate ^25,91,92^, batch align ^27^, and visualize ^28^ the resulting data.

### Defining shared inhibitor signature

For the pilot EGFRi screen, differentially expressed genes (DEGs) were calculated by fitting expression of a gene in each cell line/drug condition to a generalized linear model (GLM) modeled as a function of dose and replicate using the R package *monocle3*’s *fit_models* function. The tests were limited to genes expressed in at least 1% of all cells in the experiment. P-values for each DEG test were FDR corrected (Benjamini-Hochberg), and significant DEGs were defined as FDR < 0.01 and a normalized effect magnitude (beta coefficient) of > 0.05. The DEGs associated with each drug were joined for each cell line, and signatures were defined as the unique intersection of DEGs between all combinations of agents used. Of note, the inhibitor EAI045 was excluded from signature analysis given its low number of DEGs indicating its overall low impact on transcription. Only DEG intersections with at least 30 genes were analyzed. For each cell, genes for each signature were size-factor normalized, log transformed, and aggregated to amount to a signature score.

### Patient survival analysis

GBM TCGA data (Cell 2013) was accessed and downloaded through the cBioPortal for Cancer Genomics, specifically clinical survival information and paired RNASeqV2 mRNA expression values. Normalized bulk expression values were log10(RSEM) transformed, and signature scores for each donor were calculated as the sum of expression across signature genes. Donors were categorized as high or low signature expression of the signature based on top and bottom 50% of the cohort. Using *survival, ggsurvfit,* and *condsurv* R packages, Kaplan-Meier curves were generated for each group and compared for significant differences in survival using the Gehan-Wilcoxon test or Mantel-Haenszel as denoted. For signature comparison of other cancer types and glioma grade, TCGA PanCancer Atlas data was downloaded and processed in the same way.

### Differential gene expression analysis, calculation of aggregated gene scores for proliferation, resistance, and APM gene programs, and clustering analysis

As before, differentially expressed genes (DEGs) were calculated by fitting expression of a gene in each cell line/drug condition to a generalized linear model (GLM) modeled as a function of dose and replicate using the R package *monocle3*’s *fit_models* function. The tests were limited to the union of genes expressed in at least 1% of cells within each PDCL. P-values for each DEG test were FDR corrected (Benjamini-Hochberg), and significant DEGs were defined as FDR < 0.01 and a normalized effect magnitude (beta coefficient) of > 0.05.

Proliferation, resistance, and antigen-presentation machinery scores were calculated as in Srivatsan et al^22^. Briefly, for each cell, raw gene expression for genes within a program were retrieved from the count matrix and were size-factor normalized. The normalized expression values were aggregated by group and log normalized with a pseudocount of 1. Downstream scores were visualized as group medians or z-scored group means using R packages *Pheatmap* and *ComplexHeatmap*. GLMs of aggregate scores as a function of dose were fit using the R package *speedglm*. For resistance program scores that include both 24hr and 7d exposures, mean aggregate expression was z-scored jointly to visualize changes over time. Clustering on respective datasets was performed using *monocle3*’s *cluster_cells* function.

### U87MG NY-ESO-1 generation and induction of cytotoxic T-cell killing

U87MG were engineered to express the antigen NY-ESO-1 constitutively. Briefly, a plasmid containing the sequences for NY-ESO-1 (CTAG1B) was ordered from OriGene Technologies. qPCR were performed to obtain the NY-ESO-1 fragments with forward and reverse primers. Overhangs were added to NY-ESO1 fragments by gibson ligation to insert into a vector backbone containing a blasticidin resistance sequence. Virus was produced by transfecting HEK293T cells with the plasmid and the Lipofectamine 3000 kit (Thermo Fisher Scientific). U87MG were transduced with the supernatant containing the NY-ESO-1 virus. After 72 hours of viral exposure, U87MG NY-ESO-1 cells were selected with 10ug/mL blasticidin in culture media for 96 hours. Cells were regularly exposed to 1ug/mL blasticidin throughout expansion to ensure blasticidin resistance and NY-ESO-1 expression.

For the GBM cytotoxic T-cell killing experiment, terminally differentiated human CD8^+^ anti-NY-ESO-1 T-cells, derived by activation of CD8^+^ T cells via antigen presenting cells with NY-ESO-1 peptide, were ordered from Charles River Lab. T cells were thawed according to the manufacturer’s protocol and used for the experiment immediately. U87MG NY-ESO-1 cells were cultured and were subjected to TrypLE (Gibco) to be made single-cell in suspension. Cells were stained with CellTrace CFSE Cell Proliferation Kit (Invitrogen) according to instructions (5uM CFSE in 1X PBS, cell density 1E6/mL) to emit fluorescence at 488nm and were subsequently plated in a 96-well flat bottom plate at a density of 10E3 cells/well, in media consisting of DMEM, 10% FBS, and 10mM penicillin-streptomycin, as well as in standard culture conditions of 37°C, humidified 5% CO2/balance air environment. After adhering and acclimating overnight, the cells treated with 10uM EGFRi (AG18, AG490, AG494, AG555, tyrphostin9, CL-387785, OSI-420, osimertinib, or PD153035) or DMSO control — the inhibitors span transcriptional responses of proliferation, adaptive resistance, and antigen presentation. After 24 hours of EGFRi pre-treatment, EGFRi-supplemented media was removed, and the GBM cells were exposed to T-cells at various GBM:T-cell ratio (1:0 i.e no T-cells, 1:0.25, and 1:0.5). In total, there were 18 well replicates per DMSO + T-cell condition, and 11 well replicates for each EGFRi (excluding Tyrphostin9) + T-cell condition, and 9 well replicates for Tyrphostin9 + T-cell condition. T-cells and U87MG NY-ESO-1 cells were co-cultured in complete T-cell culture media consisting of RPMI 1640 supplemented with 10mM HEPES, 10mM L-glutamine, 10% FBS, 0.34% β-mercaptoethanol, and 10mM penicillin-streptomycin, as in ^93,94^. After 48 hours post-T-cell exposure, plates were imaged with the Zeiss confocal microscope and software, capturing 4x4 tiles of 50x total magnification images of wells with both brightfield and 488nm fluorescence. Tile images were stitched, exported as .tif files, and analyzed using a custom Python analysis pipeline implemented with OpenCV (cv2) and NumPy, as described in ^74^.

Briefly, CFSE (488nm) images were loaded and corrected for uneven background. Cells were segmented by fluorescence, and the area occupied was measured in pixels. The resulting area measurements were exported for downstream analysis and visualization in R. The well area for each EGFRi + T-cell condition was aggregated and normalized to the respective plate-specific drug-alone condition CFSE area.

### BT333 HLA-A*02:01 NY-ESO-1 generation and induction of cytotoxic T-cell killing

BT333 were engineered to express the allele HLA-A*02:01 and antigen NY-ESO-1 constitutively. NY-ESO-1 overexpression was established for BT333 as described for U87MG. For HLA-A, a plasmid containing the sequence for HLA-A*02:01 was ordered from AddGene (pENTR-HLA-A0201-His, Plasmid #108213). qPCR were performed to obtain the HLA-A*02:01 fragments with forward and reverse primers. Overhangs were added to HLA-A*02:01 fragments by gibson ligation to insert into a vector backbone containing a puromycin resistance sequence. Virus was produced by transfecting HEK293T cells with the plasmid and the Lipofectamine 3000 kit (Thermo Fisher Scientific). BT333 NY-ESO-1 cells were transduced with the supernatant containing the HLA-A*02:01 virus. After 72 hours of viral exposure, BT333 cells were selected with 1ug/mL puromycin in culture media for 96 hours. Cells were regularly exposed to puromycin and blasticidin throughout expansion to ensure resistance and HLA-A*02:01 and NY-ESO-1 expression.

For the engineered BT333:T-cell sci-Plex experiment, terminally differentiated human CD8^+^ anti-NY-ESO-1 T-cells, derived by activation of CD8^+^ T-cells via antigen presenting cells with NY-ESO-1 peptide, were ordered from Charles River Lab. T-cells were thawed according to the manufacturer’s protocol and used for the experiment immediately. BT333 HLA-A*02:01 NY-ESO-1 cells were cultured and were subjected to Accutase (StemCell Technologies) for single-cell dissociation. GBM cells were plated at 50E3 cells/well in a 96-well flat bottom plate. After acclimating overnight, cells were treated with tyrphostin9 at concentrations of 0.01, 0.1, 1, or 10uM or DMSO control. IFN-γ was then added to a final concentration of 1 ng/ml, and the cells were cultured for an additional 24 hours. Following this treatment, the medium containing drug and IFN-γ was removed and replaced with fresh medium, either without or with CD8⁺ NY-ESO-1 TCR-transduced T cells, at a GBM:T-cell ratio of 1:0.5. After 18 hours of co-culture, GBM nuclei were harvested, subjected to sci-Plex multiplexing, and sci-RNA-seq processing.

### Flow cytometry to measure HLA protein levels

BT112 or U87MG NY-ESO-1 cells were plated into 6-well dishes at a density of 2E5 cells/well and were acclimated overnight. Cells were exposed to DMSO control or 1uM of tyrphostin9, AG-555, or OSI-420. After 48 hours of chemical exposure, cells were removed from the plate with TrypLE (Gibco) and washed 2X with staining buffer (1X PBS + 0.5% w/v BSA). Cells were stained live with either a FITC conjugated mouse anti-human HLA-A2 antibody (BioLegend Cat# 343304) or FITC conjugated mouse anti-human B2M (pan-HLA, BioLegend Cat# 311404) at a dilution of 1:25 for 30 minutes in the dark at room temperature. The stained cells were washed 2X with staining buffer before flow cytometry. Fluorescence was measured on the Attune NxT Flow Cytometer, and the data was analyzed using FlowJo (version 10.9.0). Briefly, cells were gated to live, single cells and mean fluorescent intensity (MFI) for each treatment group was recorded.

### MrVI model training

MrVI^21^ is a deep generative model that performs sample stratification at single-cell resolution. As mentioned in the results, the model’s hierarchical probabilistic framework presumes cells to be generated from nested experimental designs, in which each sample is drawn from one of several experimental batches. MrVI learns two distinct latent feature spaces from a given scRNA-seq dataset, a sample-unaware *U*-space that is decoupled from the sample-of-origin and a sample-aware *Z*-space that incorporates sample-of-origin effects while accounting for technical factors across both spaces. These latent spaces can be used to dissect sample-specific heterogeneity in cell states across populations of interest in downstream analyses. The updated version of MrVI we applied features multi-head cross-attention-based decoders from the U-space to the Z-space and from the Z-space to the observed space for improved integration. Additionally, the U-space can be constrained to a lower dimensional space relative to the Z-space to further improve the mixing of different samples in the U-space.

We trained a MrVI model for each PDCL (i.e. BT112, BT228 and BT333) with the sample key defined as unique drug-dose combination and the batch key defined as replicate. For all trained models, we used the recommended default model arguments: *n_latent=30; n_latent_u=10; qz_nn_flavor=“attention”; px_nn_flavor=“attention”; use_map (qz_kwargs)=True; stop_gradients (qz_kwargs)=False; stop_gradients_mlp (qz_kwargs)=True; dropout_rate (qz_kwargs)=0.03; stop_gradients (px_kwargs)=False; stop_gradients_mlp (px_kwargs)=True; h_activation (px_kwargs)=“nn.softmax”; low_dim_batch (px_kwargs)=True; dropout_rate (px_kwargs)=0.03; learn_z_u_prior_scale=False; z_u_prior=True;* and *u_prior_mixture=False.* We used the following training arguments in tandem with the recommended default model arguments: *max_epochs=100; batch_size=256; early_stopping=True; early_stopping_patience=15; check_val_every_n_epoch=1; train_size=0.9; lr (pl_kwargs)=2e-3; n_epochs_kl_warmup (plan_kwargs)=20; max_norm (plan_kwargs)=40; eps (plan_kwargs)=1e-8;* and *weight_decay (plan_kwargs)=1e-8*. To select for gene features inputted into the MrVI models, we obtained the union of the 100 highly variable genes (determined in Scanpy) for each unique PDCL-drug data-subset then filtered for expression in at least 5% of cells included in the total dataset. To visualize the output of the MrVI models, we generated UMAPs from each the *U* and *Z* latent spaces. Specifically, we sought a single cluster with minimal substructure in the sample-unaware *U*-space that mapped to multiple clusters or blatant substructure in the sample-aware *Z*-space.

### Defining response modules

The functional relationship between the sample-unaware *U-space* and the sample-aware *Z-space* in MrVI can be used to directly estimate single-cell resolution sample-sample distance matrices. The sample-sample distances are obtained by computing the Euclidean distances between sample-specific counterfactual cell states. To avoid technical factors, counterfactual cell state predictions are made at the level of the Z latent space. In the context of our study, MrVI enabled us to relate the transcriptional effects between unique drug-dose combinations with respect to each PDCL. As we expect each PDCL to be relatively homogeneous in their response to the treatments, we take the average of the normalized distance matrices across all of the cells of each PDCL so our downstream analysis is robust to noise in individual distance matrix estimates. Hierarchical clustering of the mean sample-sample distances across cells resulted in distinct response modules (RMs) containing drug-dose combinations that induced similar transcriptional responses in a given PDCL. The hierarchical clustering depth was evaluated and determined by observation of silhouette plots.

### Covariate-specific DE analysis

To obtain a transcriptional signature for each RM we performed covariate-specific DE analysis for each PDCL, a procedure that leverages a fitted MrVI model. Along with updates to the model architecture, the updated version of MrVI includes local measures of differential expression and differential abundance at a single-cell resolution. In particular, the differential expression procedure takes in sample-specific covariates and compares the counterfactual Z latent vectors for a given U latent vector. By decoding the result of a covariate’s average impact in the Z latent space, MrVI can report log fold change (LFC) values for each gene. In simpler terms, the LFC values provide a gene-level characterization of each RM. In our analysis, we set DMSO-treated cells as the vehicle when measuring differential expression for each RM. The parameters we used to run the covariate-specific DE analysis function follow: *batch_size=32; use_vmap=True; normalize_design_matrix=True; add_batch_specific_offsets=False; mc_samples=100; store_lfc=True; store_lfc_metadata_subset=None; store_baseline=False; eps_lfc=1e-3; filter_donors=False; lambd=0.0;* and *delta=0.3*. To filter for genes included in each RM signature, we set a cutoff of absolute LFC value > 0.2 and Benjamin-Hochberg adjusted p-value < 0.05. Gene-level adjusted p-values are unobtainable from MrVI covariate-specific DE analysis. Hence, we utilized the p-values associated with general linear models (GLMs) fitted across all genes inputted into MrVI for each PDCL ^95^, performed with the R package *monocle3*’s *fit_model* function. In each GLM, RM and replicate are set as terms. As in the MrVI covariate-specific DE analysis, we set DMSO-treated cells to be vehicles. Resulting p-values were adjusted for multiple hypothesis correction with the Benjamini-Hochberg method.

### Defining response module clusters across PDCLs

We merged the MrVI covariate-specific DE analysis LFCs for RMs across all PDCLs. The associated genes were filtered to include strictly the union of significant DEGs each RM signature (FDR < 5% & LFC > 0.1). Hierarchical clustering of the normalized LFCs (z-scored for each gene) revealed RM clusters. The aim of hierarchical clustering here was to find drugs with similar transcriptional responses and normalizing LFCs helped relate RMs across PDCLs despite variation in the magnitude of their response to drug treatment. While the resulting RM clusters were largely representative of one PDCL, RMs with similar dynamics at the gene level were collapsed together indicating the RMs are representative of broader transcriptional programs.

### Defining Transcriptional Drug Classes (TDCs)

We plotted binary membership of each drug across the RM clusters as a heatmap (i.e. whether a drug is present in an RM cluster across any of its doses). Hierarchical clustering of the drugs based on their RM clusters membership enabled the formation of generalized transcriptional drug classes (TDCs). TDCs represent groups of EGFRi’s that induce a distinct transcriptional response across the PDCLs.

### Gene set enrichment analysis

For each response module, we filtered differentially expressed genes by FDR < 0.05 and an absolute LFC > 0.1. We performed gene set enrichment analysis (GSEA) separately for increasing and decreasing DEGs using the R package *piano*. Briefly, we utilized the function *runGSAhyper* to perform a Fisher’s exact test of the lists of genes against the gene sets “MSigDB Hallmark 2020”, “MSigDB Oncogenic Signatures’’, “Kinase Perturbations from GEO up” and “Kinase Perturbations from GEO down” against a background of expressed genes. Resulting p-values were adjusted for multiple hypothesis correction with the Benjamini-Hochberg method. Representative results are displayed in Figure 5 for conciseness, and results for RMs of each cell line can be found in Supp. Fig. 17. Of note, response modules for BT333 yielded no significant gene set enrichment results after multiple hypothesis correction, indicating collections of DEGs with little association with previously defined programs.

### Visualization of similarity of EGFRi transcriptional effects

Based on these TDCs, we generated lollipop plots representing the magnitude and heterogeneity of a drug’s expression of each response module (Fig. 4E, Supp. Fig. 18, Supp. Fig. 19). Briefly for a given drug, each response module’s respective up-regulated DE genes (identified by covariate-specific DE analysis) were aggregated, averaged, and log2 normalized to the mean of its correspondent DMSO control. In the lollipop plots, this quantity is represented by the size and color of each lollipop. Additionally, because aggregate score is calculated at the single-cell level, heterogeneity in response module expression could be characterized as percent cells within a drug’s high dose population with greater expression than DMSO control. This quantity is represented by the length of each lollipop. As a result, control condition lollipop plots for DMSO, media, and PBS appear largely monotone with even length. In these plots, the differences between TDCs is made more apparent. For instance, TDC5 contains only AZ5104, a demethylated metabolite of osimertinib which itself is part of TDC4. Although chemically the two compounds are almost identical, their drug class separation can be attributed to differences in response modules like BT333 RM9 and BT112 RM12 (Supp. Fig. 20).

## Data and code availability

Raw and processed data can be accessed and downloaded from NCBI GEO under accession number GSE261618. The code necessary to reproduce the analyses in this study can be found at github https://github.com/mcfaline-figueroa-lab/sci-Plex-EGFRi. The code to run MrVI can be found at https://github.com/YosefLab/mrvi under the “paper_reproducibility” tag.

## Acknowledgments

The authors would like to thank all members of the McFaline-Figueroa and Azizi labs for helpful suggestions and critical discussion on the study. The authors would like to thank Erin Bush and the JP Sulzberger Columbia Genome Center for next-generation sequencing support. The authors would also like to thank Neel Shah and lab as well as Carla Concepcion-Crisol and Yonghao Yu for helpful discussion. **Funding**: J.L.M.-F acknowledges support from grants from the NIH (R35HG011941) and the NSF (2146007). E.A. acknowledges support by grant number 2022-253560 from the Chan Zuckerberg Initiative DAF, an advised fund of Silicon Valley Community Foundation. These studies used the resources of the Cancer Center Flow Core Facility at Columbia University funded in part through Center Grant P30CA013696. **Author contributions**: J.L.M.-F conceived the project and provided overall supervision of the study; R.M.G. designed experiments; R.M.G. and N.H. analyzed and interpreted the data; R.M.G., N.H., A.W., J.H., L.S., M.V., H.F., N.O. , V.L., J.P.N. and S.W.M. performed experiments or aided in analysis. K.L.L. provided PDCL models. K.L.L., J.R.M-F, N.Y., and E.A. provided additional supervision. R.M.G., N.H., and J.L.M-F wrote the manuscript. All authors reviewed and contributed to the manuscript. **Competing Interests:** K.L.L. reports research support to DFCI from Bristol Meyers Squibb (BMS) and Eli Lilly, consulting and advisory roles to BMS, Travera, Blaze Bioscience, and Servier, and equity in Travera.

## Supplementary Figures

**Supplemental Figure 1.**
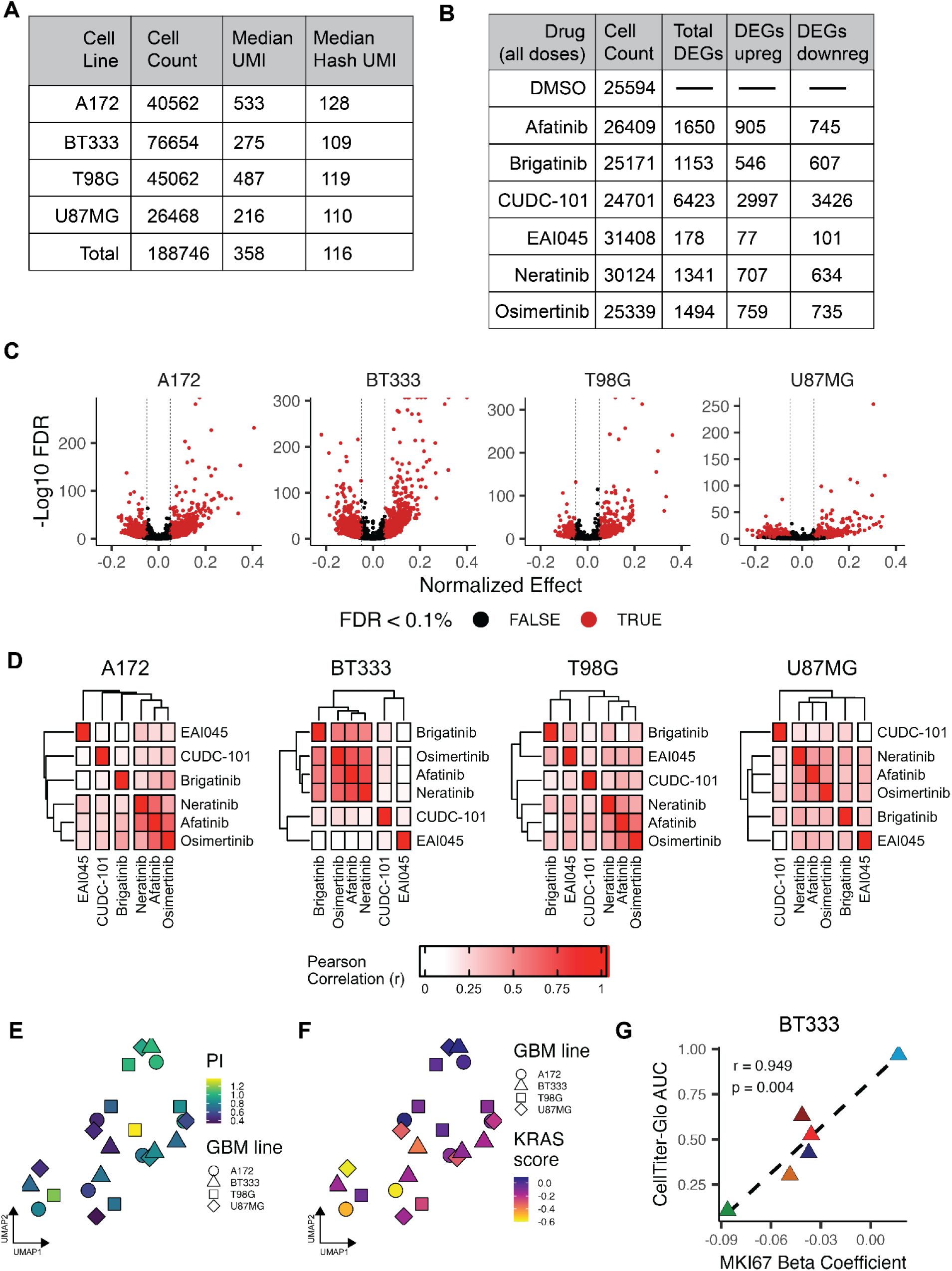
Pilot EGFRi screen QC and differential gene expression analysis. **A.** Table of experimental summary metrics by cell line. **B.** Table of experimental summary metrics by chemical agent. **C.** Volcano plots displaying the differential gene expression results for all drugs within a cell line. Only the terms from the contribution of dose as log10(dose + 0.01) from the quasi-poisson regression model fit to expression is shown. **D.** Pairwise Pearson correlation coefficients between drug’s normalized beta coefficients for significant genes (FDR < 0.1%) for each cell line. **E-F**. UMAP projections summarizing the correlation (Pearson’s coefficient of effects size estimates across the union of differentially expressed genes) between cell line-drug effects on transcription, colored by aggregate expression of proliferation index at 10uM (**E**) or KRAS signature at 10uM (**F**). **G**. Correlation between CellTiter-Glo-derived relative viability (AUC) and MKI67 beta coefficient for BT333.

**Supplemental Figure 2.**
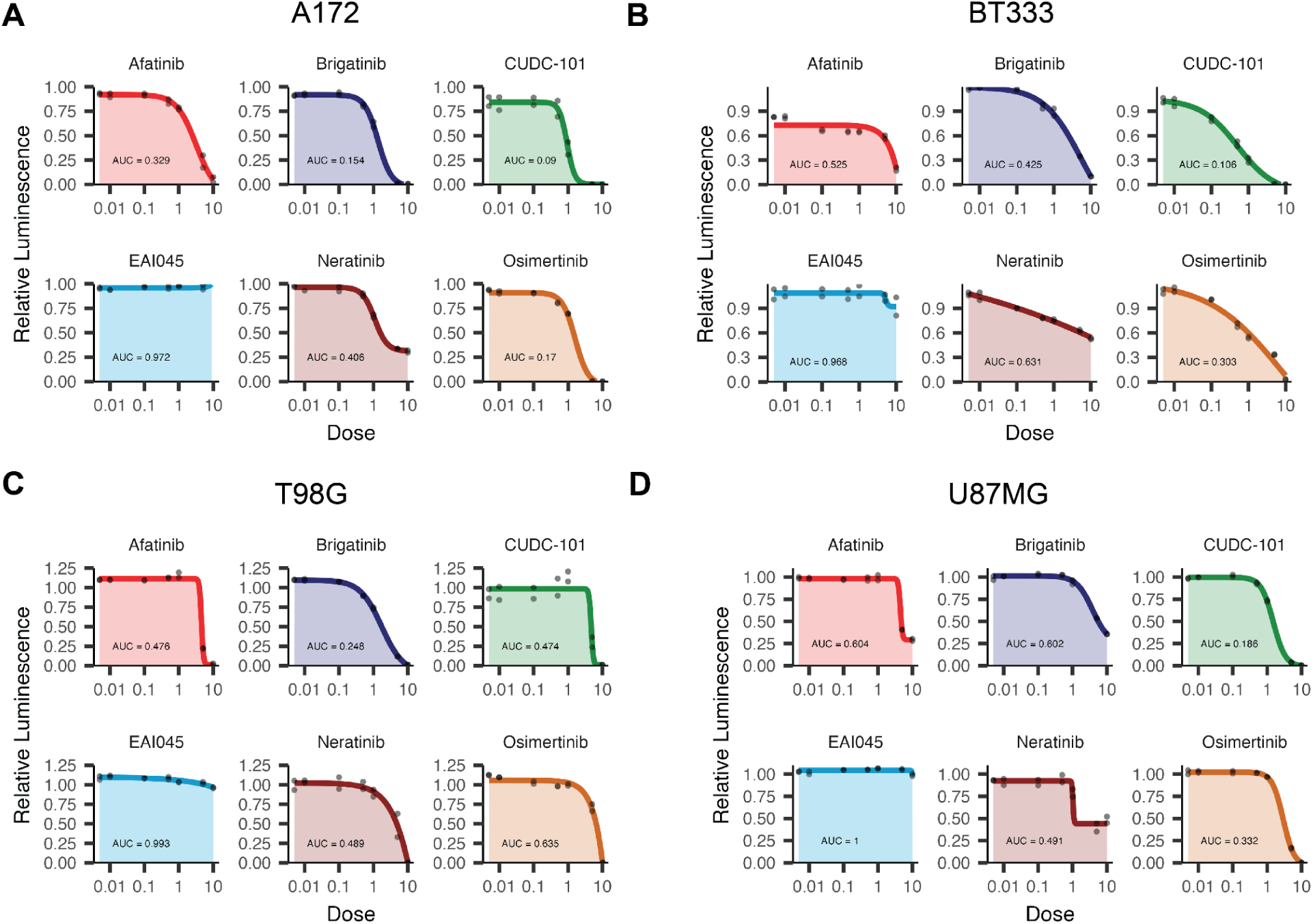
Pilot EGFRi screen orthogonal measure of GBM viability. **A-D**. Luminescence-based dose response and area under the curve (AUC) of the 6 pilot screen EGFR inhibitors for A172 (**A**), BT333 (**B**), T98G (**C**), and U87MG (**D**).

**Supplemental Figure 3.**
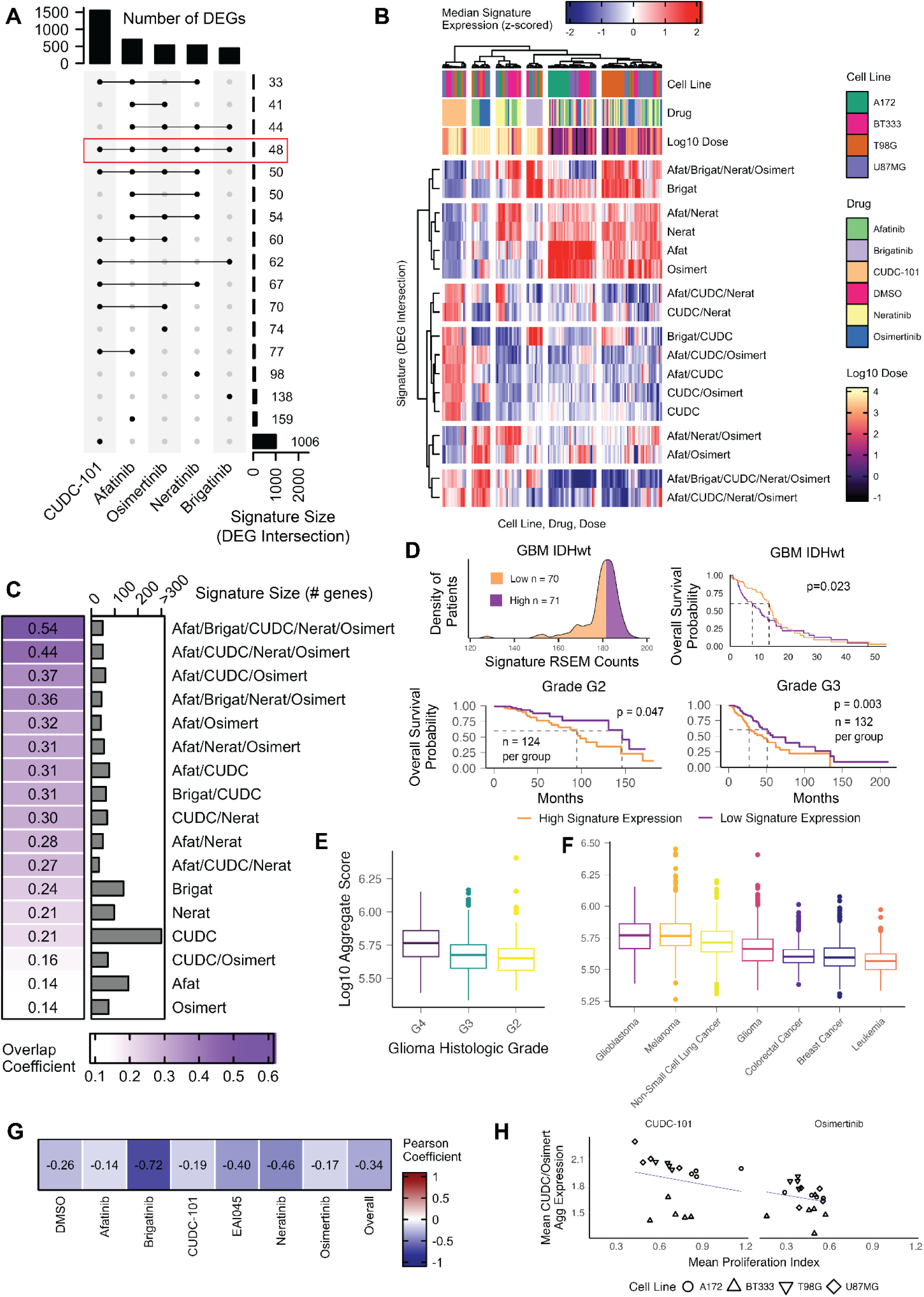
Defining chemical signatures from DEGs in the pilot EGFRi screen. **A.** Upset plot displaying the unique intersections (non-overlapping) of DEGs between the five drugs that induced substantial transcriptional changes. DEGs were limited to those upregulated in at least one cell line (normalized beta coefficient > 0.05, FDR < 0.1%). Highlighted by a red box is the 48 gene signature shared by all five inhibitors. **B.** Heatmap of z-scored aggregate signature scores by cell line, drug, dose conditions. Hierarchical clustering of the columns shows that the defined signatures are representative shared drug effects across cell lines. Hierarchical clustering of the rows illustrates that the shared five inhibitor signature is transcriptionally distinct from the other defined signatures. **C.** Overlap of pilot EGFRi signature genes with a recently defined adaptive resistance signature across established GBM cell lines and glioma stem cell models in response to the inhibition of RTK signaling ^22^. Note that overlap does not correlate with signature size. **D.** Stratification of IDHwt GBM patients from TCGA by top or bottom 50% of aggregate expression of the shared EGFR inhibitor transcriptional signature. Overall survival of IDHwt GBM patients or glioma patients stratified by the shared EGFR inhibitor transcriptional module expression (p: Gehan-Wilcoxon test). **E.** Aggregate expression distribution of the five inhibitor shared signature by glioma histologic grade from TCGA patient samples, arranged by highest median aggregate expression *(left to right)*. **F.** Aggregate expression distribution of the five inhibitor shared signature by major cancer types from PanCancer TCGA patient samples, arranged by highest median aggregate expression *(left to right)*. **G.** Heatmap of Pearson correlation coefficients between mean CUDC-101/Osimertinib signature aggregate score and mean proliferation index score, for EGFRi (10uM) pseudobulked by cell line and well replicate. The module is weakly anti-correlated with viability as measured by proliferation. **H.** Example of correlation between CUDC-101/osimertinib signature mean aggregate score and mean proliferation index score, for cells treated with 10uM CUDC-101 (*left*) and osimertinib (*right*).

**Supplemental Figure 4.**
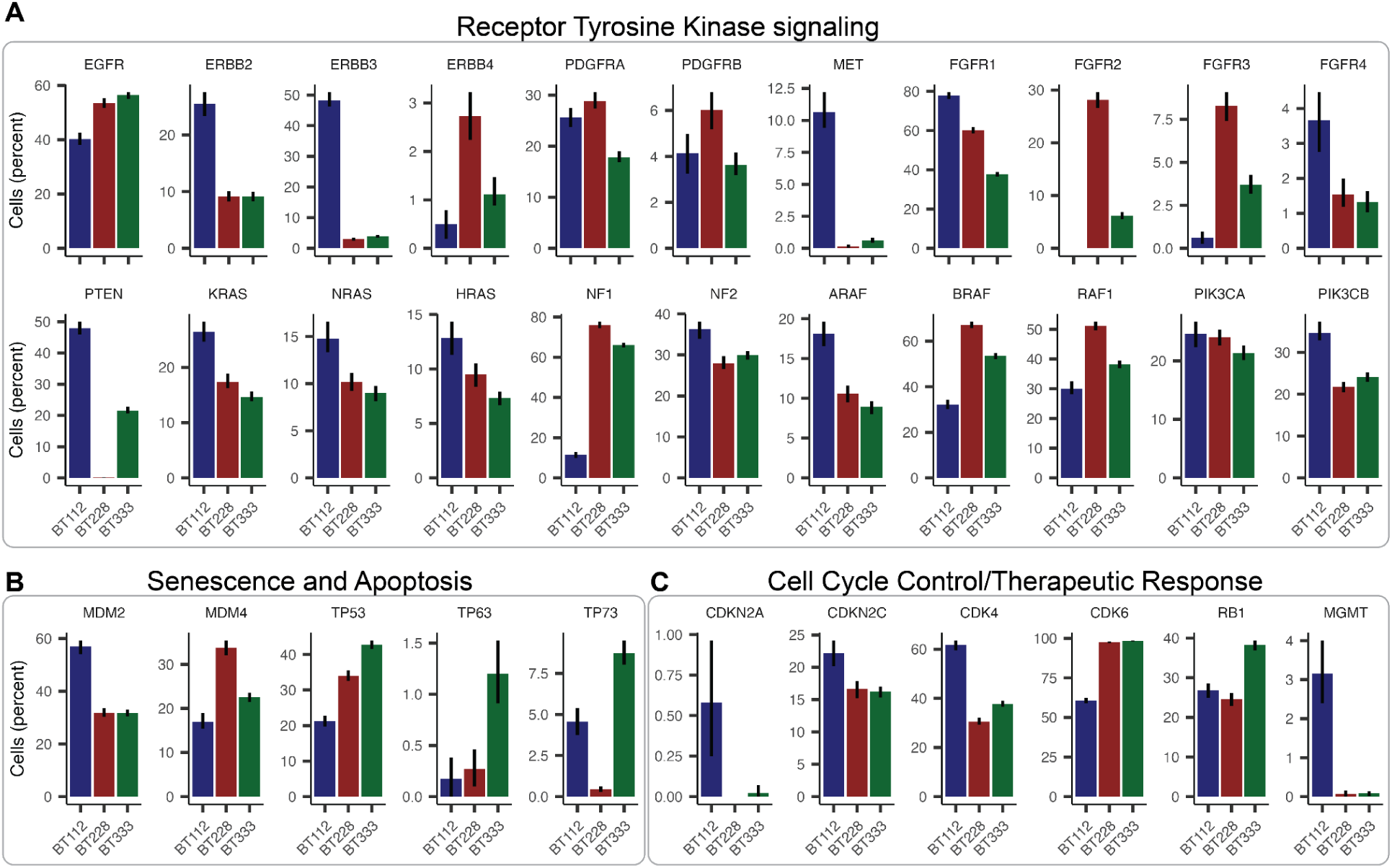
Inter-model variation in RTK pathway expression across patient-derived glioblastoma models. **A**. Expression of genes associated with receptor tyrosine kinase and EGFR-specific signaling across BT112, BT228, and BT333 patient-derived models (PDCLs). **B**. Expression of genes associated with senescence and apoptosis across PDCLs. **C**. Expression of genes associated with cell cycle control and therapeutic response across PDCLs.

**Supplemental Figure 5.**
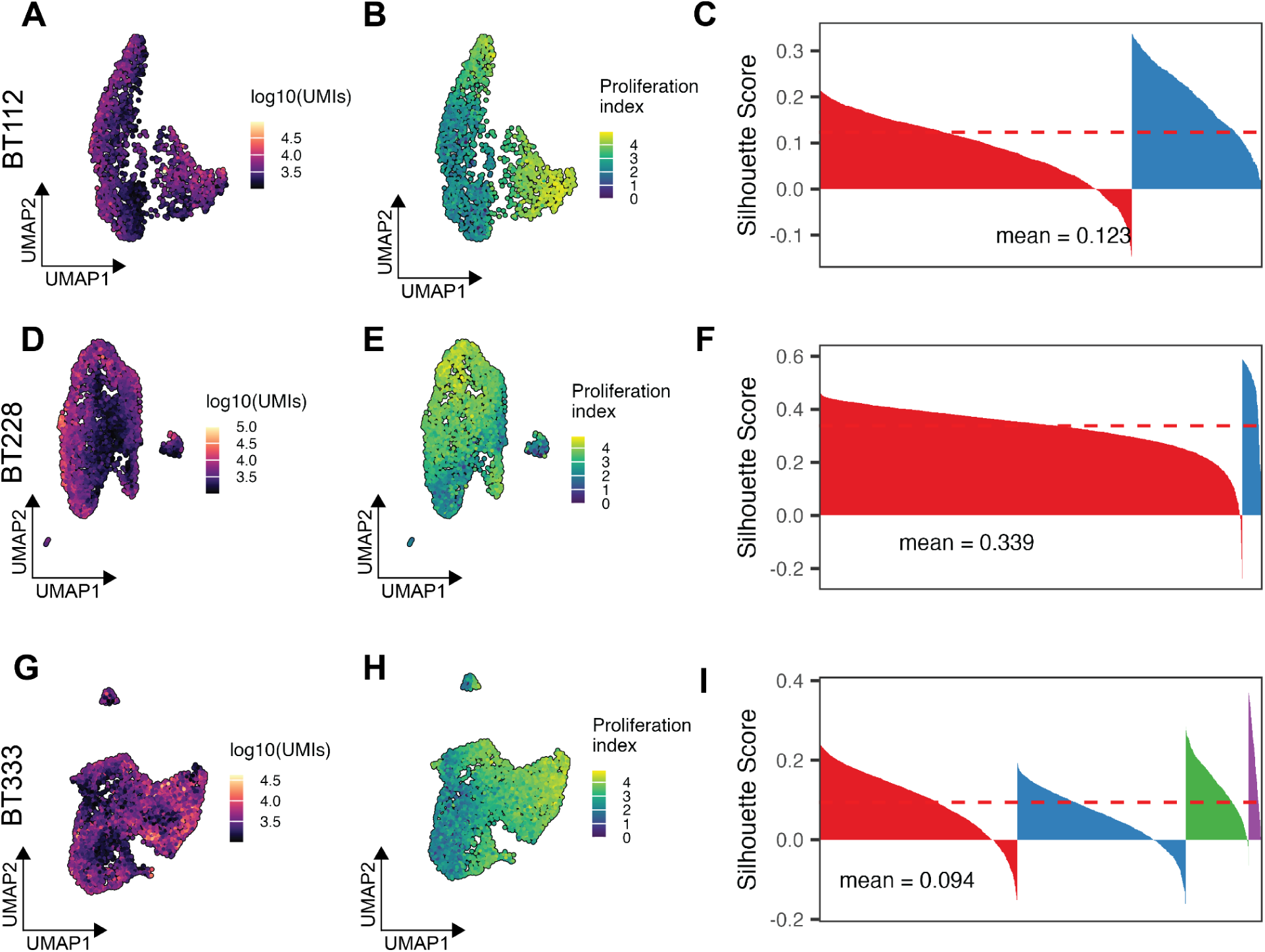
Intra-model variation across patient-derived glioblastoma models. **A-B**. UMAP embeddings of parental BT112 colored by total number of transcripts (UMIs) captured per cell (**A**) or the aggregate expression of genes associated with proliferation (**B**)**. C**. BT112 silhouette scores for Leiden clusters. **D-E**. UMAP embeddings of parental BT228 colored by total number of transcripts (UMIs) captured per cell (**D**) or the aggregate expression of genes associated with proliferation (**E**)**. F**. BT228 silhouette scores for Leiden clusters. **G-H**. UMAP embeddings of parental BT333 colored by total number of transcripts (UMIs) captured per cell (**G**) or the aggregate expression of genes associated with proliferation (**H**)**. I**. BT333 silhouette scores for Leiden clusters.

**Supplemental Figure 6.**
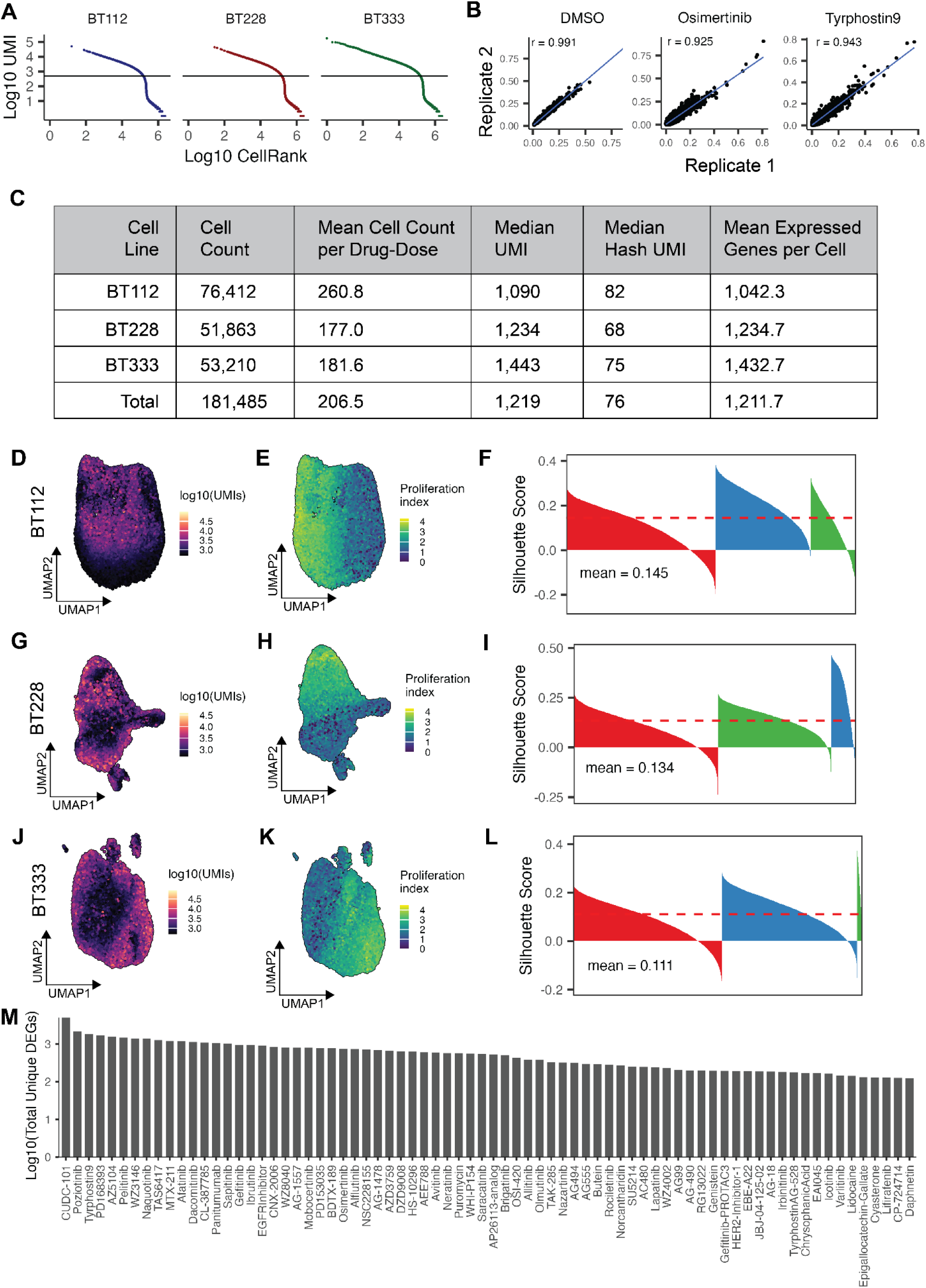
Comprehensive screen of EGFRi in patient-derived models of GBM. **A.** Kneeplots for each PDCL as UMI vs cell rank with a cut-off at 500 UMI. **B.** The normalized gene expression of replicates of the screen is highly correlated across chemical agents (DMSO, 10uM Osimertinib, 10uM Tyrphostin9 from left to right). **C.** Table of experimental summary metrics overall or by cell line. **D-E**. UMAP embeddings of 24 hour exposure BT112 cells colored by total number of transcripts (UMIs) captured per cell (**D**) or the aggregate expression of genes associated with proliferation (**E**)**. F**. 24 hour exposure BT112 silhouette scores for Leiden clusters. **G-H**. UMAP embeddings of 24 hour exposure BT228 cells colored by total number of transcripts (UMIs) captured per cell (**G**) or the aggregate expression of genes associated with proliferation (**H**)**. I**. 24 hour exposure BT228 silhouette scores for Leiden clusters. **J-K**. UMAP embeddings of 24 hour exposure BT333 cells colored by total number of transcripts (UMIs) captured per cell (**J**) or the aggregate expression of genes associated with proliferation (**K**)**. L**. 24 hour exposure BT333 silhouette scores for Leiden clusters. **M.** Dose-dependent differentially expressed genes per compound determined by significant coefficients from fitting a quasipoisson regression model, ordered by decreasing count.

**Supplemental Figure 7.**
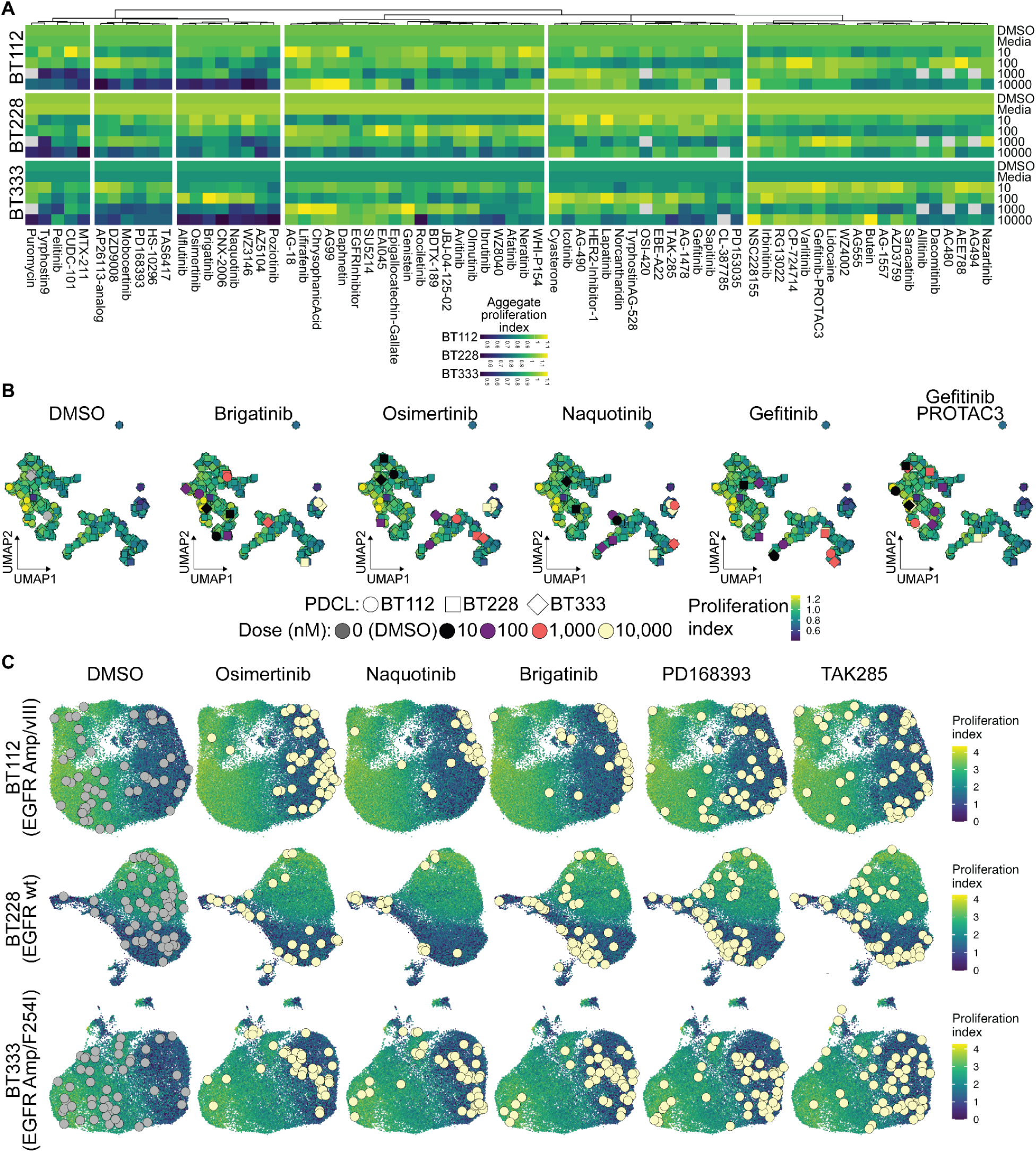
EGFRi effects on proliferative gene expression. **A.** Heatmap of mean aggregate proliferation score as for each PDCL, drug, dose, as in Fig. 3A. **B.** UMAP projection of EGFRi PDCL, drug, dose points represented by features as log2 normalized expression to DMSO control. Mean aggregate proliferation index is represented by the underlying viridis color scale, and select drugs are faceted and colored to highlight the differential effects of each drug-dose combination. **C.** UMAP projection of single-cells treated with EGFRi embedded individually for each PDCL. Mean aggregate proliferation index is represented by the underlying viridis color scale, and a sample of cells that were exposed to 10uM EGFRi are faceted and colored to highlight differential effects.

**Supplemental Figure 8.**
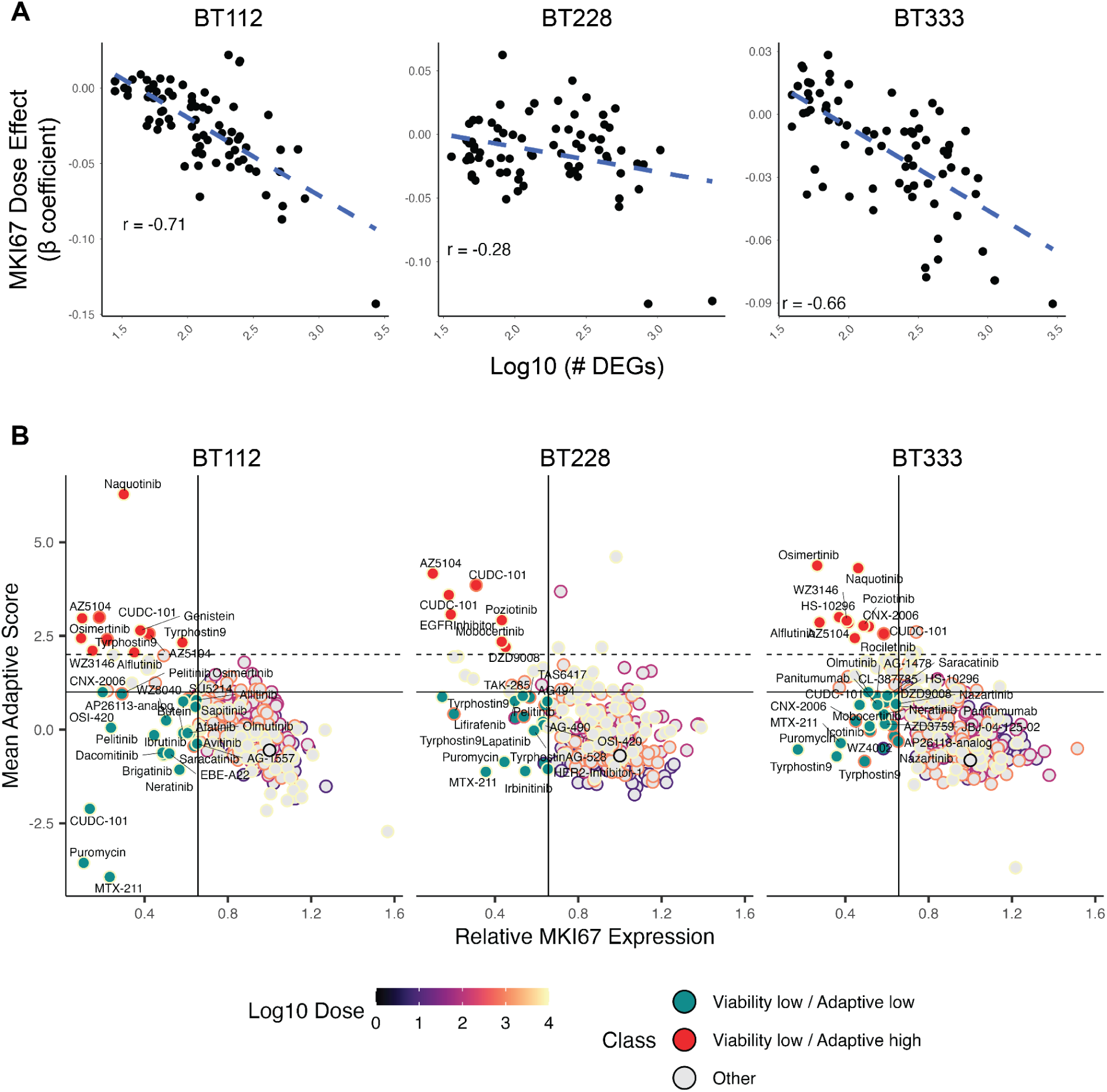
Correlation between MKI67 expression and global transcriptional impact or adaptive resistance signaling. **A.** MKI67 beta coefficient has a varying correlation with global transcriptional impact for BT112, BT228, and BT333 (left to right). The scatter plots represent the Pearson correlation within a cell line of inhibitor impact on MKI67 (normalized beta coefficient from quasi-Poisson regression of MKI67 as a function of dose) versus overall transcriptional changes (as total number of DEGs). **B.** Drug-dose group mean adaptive score versus mean MKI67 expression for BT112, BT228, and BT333 (left to right) reveals MTX-211 as an agent that alters viability while minimizing an adaptive response. Outside color of points denotes the dose while the fill color of points designates the ability to alter viability and adaptive signatures above or below the mean + sd or mean + 2sd of all drug-dose groups.

**Supplemental Figure 9.**
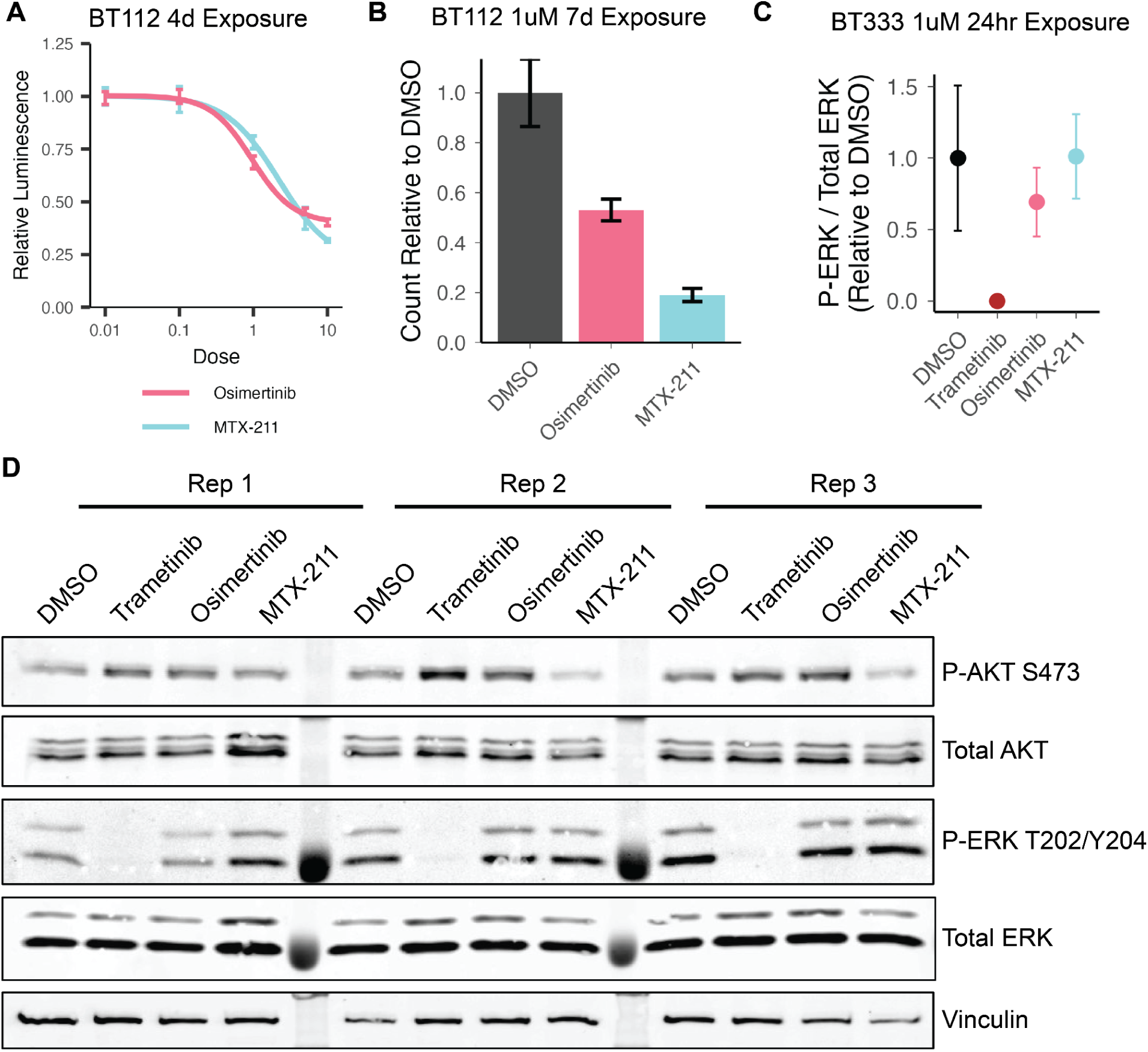
Induction of transcriptional adaptive resistance correlates with effects on proliferation and downstream MAPK protein phosphorylation levels. **A**. Relative cell counts of BT112 cells treated with 1uM exposures of the indicated inhibitors for 7 days. **B**. Relative phosphorylated ERK1/2 levels for BT333 cells treated with 1uM exposures of the indicated inhibitors for 24 hours, quantified from Western blots of 3 replicates. **C**. Western blots of BT333 cells treated with 1uM exposures of the indicated inhibitors for 24 hours.

**Supplemental Figure 10.**
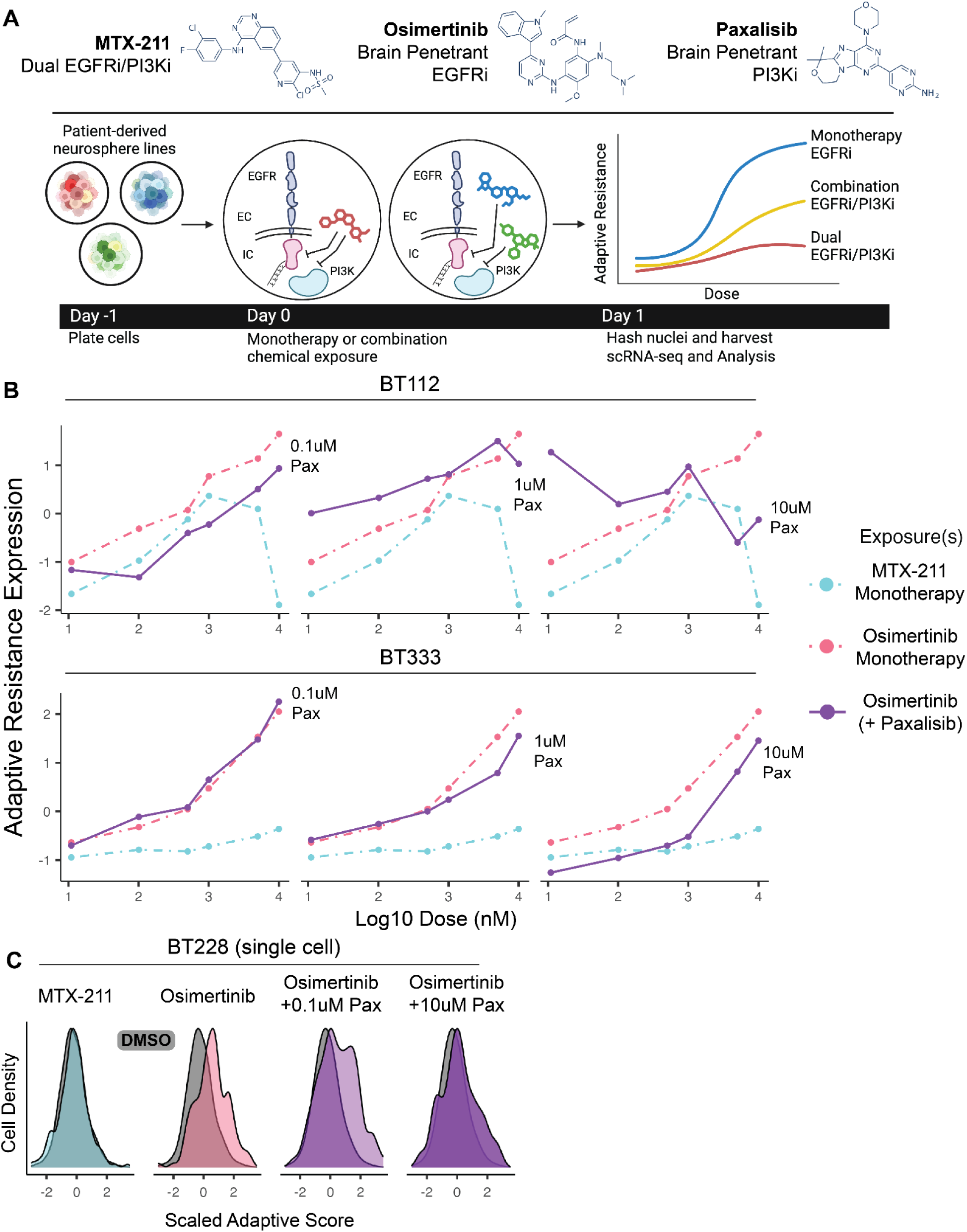
Combination exposure with clinically-relevant brain penetrant EGFRi and PI3Ki partially phenocopies dual inhibitor properties. **A**. Overview of our approach to compare adaptive resistance in combination osimertinib and paxalisib to the dual EGFRi/PI3Ki MTX-211 **B**. Adaptive resistance score as a function of drug dose for BT112 (top) and BT333 (bottom). Dotted lines represent monotherapy of MTX-211 and osimertinib and solid lines represent osimertinib at a specified constant paxalisib concentration. Paxalisib lowers the induction of adaptive resistance. **C**. Distributions of single-cell adaptive resistance scores for BT228 exposed to DMSO or a 10 uM concentration of the specified exposure (as in **B**) as monotherapy or combination.

**Supplemental Figure 11.**
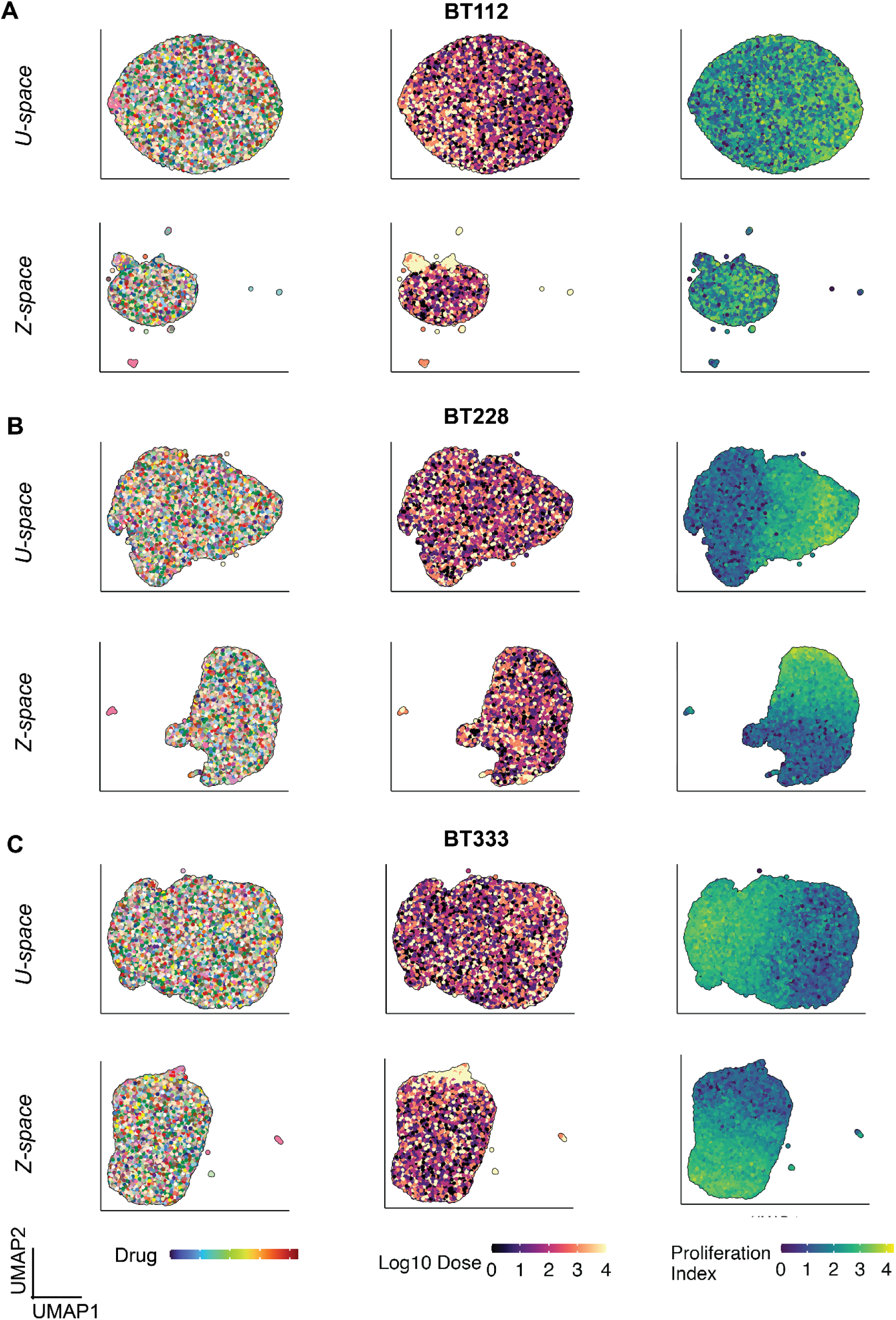
MrVI learned sample-unare and aware latent embeddings. UMAP dimensionality reduction is performed on *U-space* (top) and *Z-space* (bottom) latent embeddings, resulting in two-dimensional representations of **(A)** BT112, **(B)** BT228, and **(C)** BT333. Note that in the UMAPs of the Z-space, the largest clusters consist largely of cells treated with low doses or compounds that do not have a substantial molecular effect.

**Supplemental Figure 12.**
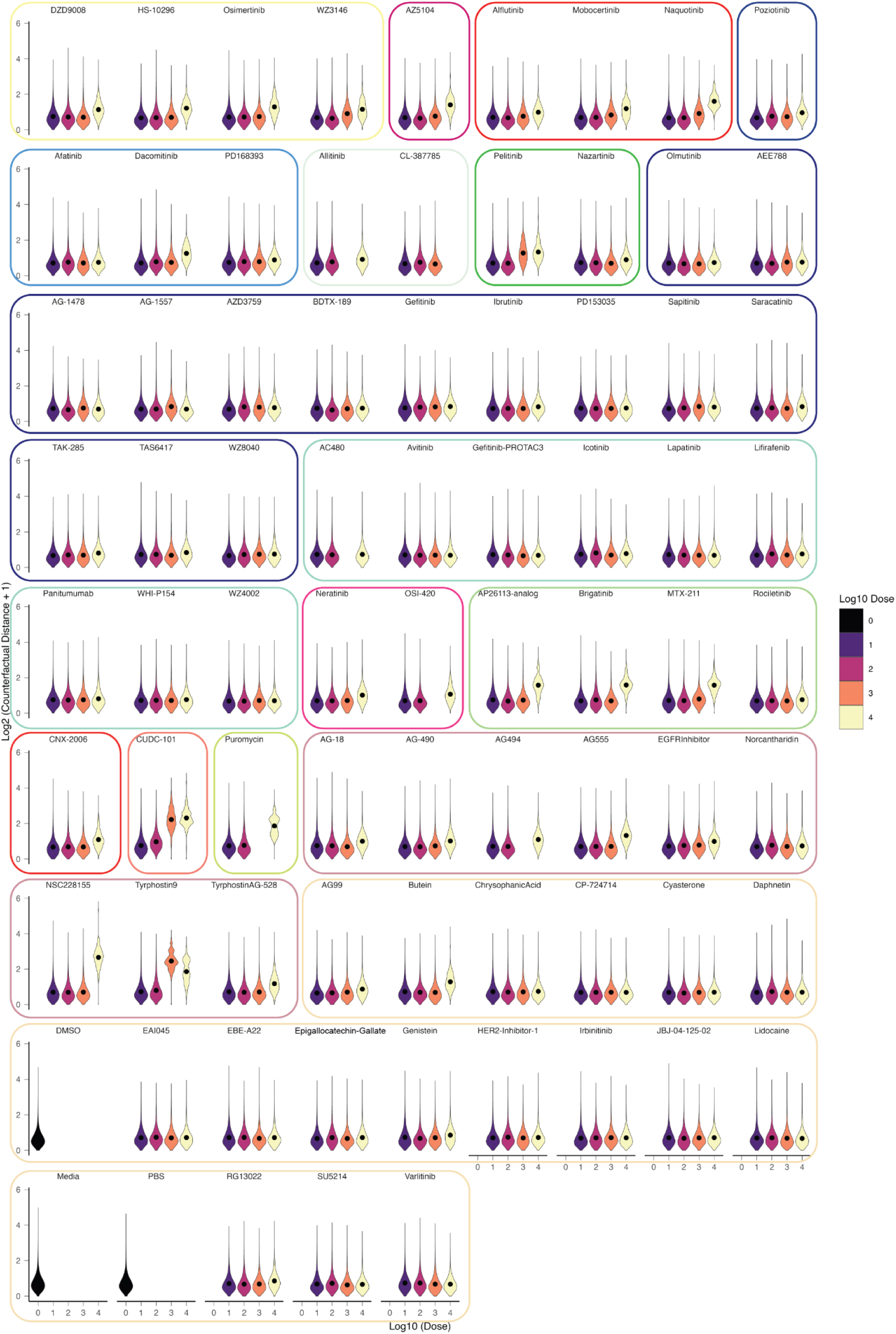
Distribution of counterfactual sample-sample distances across drug-dose conditions for BT112. The sample-sample distances from a given cell’s drug-dose condition to all other drug-dose conditions, including its own, are included as individual points in the violin plots. Hence, the number of cells x the number of drug-dose conditions data points are utilized. These plots exhibit the single-cell heterogeneity of drug-dose-specific induced transcriptional responses uncovered by MrVI.

**Supplemental Figure 13.**
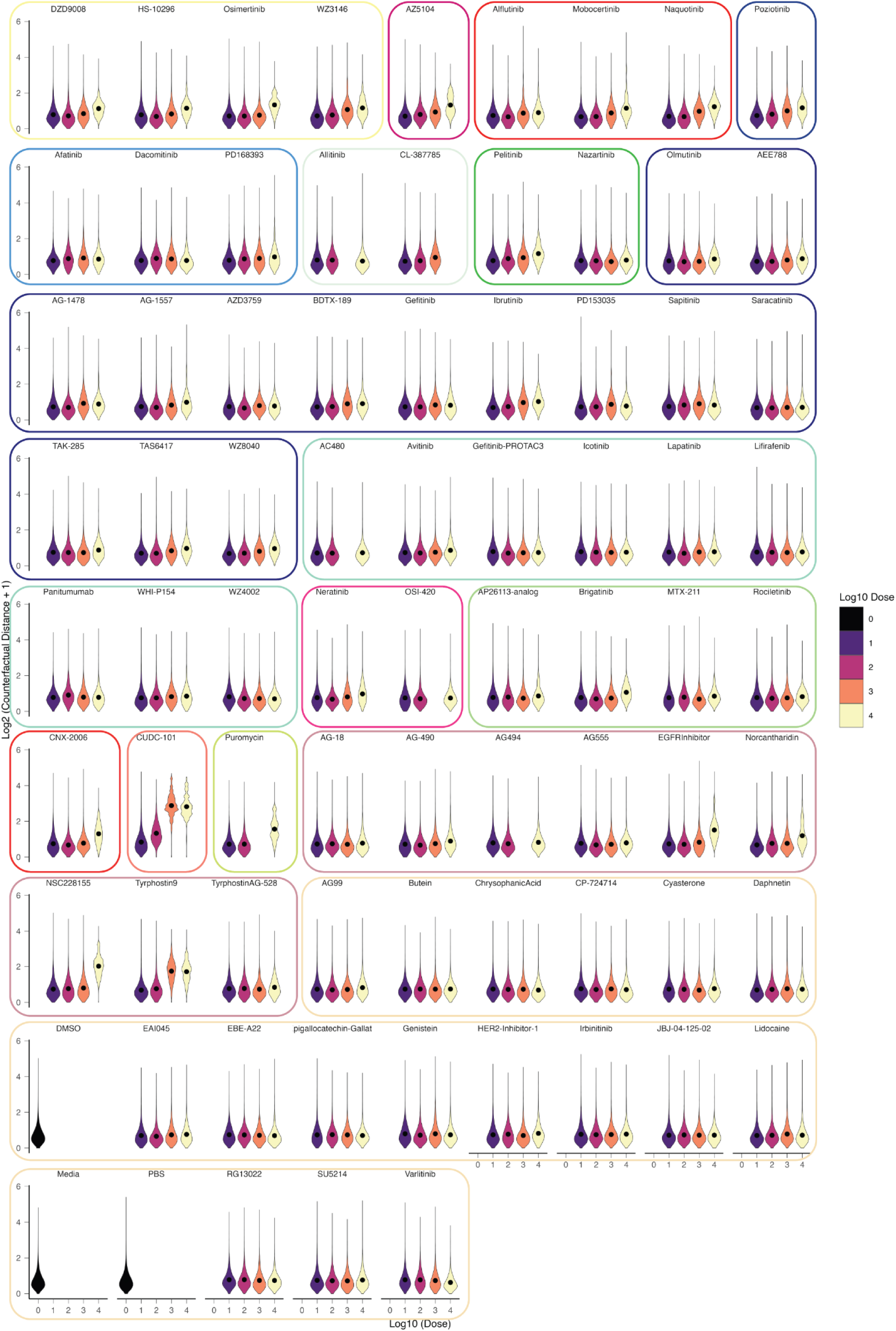
Distribution of counterfactual sample-sample distances across drug-dose conditions for BT228. The sample-sample distances from a given cell’s drug-dose condition to all other drug-dose conditions, including its own, are included as individual points in the violin plots. Hence, the number of cells x the number of drug-dose conditions data points are utilized. These plots exhibit the single-cell heterogeneity of drug-dose-specific induced transcriptional responses uncovered by MrVI.

**Supplemental Figure 14.**
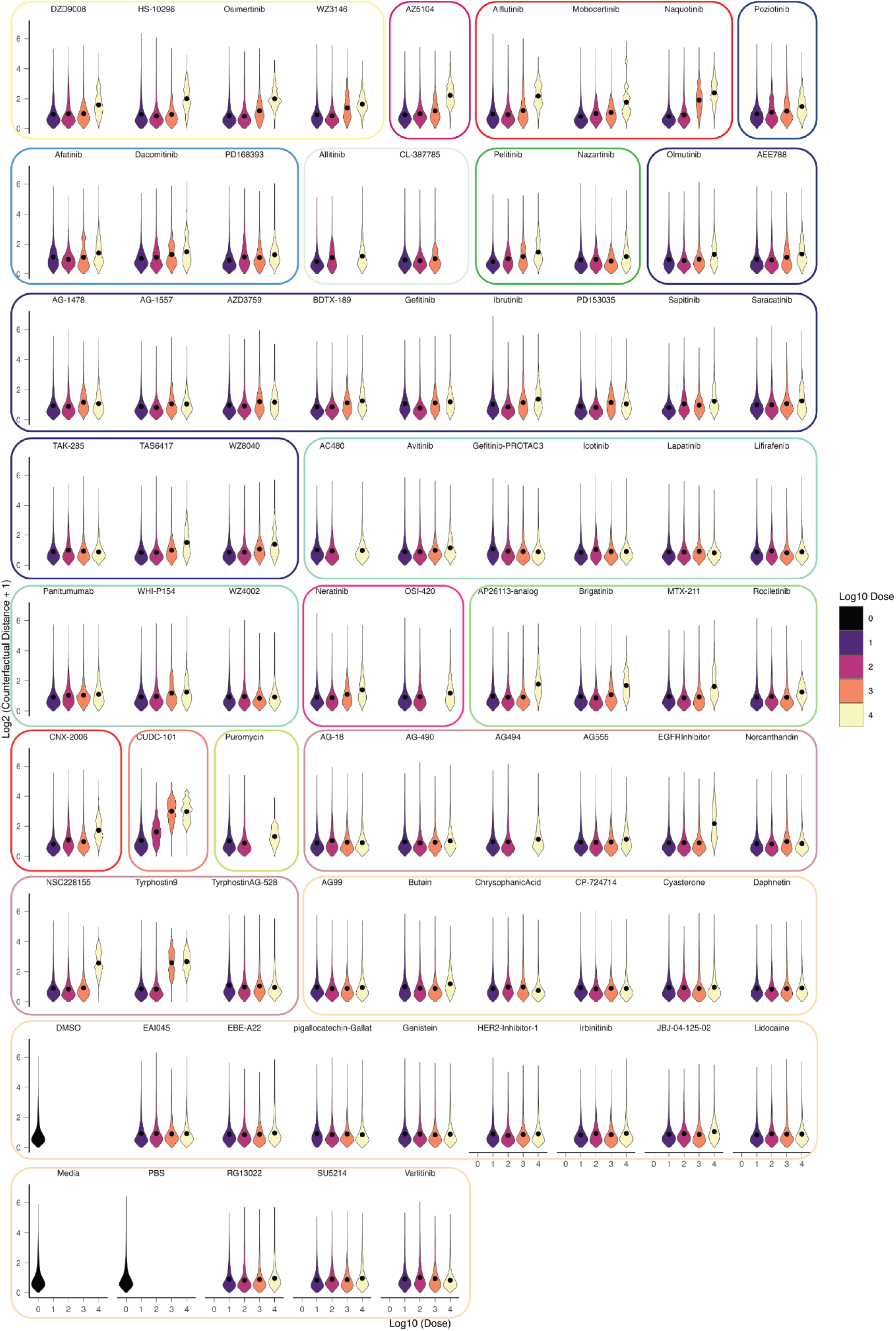
Distribution of counterfactual sample-sample distances across drug-dose conditions for BT333. The sample-sample distances from a given cell’s drug-dose condition to all other drug-dose conditions, including its own, are included as individual points in the violin plots. Hence, the number of cells x the number of drug-dose conditions data points are utilized. These plots exhibit the single-cell heterogeneity of drug-dose-specific induced transcriptional responses uncovered by MrVI.

**Supplemental Figure 15.**
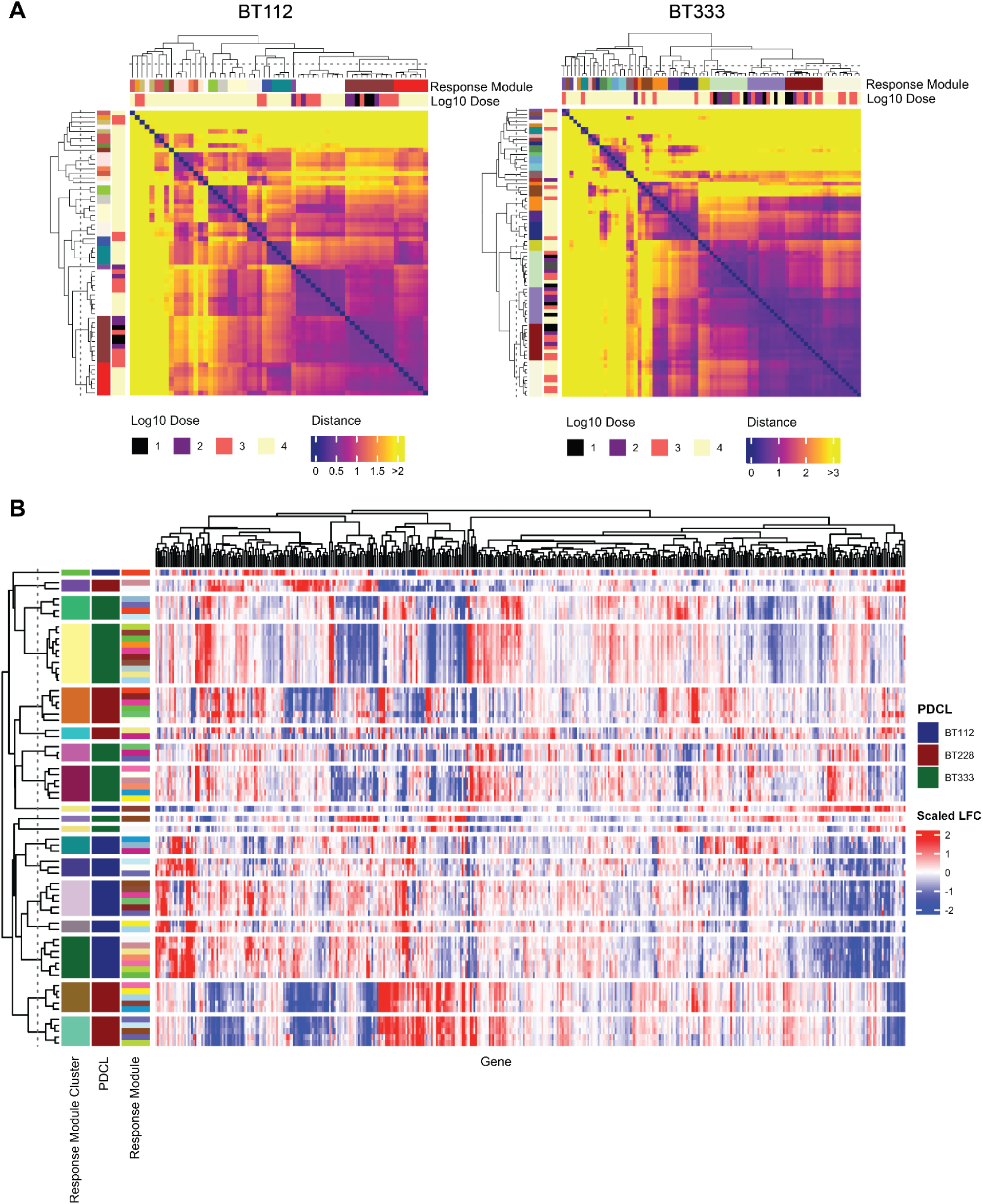
Sample-sample distances represent transcriptional similarity between drug-dose conditions. Sample-sample distances for a given cell are more specifically computed by calculating the Euclidean distances between sample-specific counterfactual cell states. Counterfactual predictions are estimated by leveraging the functional relationship between the sample-unaware U-space and sample-aware Z-space. **A.** Mean sample-sample distances across cells shown for BT112 and BT333, with the number of drug-dose conditions sampled for display. To sample, at most 10 random drug-dose conditions were selected from each response module. **B.** Log fold-changes (LFCs) across differentially expressed genes for each response module, as determined by covariate-specific differential expression analysis. Genes were considered differentially expressed if they had a LFC > 0.1 and an FDR < 5%. FDR values were obtained with pseudo-bulk general linear models.

**Supplemental Figure 16.**
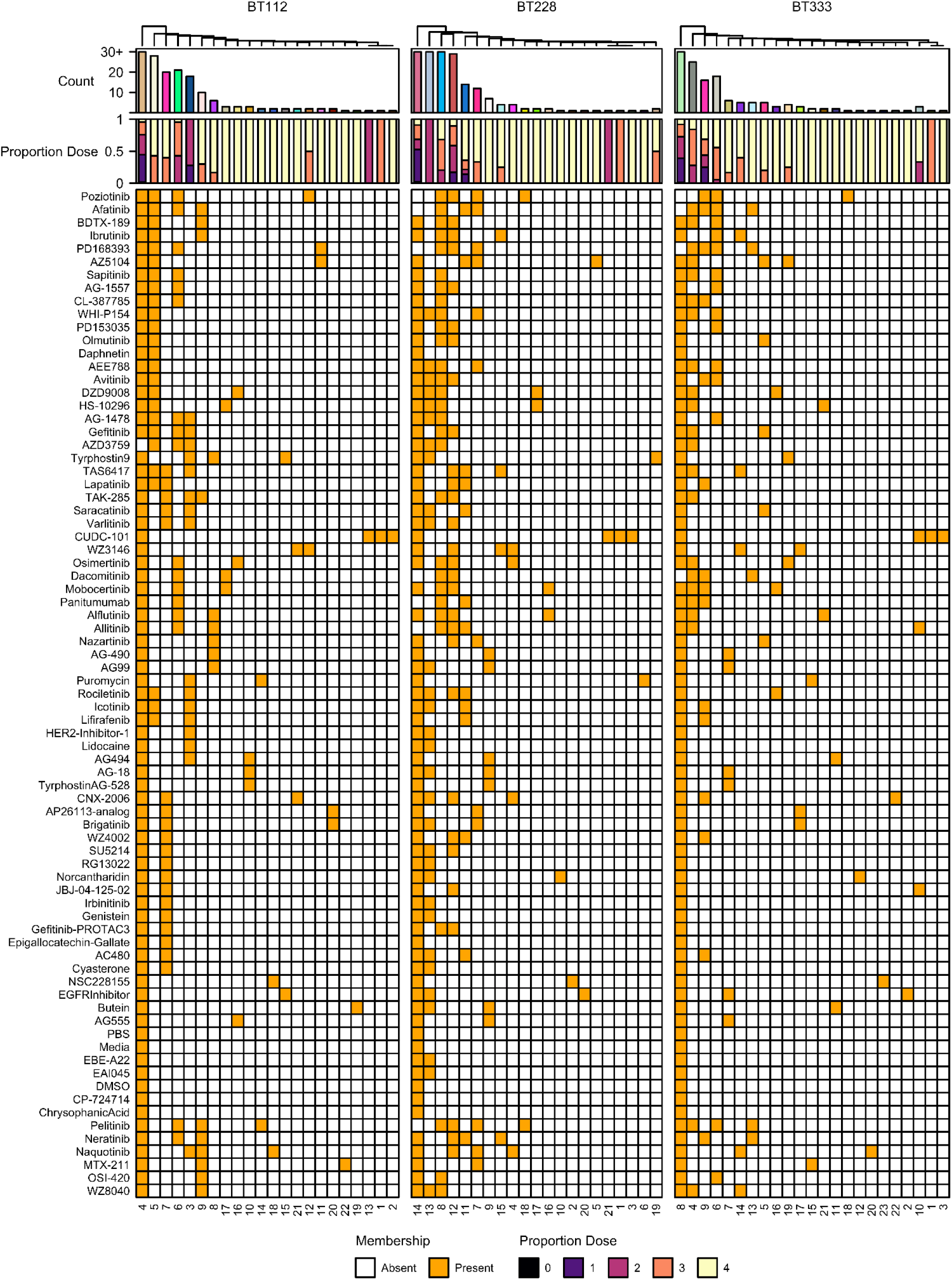
Drug-dose membership across response modules for each patient-derived model. The response module axes belonging to each PDCL are hierarchically clustered individually, and the drugs axis for each PDCL is clustered according to the hierarchical clustering of the BT112 drug axis. Here, drugs are shown to be similarly grouped across cell lines in accordance to their ability to induce specific transcriptional responses.

**Supplemental Figure 17.**
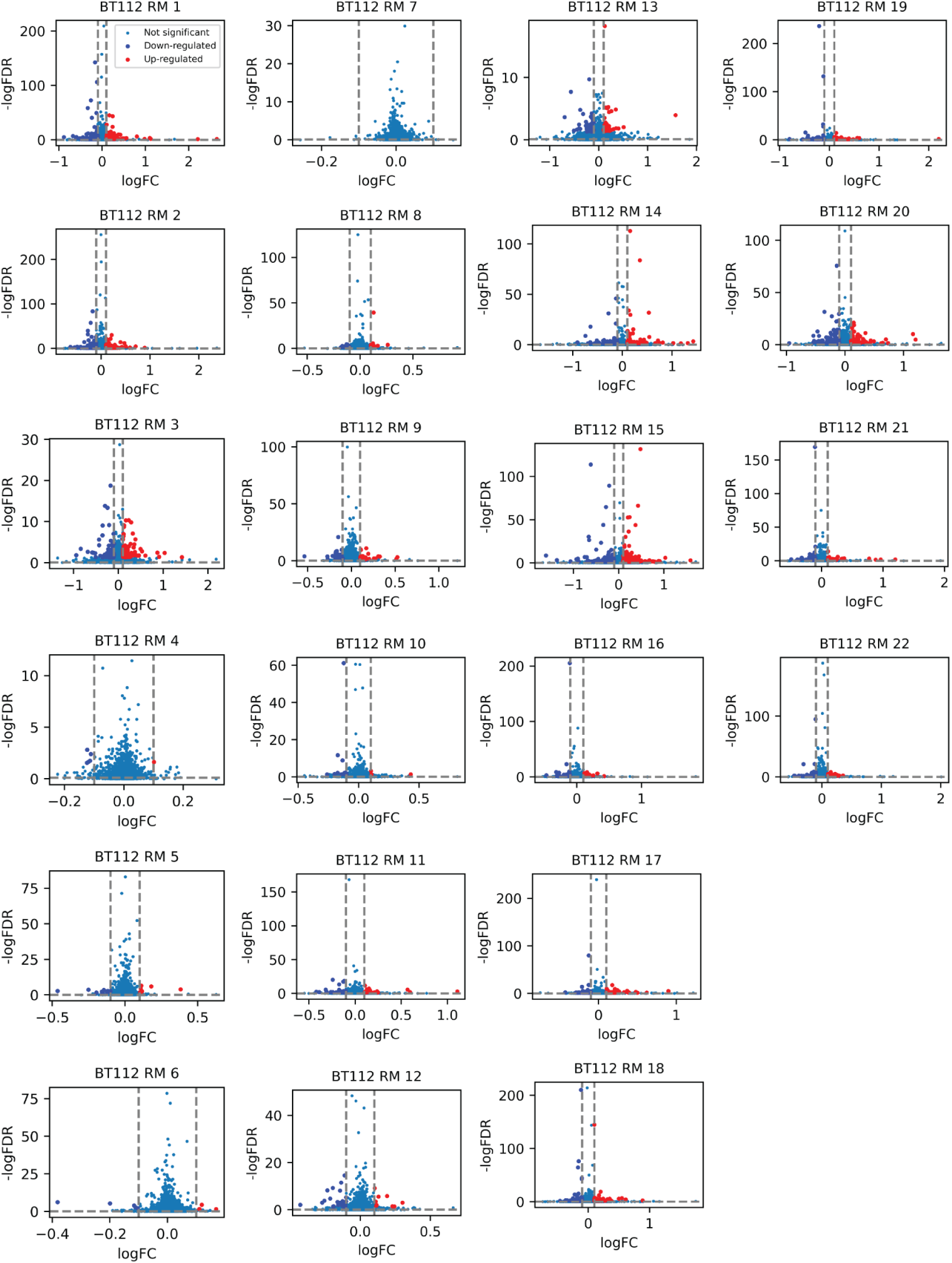
Differentially expressed genes of response modules (RMs) for BT112. The logFCs (threshold = 0.1) were obtained with MrVI multivariate analysis and the FDR-adjusted p-values (threshold = 0.05) were obtained using a quasi-poisson regression model.

**Supplemental Figure 18.**
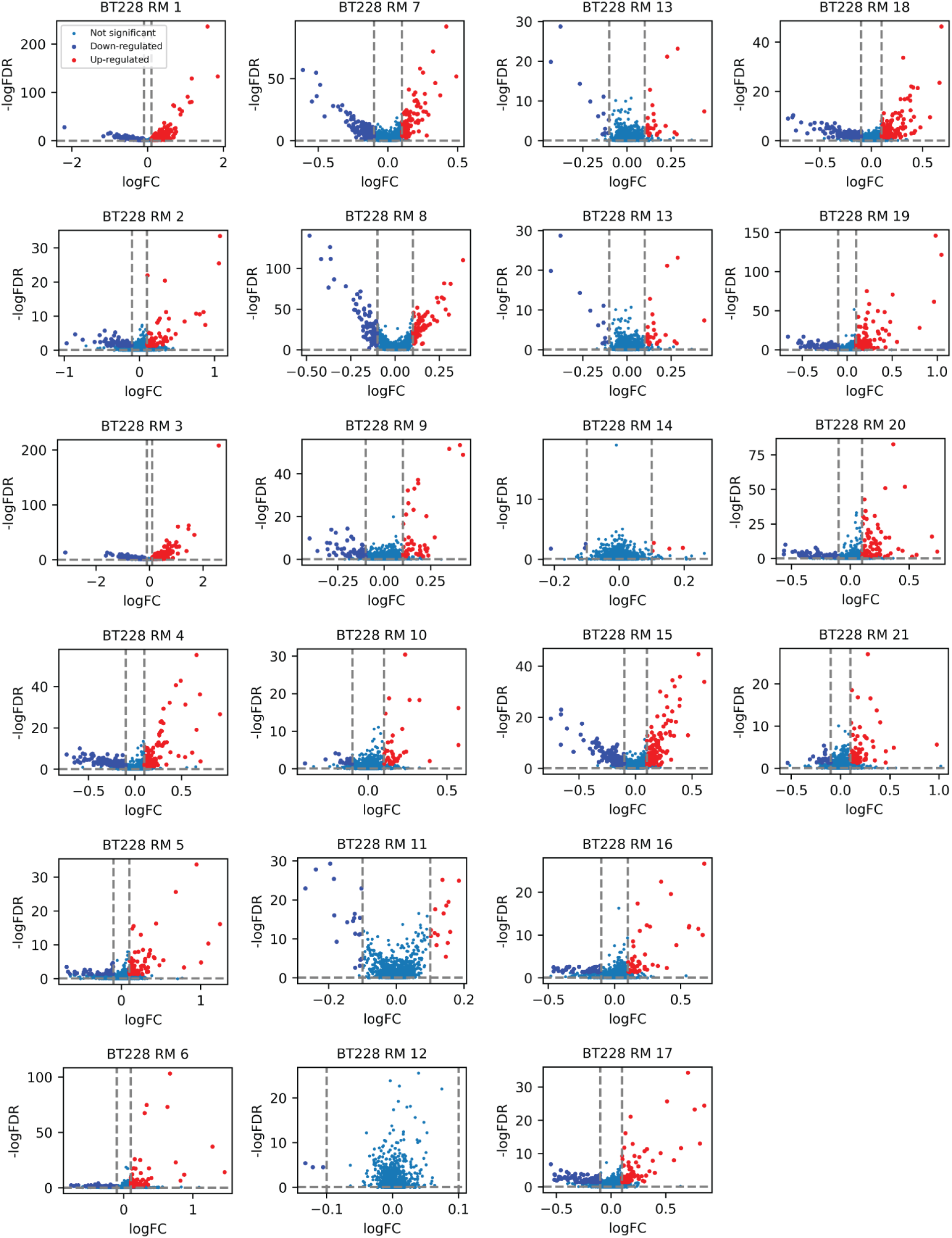
Differentially expressed genes of response modules (RMs) for BT228. The logFCs (threshold = 0.1) were obtained with MrVI multivariate analysis and the FDR-adjusted p-values (threshold = 0.05) were obtained using a quasi-poisson regression model

**Supplemental Figure 19.**
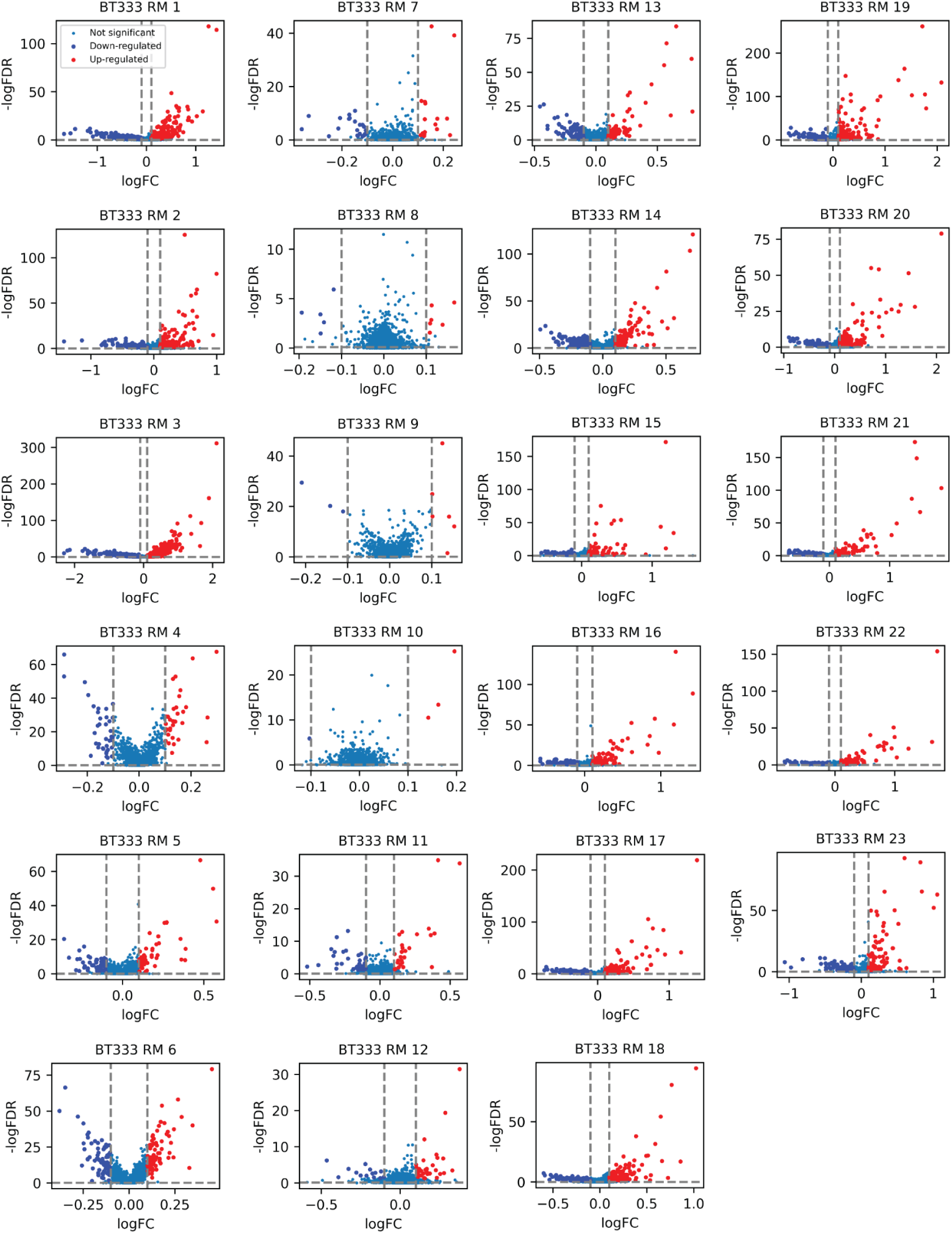
Differentially expressed genes of response modules (RMs) for BT333. The logFCs (threshold = 0.1) were obtained with MrVI multivariate analysis and the FDR-adjusted p-values (threshold = 0.05) were obtained using a quasi-poisson regression model.

**Supplemental Figure 20.**
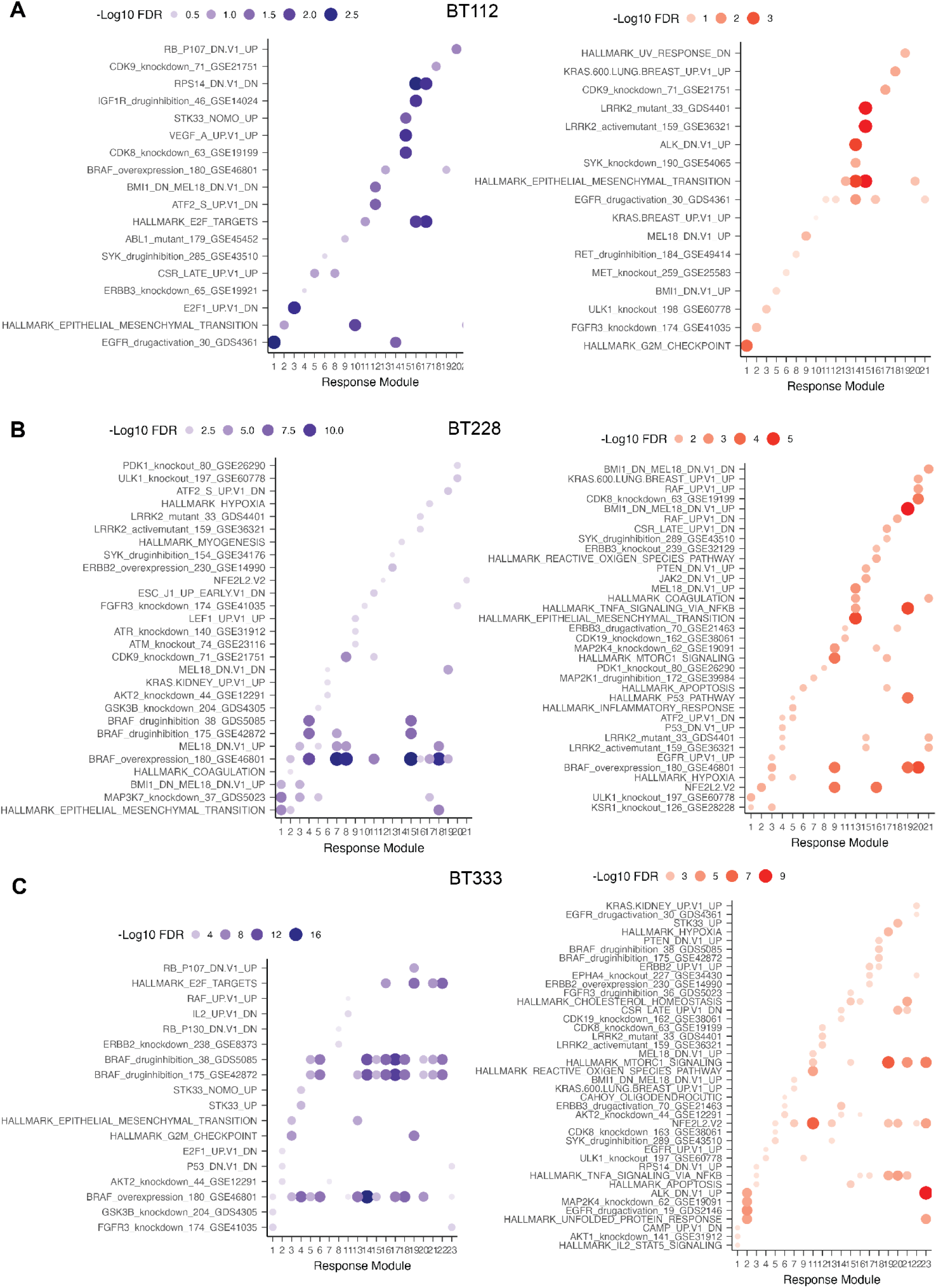
Gene-set enrichment analysis of upregulated and downregulated response module differentially expressed genes for (A) BT112, (B) BT228, **(C)** BT333. Of note, several BT112 RMs were not significantly enriched for the tested gene sets after multiple hypothesis correction, but top gene sets are displayed.

**Supplemental Figure 21.**
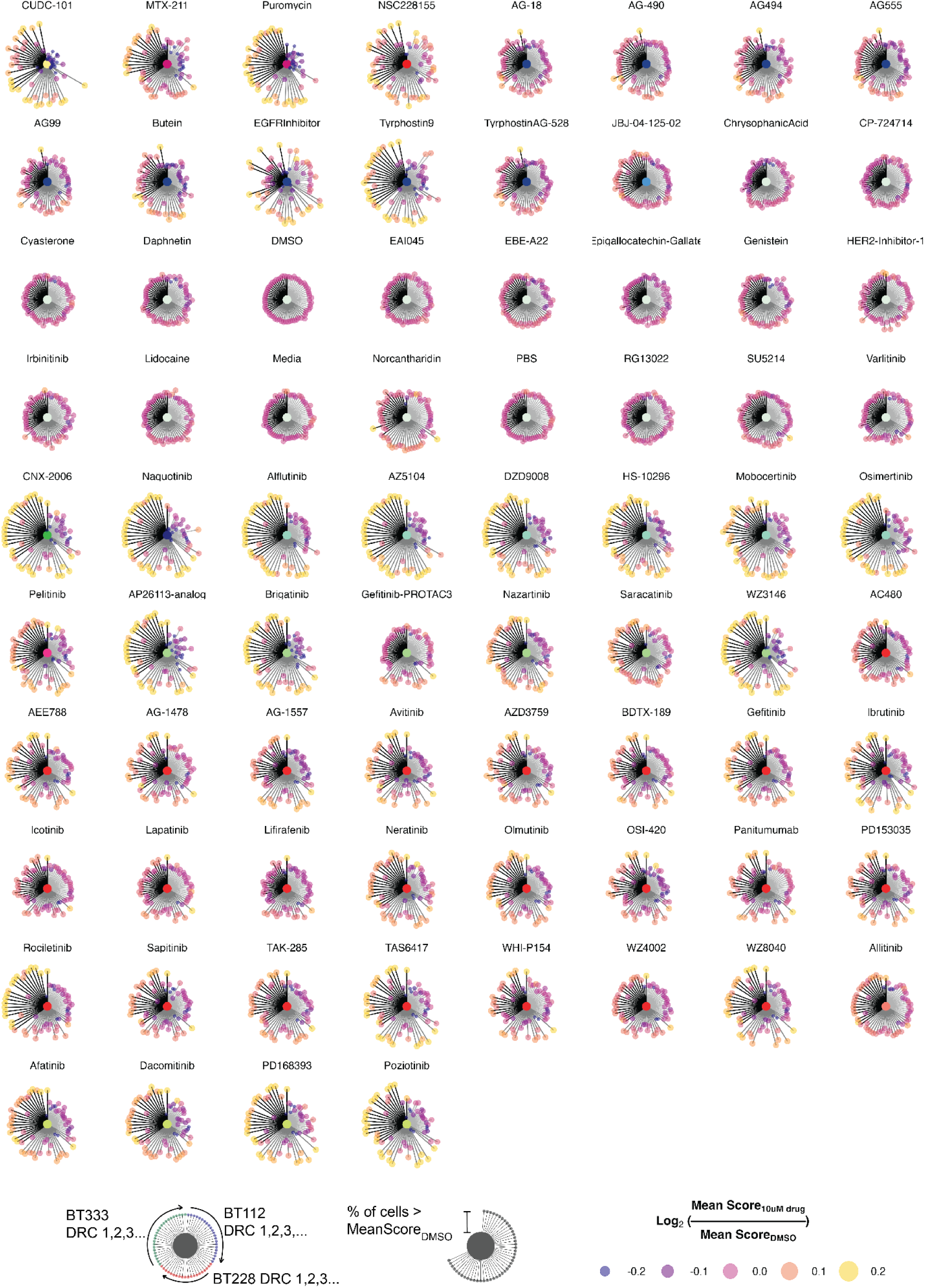
Lollipop visualization of EGFRi response module expression. Briefly, for a given drug, each response module’s respective up-regulated DE genes (identified by covariate-specific DE analysis) were aggregated, averaged, and log2 normalized to the mean of its corresponding DMSO control. This quantity is represented by the size and color of each lollipop. Additionally, because aggregate score is calculated at the single-cell level, heterogeneity in response module expression could be characterized as percent cells within a drug’s high dose population with greater expression than DMSO control. This quantity is represented by the length of each lollipop. The center color of each plot represents its TDC membership.

**Supplemental Figure 22.**
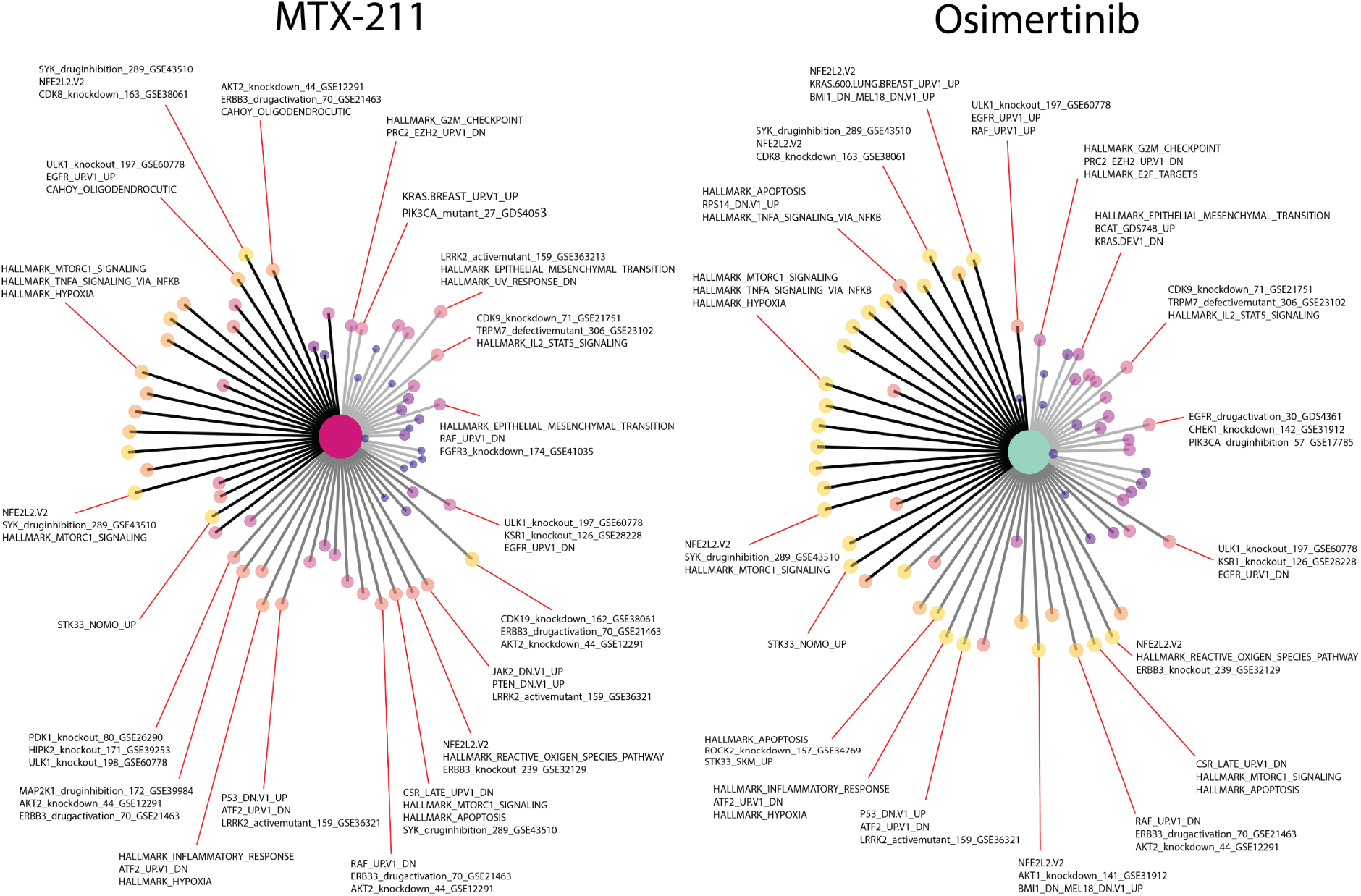
Lollipop plot examples labeled with representative gene sets. The quantities that determine the size and color of each lollipop are described in **Supp. Fig. 21**. GSEA enrichment analysis was performed with the up-regulated DE genes for each response module across PDCLs. Each lollipop is annotated with one to three of the most significantly enriched gene sets in the appropriate response module.

**Supplemental Figure 23.**
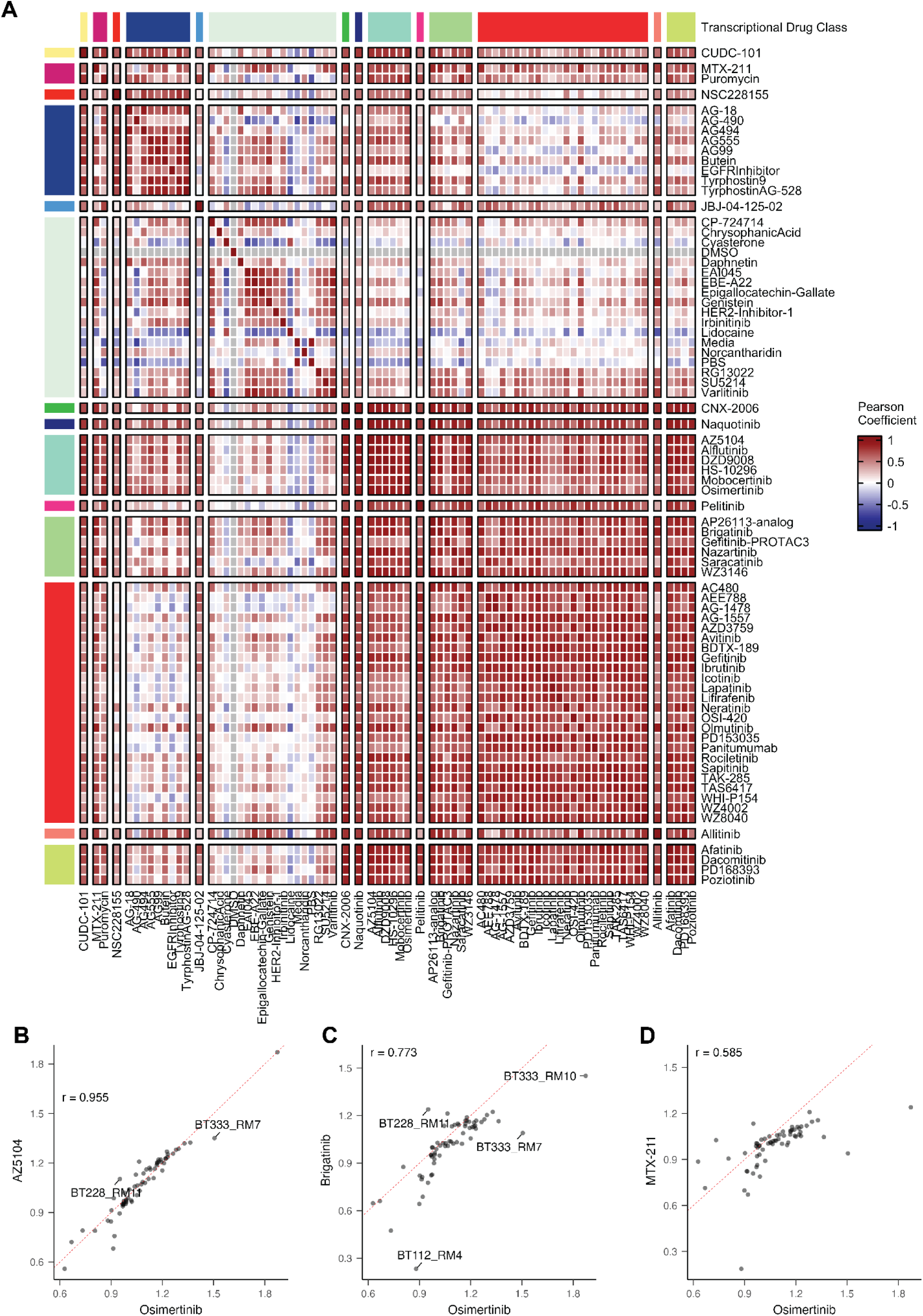
EGFRi within TDCs are more similar than drugs in other TDCs by response module expression. **A.** Pairwise Pearson correlation of a 10uM drug’s response module aggregate expression. **B-D.** Osimertinib RM expression is highly correlated with **(B)** AZ5104, a fellow member of TDC9. Osimertinib RM expression is less correlated with **(C)** Brigatinib and even less so with **(D)** MTX-211, which are part of TDC11 and TDC2, respectively.

**Supplemental Figure 24.**
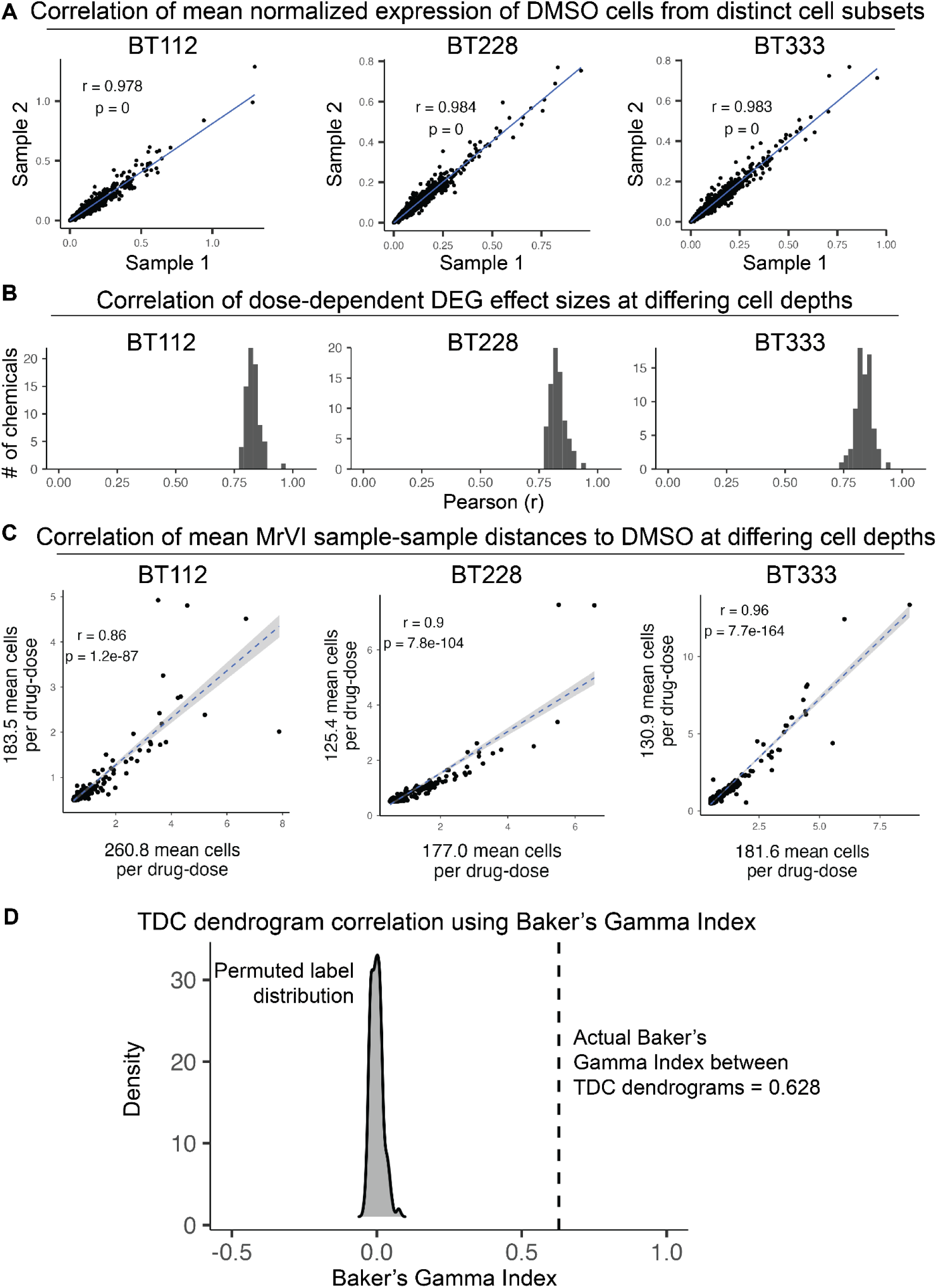
sci-Plex EGFRi data robustness to cell downsampling. **A**. Correlation of mean normalized gene expression of DMSO-treated cells from two exclusive cell samples. Pearson correlation (r) coefficient and p-value are displayed. **B**. Distribution of Pearson correlation (r) coefficients of dose-related beta coefficients derived from experiments of differing cell depths (see axes of C). **C**. Correlation of MrVI-derived mean sample-sample (drug-dose) distances between experiments of differing cell depths. **D.** Similarity of hierarchical hierarchical clusters of TDCs derived at two different cell depths using Baker’s Gamma Index. A distribution of random Baker’s Gamma Indices was generated over 100 trials by calculating the index after permuting labels of one dendrogram. The actual Baker’s Gamma Index between the TDCs is 0.628, where an index of 1 represents completely correlated dendrograms and 0 represents completely uncorrelated dendrograms.

**Supplemental Figure 25.**
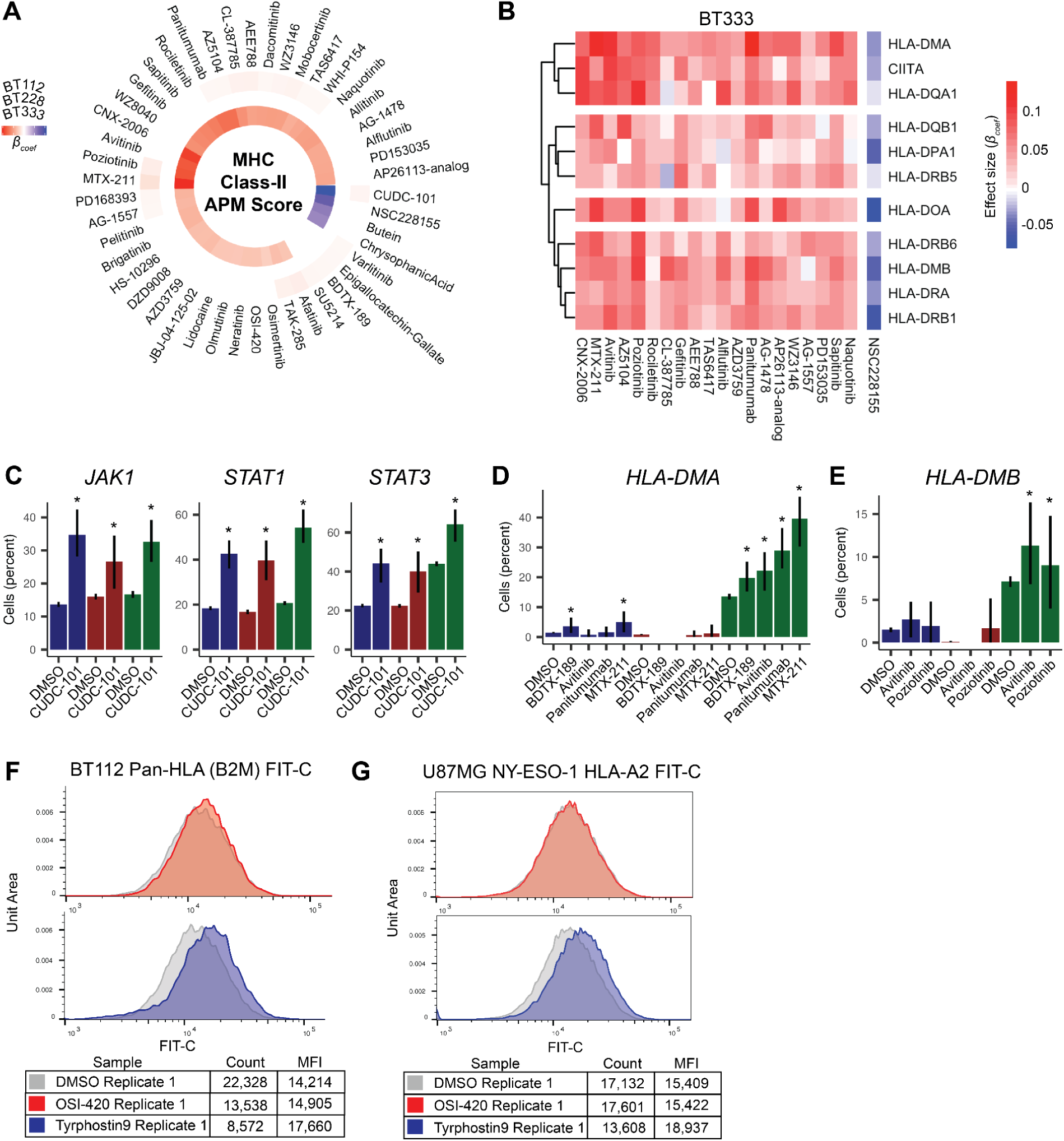
A subset of EGFRis modulate immunogenicity. **A**. Circos heatmaps of beta coefficients for the dose term of a linear regression model of the effect of EGFR inhibition on the expression of genes associated with the MHC class II antigen processing and presentation machinery (APM) in one or more PDCLs. Only significant coefficients (FDR < 5%, Wald test) are shown. **B**. Heatmap of the effect of EGFR inhibition on genes associated with the MHC-CII APM for a subset of EGFRi’s that significantly increase aggregated MHC-CII APM expression in BT333 from (**A**). Note that EGFR activation using NSC228155 leads to decreases in MHC-CII APM. **C**. Percent of cells expressing *JAK1*, *STAT1* or *STAT3* after exposure to 10 μM CUDC-101 or DMSO vehicle control. Colors denote PDCL models: navy = BT112, dark red = BT228, dark green = BT333. Asterisks denote exposures that lead to a significant change in exposure relative to control (FDR < 5%). **D-E**. Percent of cells expressing *HLA-DMA* (**D**) or *HLA-DMB* (**E**) after exposure to the top dose of the specified agent. Asterisks denote exposures that lead to a significant change in exposure relative to control (FDR < 5%). Colors as in (**C**). **F-G**. Representative FIT-C fluorescence density plots of BT112 B2M (pan-HLA) (**F**) protein expression 48hrs post-exposure to 1uM OSI-420 (top) or tyrphostin9 (bottom) exposure. Representative FIT-C fluorescence density plots of U87MG NY-ESO-1 HLA-A*02:01 (**G**) protein expression 48hrs post-exposure to 1uM OSI-420 (*top*) or tyrphostin9 (*bottom*) exposure.

**Supplemental Figure 26.**
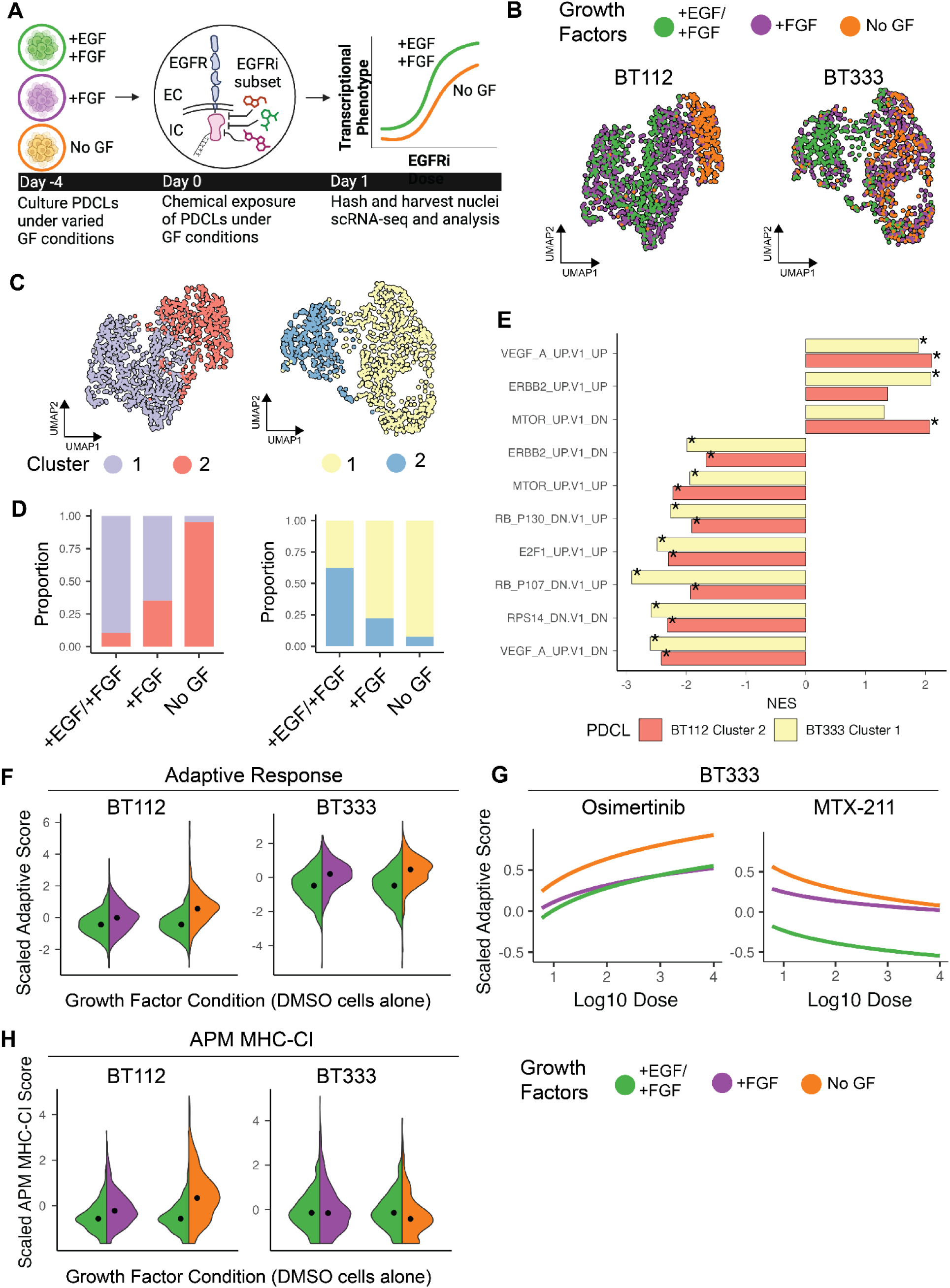
Patient-derived cell line expression changes upon exogenous growth factor removal but consistent transcriptional phenotypic responses to inhibitors. **A.** UMAP of BT112 (*left*) and BT333 (*right*) DMSO-treated cells colored by growth-factor condition. **B**. UMAP of BT112 (*left*) and BT333 (*right*) DMSO-treated cells colored by Leiden cluster. **C**. Proportion of cells within each cluster for each growth factor condition, BT112 (*left*) and BT333 (*right*). **D**. Gene set enrichment for BT112 and BT333 NoGF clusters. A positive normalized effect score (NES) indicates upregulation for the designated cluster (NoGF dominant). Asterisks indicate significance (FDR < 0.05).

**Supplemental Figure 27.**
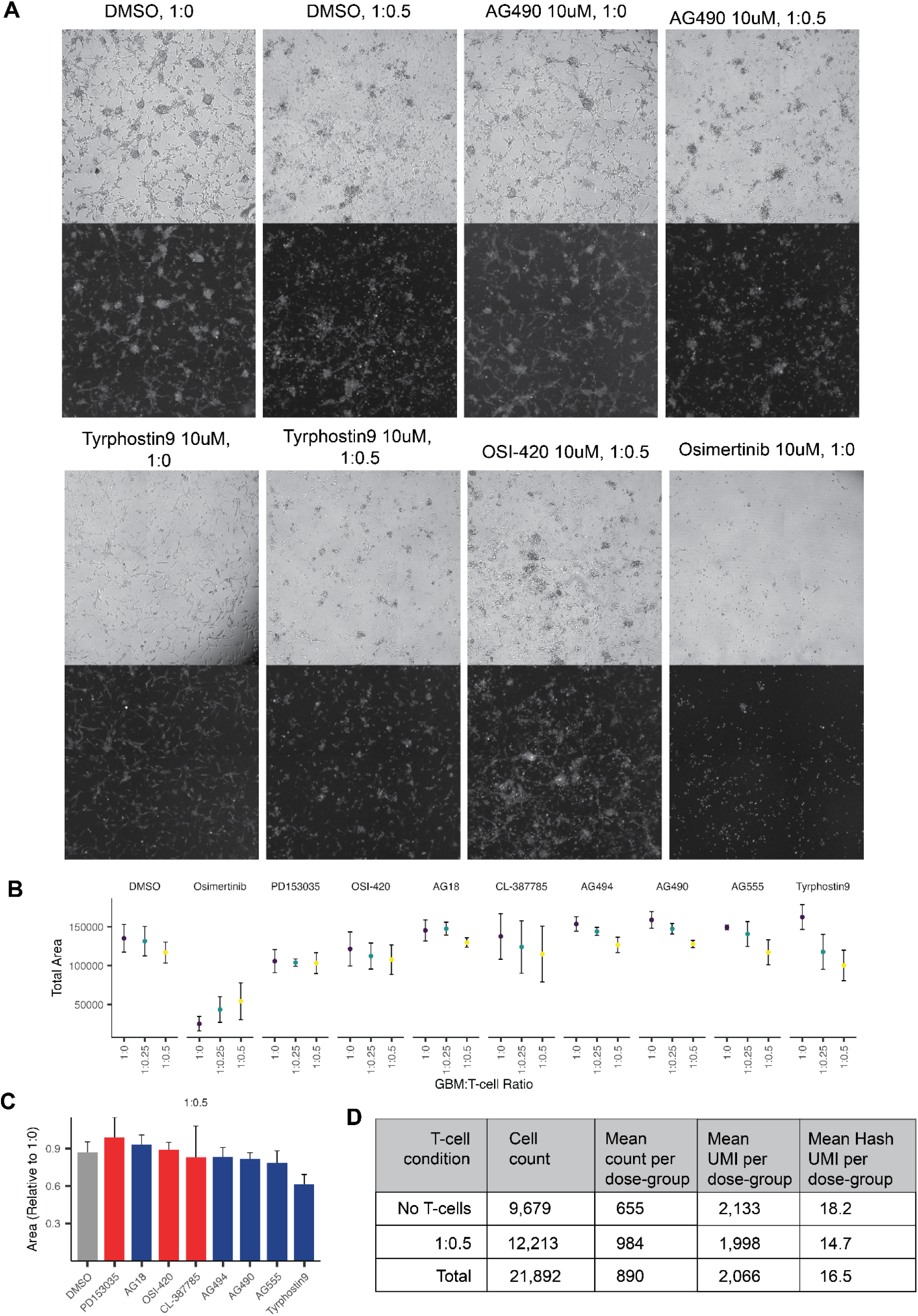
Assessing GBM cytotoxic T-cell killing after pre-exposure to select EGFRi. **A.** Representative U87MG NY-ESO-1 brightfield (top) and fluorescent 488nm (bottom) 48hr images for select EGFRi pre-treatment and GBM:T-cell ratios, denoted as titles. Osimertinib images were included to demonstrate its drastic effect on U87MG cells even absent of CD8^+^ T-cells. **B.** Raw measurements of occupied CFSE area for 10uM EGFRi pre-treated U87MG NY-ESO-1 at 48hrs post-T-cell addition. It is important to note that osimertinib high dose treatment maximally decreased U87MG CFSE area and that addition of T-cells did not lead to any further decrease. **C.** Quantifications as in **Fig. 5F** for cells exposed to the specified inhibitor and a 1:0.5 ratio of GBM:T-cells. The color of the bars denote TDC membership, and the inhibitors in J are ordered by DMSO then increasing T-cell effect, as decreasing mean relative area. Error bars represent standard deviation from the mean across replicates (n_DMSO,T-cell_ = 34, n_drug,T-cell_ = 11, n_Tyrphostin9,T-cell_ = 9). **D.** Table of experimental summary metrics for the engineered BT333:T-cell culture experiment to assess tyrphostin9-induced APM effects.

## Supplementary Tables

**Supplementary Table 1: Pilot screen: Differentially expressed genes as a function of dose for each EGFR inhibitor.**

**Supplementary Table 2: Basal PDCL: Differentially expressed genes as a function of PDCL cluster.**

**Supplementary Table 3: Chemicals used in comprehensive EGFR inhibitor screen.**

**Supplementary Table 4: Comprehensive screen: Differentially expressed genes as a function of dose for each EGFR inhibitor.**

**Supplementary Table 5: Comprehensive screen: Significant differentially expressed genes as a function of MrVI-determined response module.**

**Supplementary Table 6: Membership of each drug within each Transcriptional Drug Class (TDC).**

